# The characteristics of inducible defenses influence predator-prey dynamics

**DOI:** 10.1101/2023.08.24.554708

**Authors:** Michael H. Cortez, Emily Mila, Edward Hammill

## Abstract

Empirical and theoretical studies suggest inducible and evolving defenses have different effects on predator-prey dynamics. However, theory for inducible defenses does not account for differences in response stimuli, reversibility, and within-generation versus transgenerational responses. We use predator-prey models to explore how these characteristics influence the effects of inducible defenses on stability and predator-prey oscillations. We find that while inducible defenses are typically stabilizing, defenses responding to any stimuli (predator density, conspecific and predator densities, predation rates, or the fitness gradient) can be destabilizing. Also, while induced defenses typically shorten predator-prey phase lags, defenses that respond to prey densities or the fitness gradient can increase the lags. Our theory helps explain why inducible defenses are less likely to be destabilizing and increase phase lags than evolving defenses: inducible defenses typically have negative feedbacks of the trait on its own dynamics whereas evolving defenses under disruptive selection can have positive feedbacks.

## 1 Introduction

Intraspecific variation can shape the dynamics and composition of ecological communities (Bolker et al., 2003; Miner et al., 2005; Agrawal et al., 2007; Kishida et al., 2010; Bolnick et al., 2011; Forsman, 2015). In predator-prey systems, intraspecific variation in prey defense can be due to genetic variation or inducible defenses, wherein plastic changes in individual behavior, life history, or morphology reduce exploitation by predators (Tollrian and Harvell, 1999; Matz and Kjelleberg, 2005; Van Donk et al., 2011). Evolving and inducible defenses have the potential to drive population-level changes in defense that are as fast as changes in population densities (Grosklos and Cortez, 2021), and the coupling of density dynamics with inducible defenses (Verschoor et al., 2004; Lurling et al., 2005; van der Stap et al., 2006, 2009) or rapidly evolving defenses (Yoshida et al., 2003; Becks et al., 2010; Hiltunen et al., 2014a; Yamamichi et al., 2015) can alter predator-prey dynamics. This study explores how the characteristics of inducible defenses influence their effects on predator-prey dynamics, and explains why inducible and evolving defenses have different population-level effects.

Empirical studies suggest inducible and evolving defenses can affect predator-prey dynamics in different ways. Eco-evolutionary studies have shown that evolving defenses can be stabilizing or destabilizing (Yoshida et al., 2004; Becks et al., 2010; Kasada et al., 2014) and increase the phase-lags in predator-prey oscillations (Yoshida et al., 2003, 2007; Hiltunen et al., 2014b). In contrast, inducible defense studies only report stabilizing effects (Kusch, 1993; Verschoor et al., 2004; Lurling et al., 2005; van der Stap et al., 2006, 2009; Boeing and Ramcharan, 2010) and increased phase lags have not been observed. These contrasting effects are supported by theory predicting that evolving defenses can be stabilizing or destabilizing and shorten or lengthen phase lags (Abrams and Matsuda, 1997; Jones et al., 2009; Cortez and Ellner, 2010; Cortez, 2016) whereas inducible defense are stabilizing and shorten lags (Abrams and Walters, 1996; Vos et al., 2004; Ramos-Jiliberto, 2003; Mougi and Kishida, 2009; Cortez, 2011; Yamamichi et al., 2011). Altogether, this body of work suggests inducible defenses have a narrower set of effects on predator-prey dynamics than evolving defenses.

However, this conclusion is likely premature because we have a limited understanding of how mechanistic differences between inducible defenses influence their effects on population dynamics (Bolker et al., 2003; Miner et al., 2005; Kishida et al., 2010). In particular, induction is commonly stimulated by predator chemical cues (Tollrian and Harvell, 1999; Aŕanguiz-Acuña et al., 2010; Auld and Relyea, 2011; Belovsky et al., 2011), but induction levels also can respond to resource availability (Anholt et al., 1996; Van Donk et al., 1999), conspecific density (Relyea, 2004; Teplitsky and Laurila, 2007; Van Buskirk et al., 2011; Tollrian et al., 2015), and predation cues such as diet-dependent predator karimones and conspecific alarm cues (Buskirk and Arioli, 2002; Schoeppner and Relyea, 2009; Van Donk et al., 2011; Gu et al., 2020, 2023). In addition, inducible defenses can be reversible or irreversible (Tollrian and Harvell (1999) and references within) and response timing can be within an individual’s lifetime, with induction times ranging from hours to weeks (Kuhlmann and Heckmann, 1985; Schoeppner and Relyea, 2009; Relyea and Auld, 2004; Auld and Relyea, 2011), or transgenerational (i.e., maternally induced defenses; Halbach 1970; Agrawal et al. 1999; Shimada et al. 2010). Current theory has limited ability to predict how these differences scale up to affect population-level dynamics because modeling studies primarily focus on reversible, within-generation inducible defenses driven by predator cues or cues that allow individuals to maximize individual fitness (Ives and Dobson, 1987; Ramos-Jiliberto, 2003; Abrams, 2003; Vos et al., 2004; Mougi and Kishida, 2009; Cortez, 2011; Yamamichi et al., 2011; Kovach-Orr and Fussmann, 2013). Thus, current theory only addresses a fraction of the observed variation in characteristics of inducible defenses.

In this study, we analyze predator-prey models with inducible prey defenses and explore how differences in response stimuli, reversibility, and timing influence the effects of inducible defenses on stability and predator-prey oscillations. We identify when inducible defenses have stabilizing versus destabilizing effects and when they increase and decrease predator-prey phase lags. We then use those conditions to explain why inducible and evolving defense typically have different effects on predator-prey dynamics.

## 2 Models

### 2.1 Predator-prey model with a reversible inducible defense

We start with a predator-prey model where undefended prey have trait value *α*_1_ and defended prey have trait value *α*_2_ (*α*_1_ *< α*_2_), prey can change their phenotype during their lifetime, induction is reversible, and prey offspring initially have the same phenotype as their parent. Individual induction and loss of induction rates are *ɛ_I_P_I_* and *ɛ_L_P_L_*, respectively, where *ɛ_I_* and *ɛ_L_* are the maximum rates and the fractions *P_I_* and *P_L_* define how the rates vary due to environmental cues. The dynamics of the densities of each prey type (*x_i_*) and the predator (*y*) in the dimorphic trait model are

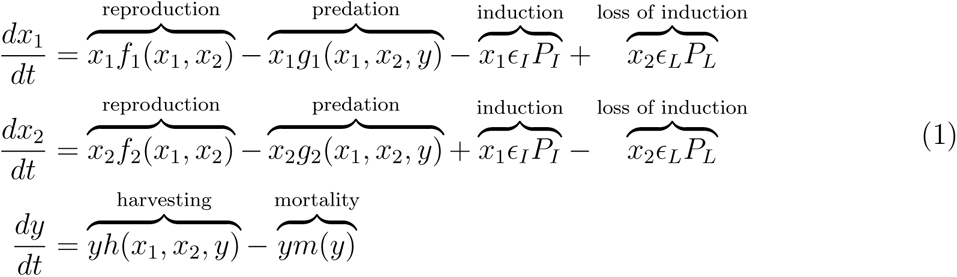

where for individuals with phenotype *α_i_*, *f_i_*(*x*_1_*, x*_2_) is the per capita growth rate in the absence of predation and *g_i_*(*x*_1_*, x*_2_*, y*) is the per capita predation rate; *yh*(*x*_1_*, x*_2_*, y*) is the predator harvesting rate; and *ym*(*y*) is the predator mortality rate of the predator. The notation for all models is summarized in Table 1. We assume the functions satisfy conditions consistent with intraspecific prey competition, a predatory interaction, and intraspecific predator competition or predator interference; see Appendix S1.1. In addition, we assume the prey reproduction and predation rates are decreasing functions of prey defense, which results in a trade-off between predation rates and reproductive output.

**Table 1:**
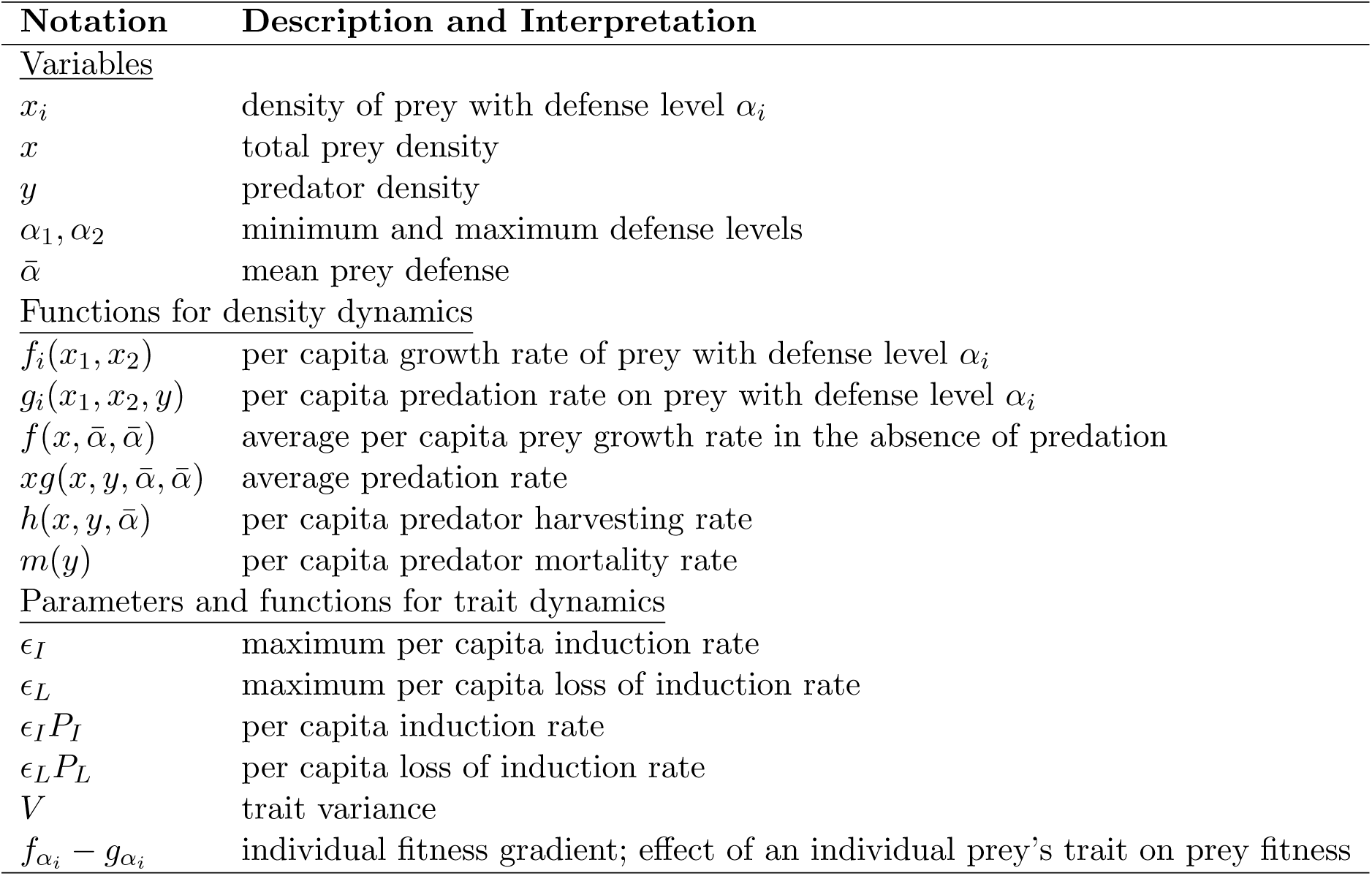
Model variables, parameters, and important quantities.

To facilitate comparisons with other inducible defense models and evolving defense models, we approximate the discrete trait model by a continuous trait model that describes the dynamics of the total prey density, *x* = *x*_1_ + *x*_2_, and mean prey defense, 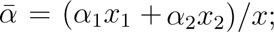 see Appendix S1.1 for details. This allows us to simultaneously analyze continuous and discrete inducible defenses and interpret our results in terms of the average of a continuous trait or fractions of defended and undefended individuals. The continuous trait model is

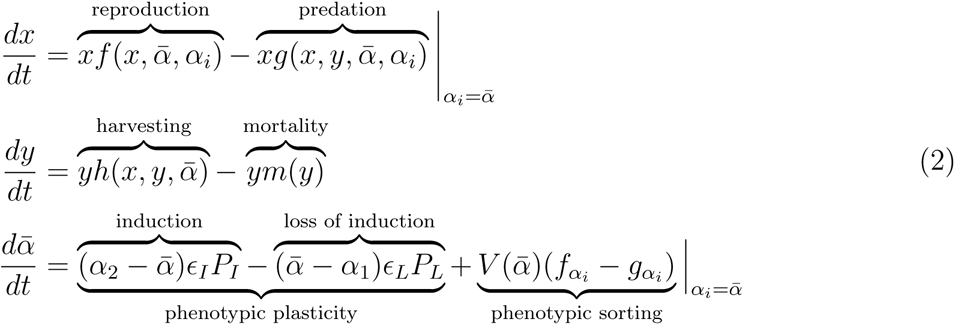

where *f*, *g*, and *h* are the average per capita prey growth, predation, and harvesting rates. In the prey and predator equations, the individual trait value (*α_i_*) is a place-holding variable that allows for prey fitness to be frequency-dependent. The prey and predator equations are evaluated at the mean trait value 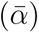 because the prey and predator growth rates are determined by the average defense level.

In the trait equation, mean defense changes due to phenotypic plasticity and phenotypic sorting (*sensu* Yamamichi et al. 2019). The phenotypic plasticity component accounts for trait change caused by induction and loss of induction. The phenotypic sorting component accounts for trait change caused by differential survival and reproduction of the different phenotypes. The direction of change is determined by the individual fitness gradient (*f_α_i__ −g_α_i__*), which drives the mean trait in the direction of increasing fitness. (Throughout, subscript variables denote partial derivatives, e.g., *f_α_i__*= *∂f/∂α_i_*.) The variance in prey defense, 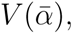 depends on the mean trait because the variance approaches zero when the mean trait approaches *α*_1_ or *α*_2_. The variance is small when all individuals have identical levels of induction and large when, e.g., there is imperfect switching or inconsistent assessment of environmental cues by the prey.

### 2.2 Accounting for inducible defenses that differ in response stimuli, reversibility, and timing

Instead of explicitly modeling the cues affecting induction and loss of induction, we assume induction and loss of induction rates depend on the model state variables. In effect, we assume cues degrade quickly and provide an accurate measure of their source; this omits delays that could potentially be destabilizing (Yamamichi et al., 2019). We analyze four classes of inducible defenses that respond to different sets of cues (see Appendix S1.4 for mathematical details). Class 1 inducible defenses respond to predator density such that higher predator density increases induction rates and decreases loss of induction rates. Class 2 inducible defenses respond to predator and prey (conspecific) densities such that higher predator density and lower prey density increase induction rates and decrease loss of induction rates. Class 3 inducible defenses respond to predation rates (e.g., alarm cues from conspecifics) such that higher predation rates increase induction rates and decrease loss of induction rates. Class 4 inducible defenses assume individuals respond to a comprehensive set of cues that allows individuals to maximize per capita fitness. Mathematically, we assume induction and loss of induction rates depend on the fitness gradient (*f_α_i__ − g_α_i__*) such that induction rates increase and loss of induction decrease whenever more defended prey have higher fitness.

Many prior studies have analyzed specific forms of models (1) and (2) with Class 1 (Vos et al., 2004; Ramos-Jiliberto, 2003; Cortez, 2011; Yamamichi et al., 2011; Kovach-Orr and Fussmann, 2013) or Class 4 (Ives and Dobson, 1987; Abrams, 2003) inducible defenses where induction and loss of induction have equal maximum rates (*ɛ_I_* = *ɛ_L_*) and equal thresholds (resulting in *P_I_* + *P_L_* = 1). Our study extends and generalizes this prior work by allowing for other response stimuli and unequal maximum rates (*ɛ_I_* ≠ *ɛ_L_*) and unequal thresholds (*P_I_* + *P_L_* ≠ 1) for induction and loss of induction. Note that we do not focus on the latter two because they do not qualitatively alter our predictions. Fitness gradient models and optimal trait models are two other kinds of inducible defense models in the literature (reviewed in Yamamichi et al. (2019)). Our results are likely to apply to fitness gradient and optimal trait models, because the dynamics of those models are often qualitatively similar to the dynamics of the continuous trait model (2) with Class 4 inducible defenses.

We study irreversible and transgenerational inducible defenses using two additional continuous trait models that can be derived from discrete trait models (Box 1). The second model is identical to the first except that all offspring are initially undefended; that model allows us to analyze irreversible defenses by setting the loss of induction rate to zero (*ɛ_L_P_L_* = 0). The third model assumes transgenerational responses, meaning an individual’s phenotype is determined at birth. Importantly, the continuous trait versions of all three models have similar structures (Box 1): the prey and predator equations are identical, the phenotypic plasticity terms of the trait equations are nearly identical, and only the phenotypic sorting terms of the trait equations differ because of the different assumptions about the reversibility and timing of induction. This similarity allows us to identify how differences in reversibility and timing influence the effects of inducible defenses on predator-prey dynamics.

#### Box 1: Structural similarities between alternative continuous trait inducible defense models

The structure of the continuous trait model (1) is similar to the structures of two continuous trait models we use to approximate the dynamics of discrete trait models with irreversible or transgenerational inducible defenses. Here, we briefly introduce the discrete and continuous trait versions of the other two models and refer the reader to Appendices S1.2 and S1.3 for additional details. In the following, we do not show the predator equations because they are identical to the predator equations in models (1) and (2).

To study irreversible defenses, we start with a discrete trait model that is identical to model (1) except that all offspring are initially undefended,

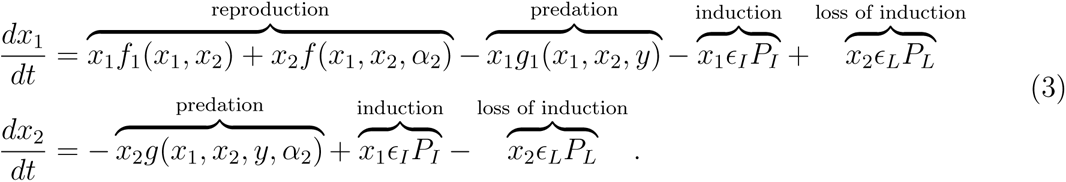

All terms are interpreted as in the dimorphic trait model (1). We approximate the discrete trait model (3) with a continuous trait model of the total prey density, *x* = *x*_1_ + *x*_2_, and mean prey trait, 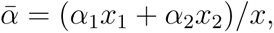

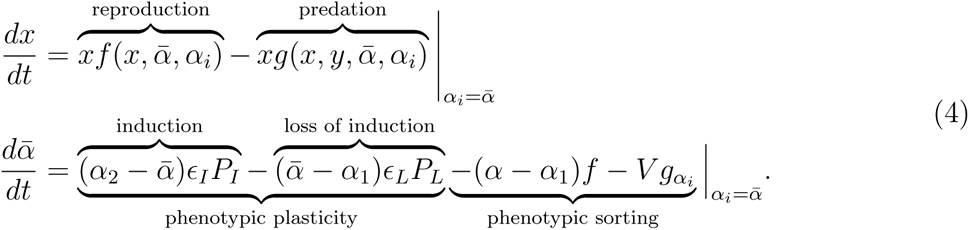

The phenotypic plasticity terms of the trait equation are identical to those in the continuous trait model (2). The phenotypic sorting terms accounts for differing predation rates on the different prey phenotypes and how the mean trait decreases when offspring are produced. Note that if the maximum rates of induction and loss of induction are large (*ɛ_I_* and *ɛ_L_* large), then the phenotypic sorting terms are negligibly small and the trait dynamics are determined by just the phenotypic plasticity terms.

Importantly, the inducible defense becomes irreversible in models (3) and (4) when loss of induction is not possible (*ɛ_L_P_L_* = 0). Thus, we can study the effects of irreversibility and loss of induction being slower than induction by decreasing the maximum rate of loss of induction (*ɛ_L_*) to zero.

To study transgenerational inducible defenses, we use a discrete trait model where an individual’s trait is determined at birth or hatching. We assume all parents produce the same fractions of induced (*P_I_*) and uninduced (*P_L_*) offspring and those fractions depend on environmental stimuli. The equations for undefended prey (*x*_1_) and defended prey (*x*_2_) are

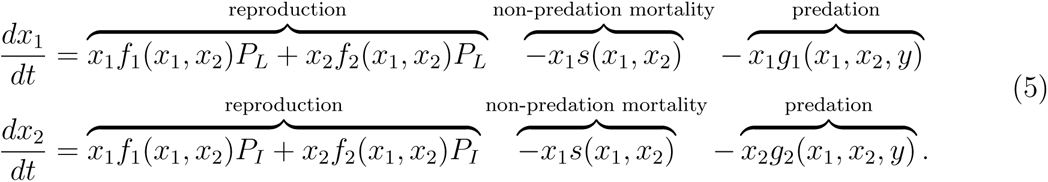

In the reproduction terms, *x_i_f_i_*(*x*_1_*, x*_2_)*P_I_* and *x_i_f_i_*(*x*_1_*, x*_2_)*P_L_* are the rates at which prey *i* produces induced and uninduced offspring. The prey reproduction (*f_i_*) and non-predation mortality (*s*) rates are combined into one term in the other models, but we keep them separate in this model in order to avoid confusion about the effects of plasticity. We approximate the discrete trait model (5) with the continuous trait model of the total prey density (*x*) and mean prey trait 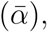

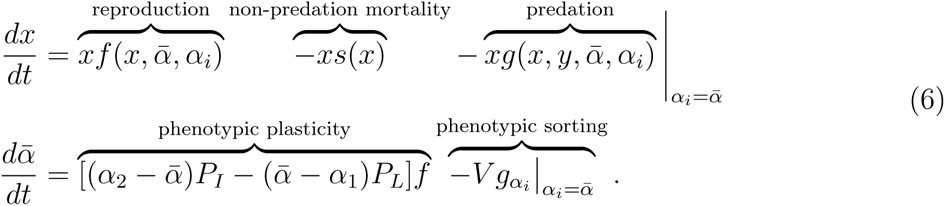

In the trait equation, the phenotypic plasticity term accounts for trait change due to the production of induced and uninduced offspring. The phenotypic sorting terms account for differing predation rates on the prey phenotypes. Note that if there is rapid turnover the prey population, meaning high reproduction rates (*f* large) and high non-predation mortality rates (*s* large), then the phenotypic sorting terms are negligibly small and the trait dynamics are effectively determined by just the phenotypic plasticity terms; see Appendix S1.3 for mathematical details.

The continuous trait versions of the three models have similar structures. First, the prey and predator density equations are identical. Second, the phenotypic plasticity terms of the trait equations are nearly identical and have qualitatively similar dynamics because the prey reproduction rate *f* in model (6) plays a role analogous to the maximum rates *ɛ_I_* and *ɛ_L_* in models (2) and (4). Third, all models account for phenotypic sorting, but the assumptions about the reversibility and timing of responses result in different phenotypic sorting terms.

This similarity in structure allows us to identify how response stimuli, irreversibility, and timing affect predator-prey dynamics. First, because response stimuli only affect the phenotypic plasticity component of trait change and the phenotypic plasticity components of the models are effectively identical, the effects of response stimuli are the same for all models. Moreover, when rates of phenotypic plasticity are fast, the effects of inducible defenses are determined by just the phenotypic plasticity terms. Second, the effects of irreversibility are determined by the effects of decreasing *ɛ_L_* to zero is model (4). Third, the effects of transgenerational responses are determined by the consequences of the phenotypic plasticity terms in model (6) depending on the prey reproduction rate. In total, the similar structures of the continuous trait models facilitate the identification of how differences in the characteristics of inducible defenses influence the population-level dynamics of predator-prey systems.

The forms of the continuous trait versions of the models also facilitate comparisons of the effects of inducible and evolving defenses. Specifically, in the limit where there is no induction or loss of induction (*ɛ_I_* = *ɛ_L_* = 0), the trait equation in model (2) is identical to the trait equation used in quantitative genetics models of evolving defenses (Abrams et al., 1993; Cortez, 2016). Thus, any differences between the effects of inducible defenses in model (2) and evolving defenses must be due to the phenotypic plasticity component of trait change. This study does not present an analysis of the evolving defense model (see Cortez 2016), but the effects of evolving defenses are almost identical to the effects of Class 4 inducible defenses in model (2) because both maximize individual prey fitness.

## 3 Results

### 3.1 Determining when mechanistic differences matter

Regardless of the reversibility and timing of induction, the effects of response stimuli are the same for all inducible defenses. Moreover, when rates of phenotypic plasticity are fast, the effects of the inducible defense are entirely determined by the response stimuli. Thus, reversible and irreversible intragenerational responses and transgenerational responses have identical effects on equilibrium stability and predator-prey phase lags when rates of phenotypic plasticity are sufficiently fast. Biologically, rates of phenotypic plasticity are fast for intragenerational responses when rates of induction and loss of induction are large (*ɛ_I_*, *ɛ_L_*) and for transgenerational responses when there is rapid turnover in the prey population (Box 1). Mathematically, this is because the stimuli only affect the phenotypic plasticity component of trait change, the phenotypic plasticity components of the models are effectively identical, and the effects of the phenotypic plasticity components are much larger than all other effects when rates of phenotypic plasticity are large (Box 1).

When rates of phenotypic plasticity are not as rapid, the effects of phenotypic sorting, reversibility, and timing of induction are non-negligible and potentially can alter how inducible defenses affect predator-prey dynamics. We predict that the effects of phenotypic sorting are negligible for most systems. This is because, in numerical simulations, the effects of phenotypic sorting were almost always negligibly small relative to the effects of phenotypic plasticity. The only exception was when rates of phenotypic plasticity were much slower than the prey reproduction and predation rates. However, this scenario does not match the rates of induction and loss of induction in the empirical studies cited in the Introduction.

As as consequence of the above, the following sections first explore the effects of response stimuli on equilibrium stability and predator-prey phase lags and then address how irreversibility and timing alter the effects. For completeness, the effects of phenotypic sorting are addressed in Appendices S1.5 and S1.6.

### 3.2 Effects of inducible defenses on system stability

We explore when inducible defenses stabilize or destabilize equilibria using the Routh-Hurwitz criteria for equilibrium stability (see Appendix S1.5). Here, stabilization means inducible defenses dampen or prevent oscillations that would occur in the absence of induction and destabilization means inducible defenses cause cycles around an equilibrium that would be stable in the absence of induction.

Inducible defenses are typically stabilizing when the response stimuli are predator density, conspecific and predator density, or predation rates (Classes 1-3, respectively). However, destabilization can occur in two scenarios. First, for Classes 1-3, destabilization can occur when increased mean defense greatly decreases prey fitness (e.g., the prey defense is aggression and increased average aggression reduces prey reproduction rates because of strong negative intraspecific prey interactions; 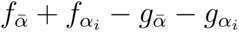 negative and large in magnitude). Under this condition, the destabilizing negative feedback is increased predator density leads to increased mean defense, increased mean defense reduces prey density due to reduced prey fitness, and reduced prey density leads to lower predator density. Second, for Class 3, destabilization can occur when increased mean defense greatly increases prey fitness (e.g., prey have a shared defense and greater mean defense reduces predation to all prey; 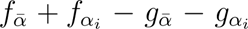 positive and large). Here, the destabilizing positive feedback is increased prey density causes higher predation rates, higher predation rates lead to higher levels of defense, and higher mean defense increases prey densities because it increases prey fitness. Overall, this produces a positive feedback where an initial increase in prey density leads to greater increases in prey density.

Inducible defenses responding to the fitness gradient (Class 4) are often stabilizing, but they can be destabilizing under two scenarios. First, the most likely scenario is when prey fitness is maximized at extreme trait values (i.e., maximal or minimal levels of defense are optimal and intermediate levels of defense are suboptimal; 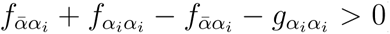). Under this condition, the destabilizing positive feedback is that small increases in mean defense lead to greater increases in defense. Second, destabilization can also occur when fitness is maximized at intermediate trait values (i.e., intermediate levels of defense are always optimal; 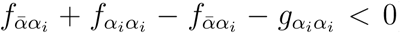) if increased mean defense decreases prey fitness (e.g., when the prey defense is increased aggression; 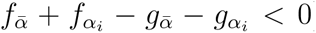) and the benefits of higher defense decrease at higher prey density (*f_xα_i__ − g_xα_i__ <* 0). Here, the destabilizing positive feedback is reduced prey density leads to higher induction rates and defense levels, and increased mean defense reduces prey density due to reduced prey fitness. Overall, this produces a positive feedback where an initial decrease in prey density leads to greater decreases in prey density. Note that the conditions for Class 4 inducible defenses are identical to the conditions for an evolving defense to be destabilizing (Cortez, 2016).

The (de)stabilizing effects listed above only change an equilibrium from unstable to stable, or vice versa, when their effects are sufficiently large. For intragenerational inducible defenses this means induction and loss of induction are sufficiently fast (*ɛ_I_, ɛ_L_* large) and for transgenerational responses this means prey reproduction rates are sufficiently high (*f* large). Figure 1 illustrates this for intragenerational inducible defenses. Stabilization requires either intermediate rates (Classes 2-4; not shown) or sufficiently fast rates (Classes 1-4; Figure 1A-C) of induction and loss of induction rates. Destabilization requires either intermediate rates (Classes 1-4; Figure 1D-F) or sufficiently fast rates (Classes 2-4; Figure 1G-I) of induction and loss of induction.

**Figure 1:**
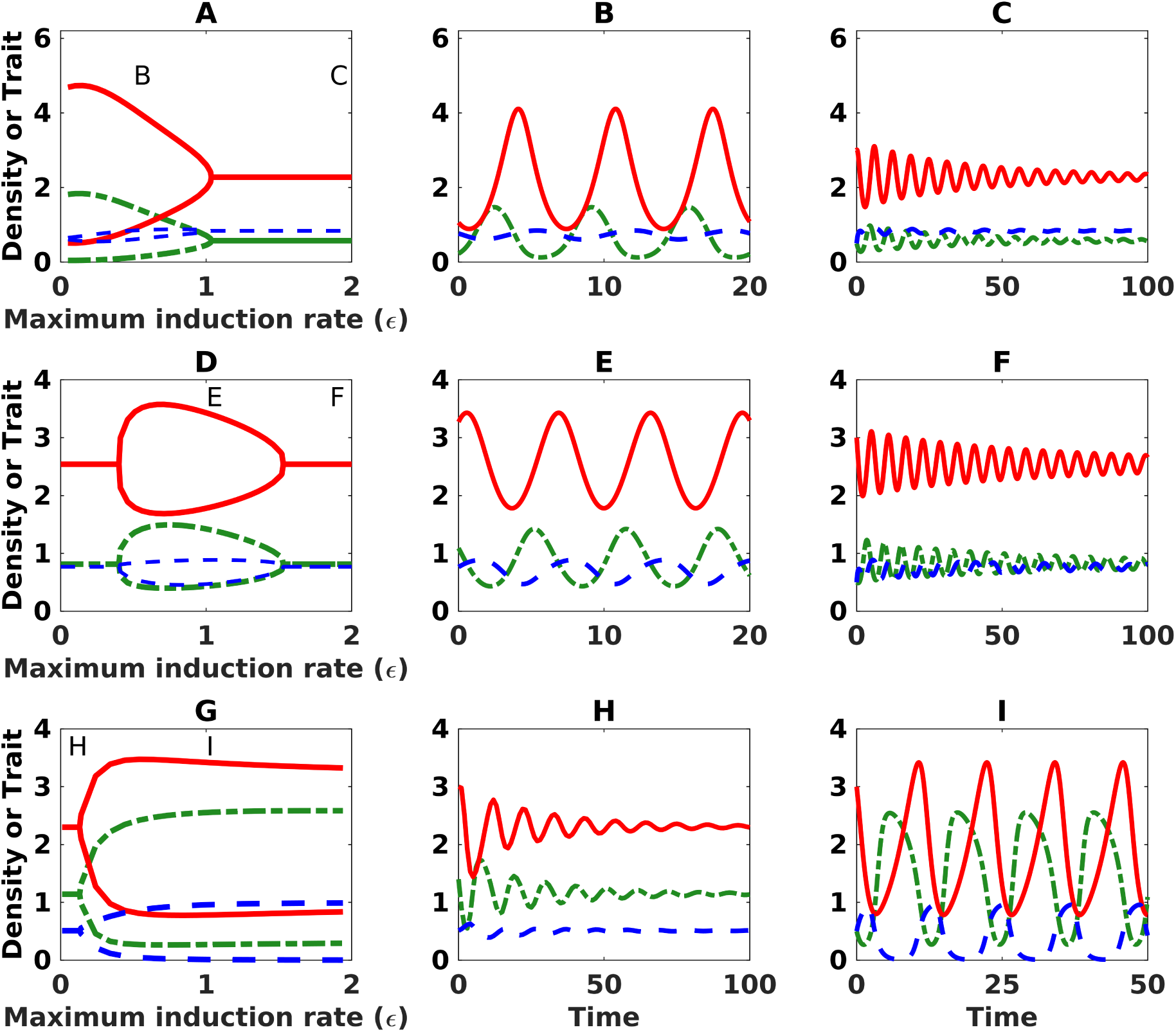
Inducible defenses can (A) stabilize oscillating systems when responses are sufficiently fast (Classes 1-4), (B) destabilize stable systems when responses are intermediate (Classes 1-4), or (C) destabilize stable systems when responses are sufficiently fast (Classes 2-4). In all panels, the maximum rates of induction and loss of induction are equal (*ɛ_I_* = *ɛ_L_* = *ɛ*). Prey density is dashed-dot green, predator density is solid red, and mean defense is dashed blue. Column 1 shows minimum and maximum asymptotic values; a single curve for a given induction rate indicates the value at a stable equilibrium and two curves indicate the minimum and maximum values during sustained oscillations. Note that in the absence of induction (i.e., for fixed defenses), the system is unstable for A and stable for D and G. Columns 2 and 3 show time series corresponding to letters in the panels of column 1. See Appendix S2 for parameter values.

We now address the effects of irreversibility and transgenerational responses. Irreversibility can have a stabilizing (Figure S1A-F) or destabilizing (Figure S2G-I) effect (see Appendix S1.5.7 for mathematical details). The stabilizing effect is caused by irreversibility leading to higher levels of mean defense (because loss of induction is not possible), which lowers prey growth rates and predation rates, both of which have stabilizing effects on the dynamics of the prey and predator densities. The destabilizing effect arises because the absence of loss of induction removes a stabilizing feedback of the trait on its own dynamics. We predict that irreversible defenses are more stabilizing than reversible defenses when increased defense causes large decreases in prey growth rates and predation rates (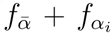 and 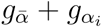 large in magnitude) and more destabilizing when changes the decreases are not as large. We note that the effects of loss of induction being slower than induction (*ɛ_L_ < ɛ_I_*), e.g., because individuals remain induced after a cue is removed (Sih, 1992; Fraker, 2008), are the same as the effects of irreversibility. Mathematically, this is because irreversibility is the limit where loss of induction rates approach zero.

Transgenerational inducible defenses can be more stabilizing or destabilizing than intragenerational inducible defenses (see Appendix S1.5.8), however in simulations transgenerational responses were almost always more destabilizing. Thus, we predict transgenerational inducible defenses are less stabilizing and more destabilizing than intragenerational inducible defenses. The destabilizing effect is larger when increased defense causes large decreases in prey growth rates (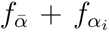 large in magnitude). It is caused by a delay wherein increased defense reduces prey reproduction rates, and reduced reproduction rates lower the rate of change for the mean trait.

### 3.3 Effects of inducible defenses on predator-prey phase lags

We explore how inducible defenses influence predator-prey phase lags by computing the phase lags at Hopf bifurcations, i.e., the parameter values where equilibria transition from stable to unstable; see Appendix S1.6 for mathematical details. This approach is guaranteed to yield accurate predictions about phase lags for parameter values near the Hopf bifurcations and when the amplitudes of the oscillations are not large. However, it may yield inaccurate predictions for large amplitude oscillations in other regions of parameter space (Ellner and Becks, 2011; Cortez, 2018).

Predator oscillations lag behind prey oscillations by a quarter-period or less for fixed defenses (Bulmer, 1975). Inducible defenses of all types typically shorten the lag. However, there are two general mechanisms through which inducible defenses can increase the lag: (i) positive feedbacks of the trait on its own dynamics 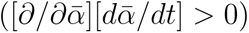 and (ii) increased prey density leading to lower levels of defense 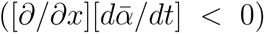. Both mechanisms delay the trough in mean prey defense, which allows predator density to increase for longer and delays the peak in predator density. We expect mechanism (i) to be more common in nature because mathematically it increases the phase lag in a larger region of parameter space than mechanism (ii).

Response stimuli influence the range of possible phase lags. First, the phase lags are shortened whenever inducible defenses respond to predator density (Class 1) or predation rates (Class 3); see Figure 2A,D for an illustration. Second, the phase lags are typically shortened by inducible defenses responding to prey and predator densities (Class 2) or the fitness gradient (Class 4). However, Class 2 inducible defenses can lengthen phase lags via mechanism (ii) and cause lags grater than a half-period when induction and loss of induction rates are sufficiently fast (*ɛ_L_, ɛ_I_* large) and highly sensitive to changes in prey density (*∂P_I_/∂x*, *∂P_L_/∂x* large in magnitude; Figure 2B,E). Class 4 inducible defenses can increase phase lags via mechanism (i) when prey fitness is maximized at extreme trait values (i.e., maximal or minimal levels of defense are always optimal and intermediate defense levels are always suboptimal; 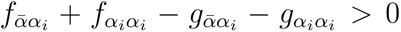) (Figure 2C,E) and via mechanism (ii) when the benefits of higher defense decrease at higher prey density (*f_xα_i__ − g_xα_i__ <* 0). Thus, we predict that inducible defenses typically shorten predator-prey phase lags, but they can increase the lags only if they respond to prey and predator densities (Class 3) or cues that allow accurate assessment of prey fitness (Class 4).

**Figure 2:**
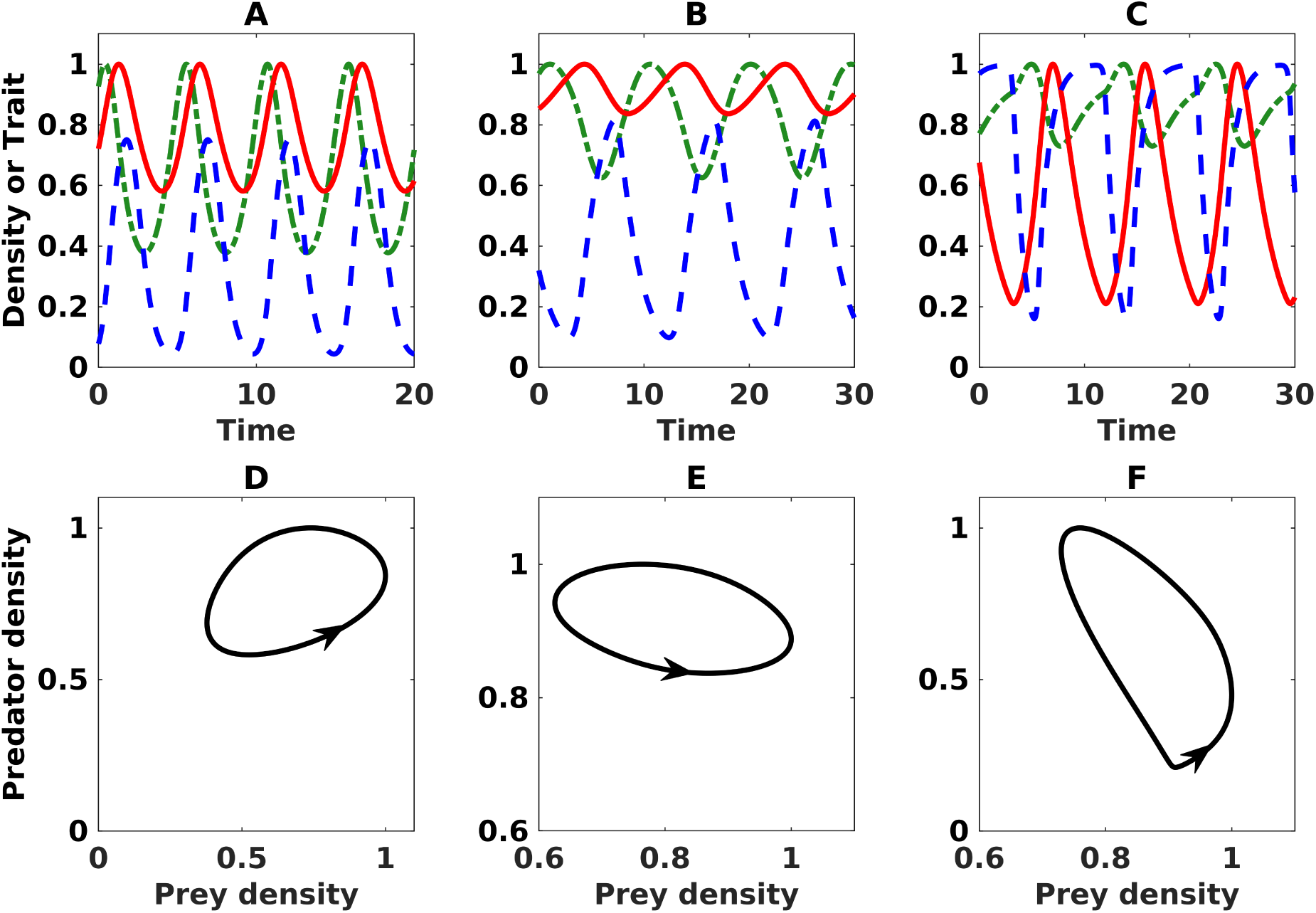
Inducible defenses (A,D) typically decrease the phase lags in predator-prey oscillations, but they can increase the lags when (B,E) induction is driven by conspecific and predator cues or (C,F) the fitness gradient. (A-C) Time series of prey density (dash-dot green), predator density (solid red), and mean defense (dashed blue) where the lag between the predator and prey oscillations is (A) less than quarter period or (B,C) between a quarter-period and a half-period. (D-F) Time series plotted in the predator-prey phase plane; arrows denote the flow of time. See Appendix S2 for parameter values.

We now address the effects of irreversibility and transgenerational responses. We predict that irreversible traits have smaller effects on phase lags than reversible traits, resulting in phase lags closer to a quarter-period. The reason is irreversibility leads to higher mean defense, which stabilizes the density dynamics of the system via reduced prey growth and predation rates. This strengthens the effects of the density dynamics relative to the effects of phenotypic plasticity, causing the phase lags to be closer to a quarter period (see Appendix S1.5.7). Thus, irreversibility can increase or decrease predator-prey phase lags, but it tends to make the lags closer to a quarter-period.

Transgenerational responses also can increase or decrease phase lags relative to intragenerational responses (see Appendix S1.5.8 for mathematical details). Longer lags occur when costs for defense are high (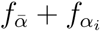 large in magnitude) because this strengthens mechanism (ii). Shorter lags occur when intraspecific prey competition is strong (*f_x_* large in magnitude) because this weakens mechanism (ii). In our simulations of transgenerational inducible defenses and irreversible inducible defenses, we did not find examples where irreversibility or transgenerational responses caused lags greater than a quarter-period when intragenerational responses did not. Based on this, we predict that lags greater than a quarter-period can only be caused by the response stimuli (specifically, Classes 2 and 4).

## 4 Discussion

In this study, we explored how the characteristics of inducible defenses influence their effects on predator-prey systems. In agreement with prior theoretical studies (Vos et al., 2004; Ramos-Jiliberto, 2003; Mougi and Kishida, 2009; Cortez, 2011; Yamamichi et al., 2011), we find that inducible defenses typically stabilize systems and reduce predator-prey phase lags. However, we also find that inducible defenses can be destabilizing and increase predator-prey phase lags, and the specific conditions under which those effects occur depend on the characteristics of the inducible defenses.

What effects are inducible defenses expected to have on predator-prey dynamics in nature? Empirical studies documenting responses to predator cues and predation cues (e.g., alarm cues from conspecifics) are common whereas fewer studies document responses to conspecific density (see references in the Introduction). If these frequencies accurately reflect the stimuli driving responses in nature, then most inducible defenses fall within Classes 1 and 3 and based on this, we predict inducible defenses are stabilizing and reduce predator-prey phase lags in most systems. This prediction agrees with laboratory studies of ciliates (Kusch, 1993), algae (Verschoor et al., 2004; Lurling et al., 2005; van der Stap et al., 2006), and waterfleas (Boeing and Ramcharan, 2010) where prey respond to predator cues and induced defenses have a stabilizing effect. However, theoretical studies argue that adaptive inducible defenses should also respond to conspecific density (Peacor 2003) or the fitness gradient (Abrams et al. 1993; Abrams 2010). This corresponds to inducible defenses in Classes 2 and 4, which can be destabilizing over a wider range of conditions and increase phase lags. These differing predictions emphasize that identifying the cues driving inducible defenses in nature is crucial in order to determine the range and likelihood of effects on predator-prey dynamics. That in turn highlights the importance of and need for empirical studies that test and report the range of stimuli that prey do and not respond to, e.g., as in Bourdeau (2010) and Gangur et al. (2017).

Regardless of what the specific effects are, we predict that it will be common for intragenerational inducible defenses to alter predator-prey dynamics. Our reasoning is that intragenerational inducible defenses have stronger effects on predator-prey dynamics when rates of induction and loss of induction are faster (Figure 1). Morphological inducible defenses are likely to have strong effects because population-level rates of change can be as fast as changes in densities (Grosklos and Cortez, 2021). Behavioral inducible defenses potentially can have even stronger effects because behavioral responses are often faster than morphological responses in the same species (Kusch, 1993; De Meester and Cousyn, 1997; Buskirk, 2002; Orizaola et al., 2012; Hammill et al., 2015). For transgenerational inducible defenses, it is less clear how commonly they will alter predator-prey dynamics because rates of induction and loss of induction depend on prey reproduction. The effects are likely to be large for species with rapid turnover, e.g, waterfleas (Agrawal et al., 1999), but for species with slow turnover rates, the magnitudes of the effects will depend on the relative rates of changes in prey densities and mean defense. Future studies on the rates of transgenerational responses are needed to estimate the strength of the effects of transgenerational defenses on empirical predator-prey dynamics.

How do the effects of inducible and evolving defense on predator-prey dynamics compare? Prior theoretical studies predict that inducible and evolving defenses have different effects on predator-prey dynamics, with inducible defenses being stabilizing and decreasing lags (Vos et al., 2004; Ramos-Jiliberto, 2003; Mougi and Kishida, 2009; Cortez, 2011; Yamamichi et al., 2011) whereas evolving defense also can be destabilizing and increase lags (Abrams and Matsuda, 1997; Jones et al., 2009; Cortez and Ellner, 2010; Cortez, 2016). This study shows that inducible defenses have the same range of effects as evolving defense, but destabilization and increased lags occur much less frequently for inducible defenses of Classes 1-3. Our analysis shows that the differing frequencies of effects are due to induction and evolution causing different feedbacks of the mean trait on its own dynamics. (Mathematically, the feedback is defined by the diagonal entry 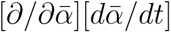 of the Jacobian.) Inducible defenses of Classes 1-3 always cause negative feedbacks and Class 4 inducible defenses and evolving defenses also cause negative feedbacks whenever intermediate trait values maximize fitness (corresponding to stabilizing selection for evolving defenses). Destabilization and increased phase lags only occur under restrictive conditions when the mean trait has a negative feedbacks on its own dynamics. In contrast, Class 4 inducible defenses and evolving defenses have positive feedbacks of the mean trait on its own dynamics whenever extreme trait values maximize fitness (corresponding to disruptive selection for evolving defenses), and such positive feedbacks are almost always destabilizing and increase phase lags. Because stabilizing and disruptive selection occur roughly equally across empirical systems (Kingsolver and Diamond, 2011), we predict evolving defenses have stabilizing versus destabilizing effects and increase versus decrease phase lags with roughly equal frequencies. In comparison, we predict inducible defenses are destabilizing and increase phase lags at a much lower frequencies, with the possible exception of Class 4 inducible defenses.

These predictions are consistent with experimental results. Specifically, in eco-evolutionary studies, evolving defenses are stabilizing (Imura et al., 2003; Kasada et al., 2014) and destabilizing (Yoshida et al., 2003, 2007; Becks et al., 2010) with roughly equal frequencies. In addition, parameterized models predict the populations are experiencing disruptive selection in all cases where evolution was destabilizing or the predator-prey phase lag was greater than a quarter period (Yoshida et al., 2003, 2007; Becks et al., 2010). In contrast, studies of inducible defenses only report stabilizing effects on predator-prey dynamics (Kusch, 1993; Verschoor et al., 2004; Lurling et al., 2005; van der Stap et al., 2006; Boeing and Ramcharan, 2010). Overall, these empirical studies lend support to our prediction that while inducible and evolving defenses can have the same range of effects on stability and predator-prey cycles, inducible defenses are much less likely to be destabilizing or increase predator-prey phase lags.

Looking beyond predator-prey systems, more work is needed to understand and compare the effects of inducible and evolving defenses on the dynamics of larger communities. For example, empirical studies of three-species communities show that inducible defenses can alter the dynamics of tri-trophic (Verschoor et al., 2004; van der Stap et al., 2007; Kishida et al., 2009), intraguild predation (Kratina et al., 2010), and two-prey-one-predator (Aŕanguiz-Acuña et al., 2011; Fischer et al., 2014) systems. Evolving defenses also can affect the dynamics of those communities (Imura et al., 2003; Hiltunen et al., 2014a; Lenhart et al., 2018). While modeling studies have explored the effects of inducible and evolving defenses in communities with more than two species (Kondoh, 2007; Urbani and Ramos-Jiliberto, 2010; Vos et al., 2004; Ramos-Jiliberto et al., 2008; Ellner and Becks, 2011; Klauschies et al., 2016), it is unknown how the mode of adaptation influences those effects. The approach taken in this study could allow one to compare models of inducible and evolving defenses and identify when they have similar versus different population-level effects.

This study extends our understanding of how inducible defense affect predator-prey dynamics by identifying how those effects depend on the characteristics of the adaptive response. It also highlights how the mode of adaptation shapes the ways in which phenotypic variation in prey defense alters the population-level dynamics of predator-prey systems.

## Acknowledgments

This study was supported by the National Science Foundation under Award DEB-1916610.

## References

Abrams, P. A. 2003. Can adaptive evolution or behaviour lead to diversification of traits determining a trade-off between foraging gain and predation risk? Ecol. Lett. 5:653–670.

Abrams, P. A. 2010. Quantitative descriptions of resource choice in ecological models. Popul. Ecol. 52:47–58.

Abrams, P. A., and H. Matsuda. 1997. Prey adaptation as a cause of predator-prey cycles. Evol. 51:1742–1750.

Abrams, P. A., H. Matsuda, and Y. Harada. 1993. Evolutionarily unstable fitness maxima and stable fitness minima of continuous traits. Evol. Ecol. 7:465–487.

Abrams, P. A., and C. J. Walters. 1996. Invulnerable prey and the paradox of enrichment. Ecol. 77:1125–1133.

Agrawal, A. A., D. D. Ackerly, F. Adler, A. E. Arnold, C. Ćaceres, D. F. Doak, E. Post, P. J. Hudson, J. Maron, K. A. Mooney, M. Power, D. Schemske, J. Stachowicz, S. Strauss, M. G. Turner, and E. Werner. 2007. Filling key gaps in population and community ecology. Front. Ecol. Environ. 5:145–152.

Agrawal, A. A., C. Laforsch, and R. Tollrian. 1999. Transgenerational induction of defences in animals and plants. Nature 401:60–63.

Anholt, B. R., D. K. Skelly, and E. E. Werner. 1996. Factors modifying antipredator behavior in larval toads. Herpetologica pages 301–313.

Aŕanguiz-Acuña, A., R. Ramos-Jiliberto, and R. O. Bustamante. 2011. Experimental evidence that induced defenses promote coexistence of zooplanktonic populations. J. Plankton. Res. 33:469–477.

Aŕanguiz-Acuña, A., R. Ramos-Jiliberto, N. Sarma, S. Sarma, R. O. Bustamante, and V. Toledo. 2010. Benefits, costs and reactivity of inducible defences: an experimental test with rotifers. Freshw. Biol. 55:2114–2122.

Auld, J. R., and R. A. Relyea. 2011. Adaptive plasticity in predator-induced defenses in a common freshwater snail: altered selection and mode of predation due to prey phenotype. Evol. Ecol. 25:189–202.

Becks, L., S. P. Ellner, L. E. Jones, and N. G. H. Jr. 2010. Reduction of adaptive genetic diversity radically alters eco-evolutionary community dynamics. Ecol. Lett. 13:989–997.

Belovsky, G. E., A. N. Laws, and J. B. Slade. 2011. Prey change behaviour with predation threat, but demographic effects vary with prey density: experiments with grasshoppers and birds. Ecol. Lett. 14:335–340.

Boeing, W. J., and C. W. Ramcharan. 2010. Inducible defences are a stabilizing factor for predator and prey populations: a field experiment. Freshw. Biol. 55:2332–2338.

Bolker, B., M. Holyoak, V. Křivan, L. Rowe, and O. Schmitz. 2003. Connecting theoretical and empirical studies of trait-mediated interactions. Ecol. 84:1101–1114.

Bolnick, D. I., P. Amarasekare, M. S. Araújo, R. Bürger, J. M. Levine, M. Novak, V. H. W. Rudolf, S. J. Schreiber, M. C. Urban, and D. A. Vasseur. 2011. Why intraspecific trait variation matters in community ecology. Trends. Ecol. Evol. 26:183–192.

Bourdeau, P. E. 2010. Cue reliability, risk sensitivity and inducible morphological defense in a marine snail. Oecologia 162:987–994.

Bulmer, M. G. 1975. Phase relations in the ten-year cycle. J. Anim. Ecol. 44:609–621.

Buskirk, J. V. 2002. Phenotypic lability and the evolution of predator-induced plasticity in tadpoles. Evol. 56:361–370.

Buskirk, J. V., and M. Arioli. 2002. Dosage response of an induced defense: how sensitive are tadpoles to predation risk? Ecol. 83:1580–1585.

Cortez, M. H. 2011. Comparing the qualitatively different effects rapidly evolving and rapidly induced defences have on predator-prey interactions. Ecol. Lett. 14:202–209.

Cortez, M. H. 2016. How the magnitude of prey genetic variation alters predator-prey eco-evolutionary dynamics. Am. Nat. 188:329–341.

Cortez, M. H. 2018. Genetic variation determines which feedbacks drive and alter predator–prey eco-evolutionary cycles. Ecol. Monogr. 88:353–371.

Cortez, M. H., and S. P. Ellner. 2010. Understanding rapid evolution in predator-prey interactions using the theory of fast-slow dynamical systems. Am. Nat. 176:E109–E127.

De Meester, L., and C. Cousyn. 1997. The change in phototactic behaviour of a *Daphnia magna* clone in the presence of fish kairomones: the effect of exposure time. Hydrobiologia 360:169–175.

Ellner, S. P., and L. Becks. 2011. Rapid prey evolution and the dynamics of two-predator food webs. Theor. Ecol. 4:133–152.

Fischer, B. B., M. Kwiatkowski, M. Ackermann, J. Krismer, S. Roffler, M. J. Suter, R. I. Eggen, and B. Matthews. 2014. Phenotypic plasticity influences the eco-evolutionary dynamics of a predator–prey system. Ecol. 95:3080–3092.

Forsman, A. 2015. Rethinking phenotypic plasticity and its consequences for individuals, populations and species. Heredity 115:276–284.

Fraker, M. E. 2008. The effect of hunger on the strength and duration of the antipredator behavioral response of green frog (*Rana clamitans*) tadpoles. Behav. Ecol. Sociobiol. 62:1201–1205.

Gangur, A. N., M. Smout, M. J. Liddell, J. E. Seymour, D. Wilson, and T. D. Northfield. 2017. Changes in predator exposure, but not in diet, induce phenotypic plasticity in scorpion venom. Proc. R. Soc. B 284:20171364.

Grosklos, G., and M. H. Cortez. 2021. Evolutionary and plastic phenotypic change can be just as fast as changes in population densities. Am. Nat. 197:47–59.

Gu, L., L. De Meester, and Z. Yang. 2023. The role of prey and predator identity in eliciting inducible defenses of daphnia. Ecol. page e4033.

Gu, L., S. Qin, N. Lu, Y. Zhao, Q. Zhou, L. Zhang, Y. Sun, Y. Huang, K. Lyu, and Z. Yang. 2020. Daphnia mitsukuri traits responding to predation cues alter its population dynamics. Ecol. Indic. 117:106587.

Halbach, U. 1970. Die ursachen der temporalvariation von *Brachionus calyciflorus Pallas (Rotatoria)*. Oecologia 4:262–318.

Hammill, E., P. Kratina, M. Vos, O. L. Petchey, and B. R. Anholt. 2015. Food web persistence is enhanced by non-trophic interactions. Oecologia 178:549–556.

Hiltunen, T., S. P. Ellner, G. Hooker, L. E. Jones, and N. G. Hairston, Jr. 2014a. Eco-evolutionary dynamics in a three-species food web with intraguild predation: Intriguingly complex. Adv. Ecol. Res. 150:41–74.

Hiltunen, T., N. G. Hairston, Jr., G. Hooker, L. E. Jones, and S. P. Ellner. 2014b. A newly discovered role of evolution in previously published consumer-resource dynamics. Ecol. Lett. 17:915–923.

Imura, D., Y. Toquenaga, and K. Fujii. 2003. Genetic variation can promote system persistence in an experimental host-parasitoid system. Popul. Ecol. 45:205–212.

Ives, A. R., and A. P. Dobson. 1987. Antipredator behavior and the population dynamics of simple predator-prey systems. Am. Nat. 130:431–447.

Jones, L. E., L. Becks, S. P. Ellner, N. G. Hairston, Jr., T. Yoshida, and G. F. Fussmann. 2009. Rapid contemporary evolution and clonal food web dynamics. Philos. Trans. R. Soc. Lond., B, Biol. Sci. 364:1579–1591.

Kasada, M., M. Yamamichi, and T. Yoshida. 2014. Form of an evolutionary tradeoff affects eco-evolutionary dynamics in a predator-prey system. PNAS 111:16035–16040.

Kingsolver, J. G., and S. E. Diamond. 2011. Phenotypic selection in natural populations: what limits directional selection? Am. Nat. 177:346–357.

Kishida, O., G. C. Trussell, A. Mougi, and K. Nishimura. 2010. Evol. ecol. of inducible morphological plasticity in predator-prey interaction: toward the practical links with popul. ecol. Popul. Ecol. 52:37–46.

Kishida, O., G. C. Trussell, and K. Nishimura. 2009. Top-down effects on antagonistic inducible defense and offense. Ecol. 90:1217–1226.

Klauschies, T., D. A. Vasseur, and U. Gaedke. 2016. Trait adaptation promotes species coexistence in diverse predator and prey communities. Ecol. Evol. 6:4141–4159.

Kondoh, M. 2007. Anti-predator defence and the complexity–stability relationship of food webs. Proc. R. Soc. B: Biol. pages 1617–1614.

Kovach-Orr, C., and G. F. Fussmann. 2013. Evolutionary and plastic rescue in multitrophic model communities. Philos. Trans. R. Soc. Lond., B, Biol. Sci. 368:20120084.

Kratina, P., E. Hammill, and B. R. Anholt. 2010. Stronger inducible defences enhance persistence of intraguild prey. J. Anim. Ecol. 79:993–999.

Kuhlmann, H., and K. Heckmann. 1985. Interspecific morphogens regulating prey-predator relationships in protozoa. Science 227:1347–1349.

Kusch, J. 1993. Behavioural and morphological changes in ciliates induced by the predator *Amoeba proteus*. Oecologia 96:354–459.

Lenhart, P. A., K. A. Jackson, and J. A. White. 2018. Heritable variation in prey defence provides refuge for subdominant predators. Proc. R. Soc. B: Biol. 285:20180523.

Lurling, M., H. Arends, W. Beekman, M. Vos, I. van der Stap, W. M. Mooij, and M. Scheffer. 2005. Effect of grazer-induced morphological changes in the green alga *Scenedesmus obliquus* on growth of the rotifer *Brachionus calyciflorus*. Pages 698–703 in Proceedings of the International Association of Theoretical and Applied Limnology. Vol. 29.

Matz, C., and S. Kjelleberg. 2005. Off the hook–how bacteria survive protozoan grazing. Trends Microbiol. 13:302–307.

Miner, B. G., S. E. Sultan, S. G. Morgan, D. K. Padilla, and R. A. Relyea. 2005. Ecological consequences of phenotypic plasticity. Trends. Ecol. Evol. 20:685–692.

Mougi, A., and O. Kishida. 2009. Reciprocal phenotypic plasticity can lead to stable predator-prey interaction. J. Anim. Ecol. 78:1172–1181.

Orizaola, G., E. Dahl, and A. Laurila. 2012. Reversibility of predator-induced plasticity and its effect at a life-history switch point. Oikos 121:44–52.

Peacor, S. D. 2003. Phenotypic modifications to conspecific density arising from predation risk assessment. Oikos 100:409–415.

Ramos-Jiliberto, R. 2003. Population dynamics of prey exhibiting inducible defenses: the role of associated costs and density-dependence. Theor. Popul. Biol. 64:221–231.

Ramos-Jiliberto, R., H. Duarte, and E. Frodden. 2008. Dynamic effects of inducible defenses in a one-prey two-predator system. Ecol. Modell. 214:242–250.

Relyea, R. A. 2004. Fine-tuned phenotypes: tadpole plasticity under 16 combinations of predators and competitors. Ecol. 85:172–179.

Relyea, R. A., and J. R. Auld. 2004. Having the guts to compete: how intestinal plasticity explains costs of inducible defenses. Ecol. Lett. 7:869–875.

Schoeppner, N. M., and R. A. Relyea. 2009. Interpreting the smells of predation: how alarm cues and kairomones induce different prey defences. Funct. Ecol. 23:1114–1121.

Shimada, M., Y. Ishii, and H. Shibao. 2010. Rapid adaptation: a new dimension for evolutionary perspectives in ecology. Popul. Ecol. 52:5–14.

Sih, A. 1992. Prey uncertainty and the balancing of antipredator and feeding needs. Am. Nat. 139:1052–1069.

Teplitsky, C., and A. Laurila. 2007. Flexible defense strategies: competition modifies investment in behavioral vs. morphological defenses. Ecol. 88:1641–1646.

Tollrian, R., S. Duggen, L. C. Weiss, C. Laforsch, and M. Kopp. 2015. Density-dependent adjustment of inducible defenses. Sci Rep 5:12736.

Tollrian, R., and C. D. Harvell. 1999. The Ecol. Evol. of Inducible Defenses. Princeton University Press, Princeton, NJ.

Urbani, P., and R. Ramos-Jiliberto. 2010. Adaptive prey behavior and the dynamics of intraguild predation systems. Ecol. Modell. 221:2628–2633.

Van Buskirk, J., M. Ferrari, D. Kueng, K. Näpflin, and N. Ritter. 2011. Prey risk assessment depends on conspecific density. Oikos 120:1235–1239.

van der Stap, I., M. Vos, B. W. Kooi, B. T. Mulling, E. van Donk, and W. M. Mooij. 2009. Algal defenses, population stability, and the risk of herbivore extinctions: a chemostat model and experiment. Ecol Res 24:1145–1153.

van der Stap, I., M. Vos, and W. M. Mooij. 2006. Linking herbivore-induced defences to population dynamics. Freshw. Biol. 51:424–434.

van der Stap, I., M. Vos, and W. M. Mooij. 2007. Inducible defenses and rotifer food chain dynamics. Hydrobiologia 593:101–110.

Van Donk, E., A. Ianora, and M. Vos. 2011. Induced defences in marine and freshwater phytoplankton: a review. Hydrobiologia 668:3–19.

Van Donk, E., M. Lürling, and W. Lampert. 1999. The Ecology and Evolution of inducible defenses, chap. Consumer-induced changes in phytoplankton: inducibility, costs, benefits, and the impact on grazers, pages 89–103. Princeton University Press Princeton, New Jersey.

Verschoor, A. M., M. Vos, and I. van der Stap. 2004. Inducible defences prevent strong population fluctuations in bi- and tritrophic food chains. Ecol. Lett. 7:1143–1148.

Vos, M., B. W. Kooi, D. L. DeAngelis, and W. M. Mooij. 2004. Inducible defences and the paradox of enrichment. Oikos 105:471–480.

Yamamichi, M., T. Klauschies, B. E. Miner, and E. van Velzen. 2019. Modelling inducible defences in predator–prey interactions: assumptions and dynamical consequences of three distinct approaches. Ecol. Lett. 22:390–404.

Yamamichi, M., C. L. Meunier, A. Peace, C. Prater, and M. A. Rúa. 2015. Rapid evolution of a consumer stoichiometric trait destabilizes consumer-producer dynamics. Oikos 124:690–969.

Yamamichi, M., T. Yoshida, and A. Sasaki. 2011. Comparing the effects of rapid evolution and phenotypic plasticity on aquatic predator-prey dynamics. Am. Nat. 178:287–304.

Yoshida, T., S. P. Ellner, L. E. Jones, B. J. M. Bohannan, R. E. Lenski, and N. G. Hairston, Jr. 2007. Cryptic population dynamics: rapid evolution masks trophic interactions. PLoS Biol. 5:1–12.

Yoshida, T., N. G. Hairston Jr., and S. P. Ellner. 2004. Evolutionary trade-off between defence against grazing and competitive ability in a simple unicellular alga. Proc. R. Soc. B: Biol. 271:1947–1953.

Yoshida, T., L. E. Jones, S. P. Ellner, G. F. Fussmann, and N. G. Hairston, Jr. 2003. Rapid evolution drives ecological dynamics in a predator-prey system. Nature 424:303–306.

