## Supplemental appendices for "The characteristics of inducible defenses influence predator-prey dynamics"

### Supporting Information for 'The characteristics of inducible defenses influence predator-prey dynamics'.

Cortez, M.H., Mila, E., and Hammill, E.

**S2 Parameter values for simulations 43**
**References 45**

### S1 Model details and analysis

This document presents additional details about the inducible defense models and our analysis. Section S1.2 addresses the intragenerational inducible defense models (1) and (2) with true breeding prey. Section S1.2 addresses the intragenerational inducible defense models (3) and (4) from Box 1 where all prey offspring are initially undefended. Section S1.3 addresses the transgenerational inducible defense models (5) and (6) from Box 1. Each section presents the model assumptions, the Jacobian of the model, and calculations showing that the continuous trait models are approximations of the discrete trait models. Section S1.4 defines the classes of inducible defenses that respond to different cues. Sections S1.5 and S1.6 provide mathematical details about the effects of inducible defenses on equilibrium stability and the predator-prey phase lags, respectively.

#### S1.1 Predator-prey model with an intragenerational inducible defense and true breeding prey

For this model, we assume prey can change their phenotype during their lifetime, induction is reversible, and prey offspring initially have the same phenotype as their parent. These assumptions are made in nearly all prior modeling studies of inducible defenses. The mean trait value can take on values in the range $[\alpha_{min}, \alpha_{max}]$ where $\alpha_{min}$ and $\alpha_{max}$ are the minimum and maximum allowable trait values.

The intragenerational inducible defense model with true breeding prey is

$$\begin{aligned}
\frac{dx}{dt} &= \overbrace{xf(x,\bar{\alpha},\alpha_i)}^{\text{reproduction}} - \overbrace{xg(x,y,\bar{\alpha},\alpha_i)}^{\text{predation}}\Bigg|_{\alpha_i=\bar{\alpha}} \\
\frac{dy}{dt} &= \overbrace{yh(x,y,\bar{\alpha})}^{\text{harvesting}} - \overbrace{ym(y)}^{\text{mortality}} \\
\frac{d\bar{\alpha}}{dt} &= \underbrace{\overbrace{(\alpha_{max}-\bar{\alpha})\epsilon_I P_I}^{\text{induction}} - \overbrace{(\bar{\alpha}-\alpha_{min})\epsilon_L P_L}^{\text{loss of induction}}}_{\text{phenotypic plasticity}} + \underbrace{V(\bar{\alpha})(f_{\alpha_i}-g_{\alpha_i})}_{\text{phenotypic sorting}}\Bigg|_{\alpha_i=\bar{\alpha}}
\end{aligned} \tag{S1}$$

where $\epsilon_I P_I$ and $\epsilon_L P_L$ are the rates of induction and loss of induction and $\epsilon_I$ and $\epsilon_L$ denote the maximum rates. In the equations, $\alpha_i$ denotes an individual's trait value and $\bar{\alpha}$ is the mean trait value.

##### S1.1.1 Model assumptions

**Equilibrium assumptions:** Throughout, we assume model (S1) has a coexistence equilibrium point $\rho = (x^*, y^*, \alpha^*)$ where both species have positive densities ($x^*, y^* > 0$). We also assume that the sign of the determinant of the Jacobian is negative ($|J| < 0$); the equilibrium point is a saddle when this condition is not met and it is unclear if the analysis of such points yields biological insight.

**Assumptions about density dynamics:** We assume the growth rates of the prey and predator are determined by the mean trait value. The functions defining reproduction ($xf$), predation ($xg$), harvesting ($yh$), and predator mortality ($ym$) satisfy mathematical conditions consistent with intraspecific competition and a predatory interaction. First, the prey per capita reproduction rate is a decreasing function of prey density due to intraspecific competition ($\partial f/\partial x < 0$). This assumption is satisfied by Lotka-Volterra, Beverton-Holt, and other models of intraspecific density regulation where per capita growth rates decrease with intraspecific density. Second, the predation and harvesting rates are increasing functions of prey density and predator density ($\partial xg/\partial x > 0$, $\partial xg/\partial y > 0$, $\partial yh/\partial x > 0$, $\partial yh/\partial y > 0$). These assumptions are satisfied by Type I, II, and III functional responses as well as other saturating functional responses where predation rates increase with increases in prey or predator density. Third, the per predator predation rate ($h$) is a non-increasing functions of predator density ($\partial h/\partial y \leq 0$). The per predator predation rate is independent of predator density ($\partial h/\partial y = 0$) for predator-independent functional responses (like the Type I, II, and III functional responses) and it is a decreasing function of predator density ($\partial h/\partial y < 0$) for predator-dependent functional responses like the Beddington-DeAngelis (Beddington 1975, DeAngelis et al. 1975) and Crowley-Martin (Crowley and Martin 1989) functional responses that account for predator interference. Fourth, the per capita predator mortality rate is a non-decreasing function of predator density ($dm/dy \geq 0$). This assumption accounts for negative intraspecific interactions between the predators that don't involve predation (e.g., intraspecific fighting).

The functions defining reproduction, predation, and harvesting also satisfy mathematical conditions that align with the costs and benefits of an induced defense. First, the prey per capita reproduction and predation rates are a decreasing functions of the individual's trait value and the mean trait value ($\partial f/\partial\bar{\alpha} < 0$, $\partial f/\partial\alpha_i < 0$, $\partial g/\partial\bar{\alpha} < 0$, and $\partial g/\partial\alpha_i < 0$). Combined, these conditions result in a trade-off where increased individual defense comes at the benefit of reduced predation and the cost of reduced reproductive output. Second, we assume the predator's harvesting rate is a decreasing function of the mean trait ($\partial xg/\partial\bar{\alpha} < 0$); this is consistent with increased defense leading to lower predation rates.

**Assumptions about trait dynamics:** The trait dynamics can be separated into a phenotypic plasticity component and a phenotypic sorting component (Yamamichi et al. 2019). The phenotypic plasticity component accounts for how mean trait changes due to induction and loss of induction. We write the induction rate in terms of a fraction ($P_I$) of its maximum rate ($\epsilon_I$) and we write the loss of induction rate in terms of a fraction ($P_L$) of its maximum ($\epsilon_L$). The fractions $P_I$ and $P_L$ define how the rates of induction and loss of induction depend on prey and predator densities; see Section S1.4.

The phenotypic sorting component of the trait dynamics accounts for how the mean trait changes due to differential differential survival and reproduction of the different phenotypes. In that term, $V(\bar{\alpha})$ is the trait variance, which depends on the mean trait value because the variance must necessarily converge to zero as the mean trait value approaches $\alpha_{min}$ or $\alpha_{max}$. The derivative $f_{\alpha_i} - g_{\alpha_i}$ is the individual fitness gradient; it determines the direction of increasing fitness. The phenotypic sorting term arises because all prey do not have identical trait values, e.g., due imperfect switching; see Section S1.1.4.

##### S1.1.2 Jacobian

When evaluated at an equilibrium, $\rho = (x^*, y^*, \alpha^*)$, the Jacobian of model (S1) simplifies to

$$J = \begin{bmatrix} \frac{\partial}{\partial x}\frac{dx}{dt} & \frac{\partial}{\partial y}\frac{dx}{dt} & \frac{\partial}{\partial \bar{\alpha}}\frac{dx}{dt} \\ \frac{\partial}{\partial x}\frac{dy}{dt} & \frac{\partial}{\partial y}\frac{dy}{dt} & \frac{\partial}{\partial \bar{\alpha}}\frac{dy}{dt} \\ \frac{\partial}{\partial x}\frac{d\bar{\alpha}}{dt} & \frac{\partial}{\partial y}\frac{d\bar{\alpha}}{dt} & \frac{\partial}{\partial \bar{\alpha}}\frac{d\bar{\alpha}}{dt} \end{bmatrix} = \begin{bmatrix} J_{11} & J_{12} & J_{13} \\ J_{21} & J_{22} & J_{23} \\ J_{31} & J_{32} & J_{33} \end{bmatrix}$$

$$= \begin{bmatrix} x(f_x - g_x) & -xg_y & f_{\bar{\alpha}} - g_{\bar{\alpha}} + f_{\alpha_i} - g_{\alpha_i} \\ yh_x & y(h_y - m_y) & yh_{\bar{\alpha}} \\ J_{31} & J_{32} & J_{33} \end{bmatrix} \tag{S2}$$

where subscript variables denote partial derivatives (e.g., $f_x = \partial f/\partial x$) and

$$J_{31} = \overbrace{\epsilon_I(\alpha_{max} - \bar{\alpha})\frac{\partial P_I}{\partial x} - \epsilon_L(\bar{\alpha} - \alpha_{min})\frac{\partial P_L}{\partial x}}^{\text{phenotypic plasticity}} + \overbrace{V(\bar{\alpha})(f_{x\alpha_i} - g_{x\alpha_i})}^{\text{phenotypic sorting}} \tag{S3}$$

$$J_{32} = \overbrace{\epsilon_I(\alpha_{max} - \bar{\alpha})\frac{\partial P_I}{\partial y} - \epsilon_L(\bar{\alpha} - \alpha_{min})\frac{\partial P_L}{\partial y}}^{\text{phenotypic plasticity}} + \overbrace{V(\bar{\alpha})(-g_{y\alpha_i})}^{\text{phenotypic sorting}} \tag{S4}$$

$$\begin{aligned} J_{33} = & \overbrace{-\epsilon_I P_I - \epsilon_L P_L + \epsilon_I(\alpha_{max} - \bar{\alpha})\frac{\partial P_I}{\partial \bar{\alpha}} - \epsilon_L(\bar{\alpha} - \alpha_{min})\frac{\partial P_L}{\partial \bar{\alpha}}}^{\text{phenotypic plasticity}} \\ & + \overbrace{V(\bar{\alpha})(f_{\bar{\alpha}\alpha_i} - g_{\bar{\alpha}\alpha_i} + f_{\alpha_i\alpha_i} - g_{\alpha_i\alpha_i}) + V'(\bar{\alpha})(f_{\alpha_i} - g_{\alpha_i})}^{\text{phenotypic sorting}} \end{aligned} \tag{S5}$$

The assumptions about the functions $f$, $g$, $h$, and $m$ determine the signs of the entries in the first two rows of the Jacobian. The sign of $J_{11}$ can be positive or negative depending on whether the predator overexploits its prey (e.g., the predator nullcline is on the left side of the hump of the prey nullcline in a Rosenzweig-MacArthur model). The other entries have known signs: $J_{12} < 0$ and $J_{21} > 0$ because of the predatory interaction between the species; $J_{22} \leq 0$ because predator interference and the intraspecific predator interactions; and $J_{23} < 0$ because increased defense reduces predation rates. The sign of $J_{13}$ can be positive or negative depending on whether increases in mean defense increase or decrease the fitness of individual prey, respectively. For example, positive values can occur if there is a shared defense (such as an excreted chemical defense), which results in all prey benefiting when mean levels of defense are higher. Negative values can occur if (i) more defended prey produce less of a shared resource, which results in lower individual fitness when mean levels of defense are higher or (ii) more defended prey are more aggressive, which results in better defended prey having stronger negative intraspecific effects.

The entries of the bottom row show how each variable affects the trait dynamics. The phenotypic plasticity terms define the sensitivity of the induction and loss of induction rates to changes in prey density (terms in $J_{31}$), predator density (terms in $J_{32}$), and mean level of defense (terms in $J_{33}$). The signs of these terms are determined by the stimulus type; see Section S1.4.

The phenotypic sorting terms define how the state variables affect the prey individual fitness gradient; the biological interpretations of the derivatives are listed below. The phenotypic sorting terms in entries $J_{31}$ and $J_{32}$ define whether the individual fitness gradient increases or decreases with variation in prey and predator densities, respectively. In entry $J_{33}$, the first phenotypic sorting term determines the concavity of the fitness surface and whether extreme trait values are optimal (positive values; corresponding to disruptive selection) or intermediate trait values are optimal (negative values; corresponding to stabilizing selection). The second phenotypic sorting term accounts for the effects changes in trait variation (which decreases as the mean trait value approaches $\alpha_{min}$ or $\alpha_{max}$ and increases when the mean trait moves away from those values) and the slope of the fitness surface. In simulations the phenotypic sorting terms were always much smaller in magnitude than the phenotypic plasticity terms, unless rates of induction were very small ($\epsilon_I$ and $\epsilon_L$ close to zero). For completeness, our analysis discusses how the phenotypic sorting terms affect equilibrium stability and predator-prey phase lags, but we expect the effects of the phenotypic sorting are negligibly small compared to the effects of phenotypic plasticity in most systems.

- $\underline{f_{x\alpha_i} - g_{x\alpha_i}}$: Effect of increased prey density on the benefits and costs of increased defense. Positive values imply greater benefits (or reduced costs) of increased individual defense when prey density is higher. Negative values imply greater costs (or reduced benefits) of increased individual defense when prey density is higher.

- $\underline{-g_{y\alpha_i}}$: Effect of increased predator on the benefits of increased defense. Positive and negative values, respectively, imply greater and reduced benefits of increased individual defense when predator density is higher. Throughout we assume $-g_{y\alpha_i} > 0$. Our reasoning is that it is very difficult to construct models where $-g_{y\alpha_i} < 0$, which suggests that that condition is unlikely to be satisfied in natural systems.

- $\underline{f_{\bar{\alpha}\alpha_i} - g_{\bar{\alpha}\alpha_i} + f_{\alpha_i\alpha_i} - g_{\alpha_i\alpha_i}}$: Effect of increased mean defense on the benefits and costs of increased defense. Positive values imply greater benefits (or reduced costs) of increased individual defense when mean prey defense is higher. Negative values imply greater costs (or reduced benefits) of increased individual defense when mean prey defense is higher.

##### S1.1.3 Relationship to evolving defense models

The form of the continuous trait inducible defense model (S1) is similar to the form of evolving defense models based on the quantitative genetics framework in Abrams et al. (1993). In particular, the evolving defense model in (Cortez 2011, 2016) has identical prey and predator density equations and the trait equation is

$$\frac{d\bar{\alpha}}{dt} = \overbrace{V(\bar{\alpha})}^{\text{variance}} \overbrace{\frac{\partial}{\partial \alpha_i}(f-g)\bigg|_{\alpha_i=\alpha}}^{\text{individuals fitness gradient}} = V(\bar{\alpha})(f_{\alpha_i} - g_{\alpha_i})\bigg|_{\alpha_i=\bar{\alpha}}. \tag{S6}$$

In the trait equation, $V(\bar{\alpha})$ is the additive genetic variance of the trait and $f_{\alpha_i} - g_{\alpha_i}$ is the individual fitness gradient. The evolving defense model can derived a clonal model following the steps in Section S1.1.4.

If $\epsilon_I = \epsilon_L = 0$, then the continuous trait inducible defense model (S1) reduces to an evolving defense model. This means that the effects of an evolving defense on predator-prey dynamics are identical to the effects of the phenotypic sorting terms of the inducible defense model (S1). In addition, the effects of an evolving defense on equilibrium stability and predator-prey phase lags are nearly identical to those of a Class 4 inducible defense (defined in Section S1.4). Intuitively, the reason is that Class 4 inducible defenses and evolving defense both maximize individual fitness. Mathematically, the reason is that the Jacobian entries for Class 4 inducible defenses almost always have the same sign as the Jacobian for an evolving defense. The only exception is when the fitness surface is very flat such that $f_{\bar{\alpha}\alpha_i} - g_{\bar{\alpha}\alpha_i} + f_{\alpha_i\alpha_i} - g_{\alpha_i\alpha_i}$ is very close to zero. In this case, entry $J_{33}$ of the Jacobian for the inducible defense model can be negative when the corresponding entry of the Jacobian for the evolving defense model is positive. We do not expect this situation to arise often in nature.

##### S1.1.4 Derivation from multimorphic discrete trait models

Here, we show that the continuous trait model (S1) can approximate the dynamics of discrete trait models. First, we extend the derivations in prior studies (Cortez 2011, Yamamichi et al. 2019) by showing how model (S1) approximates the dynamics of a dimorphic discrete trait model. Second, we use similar calculations to show that a continuous trait model with a similar form can be derived from a discrete trait model with more than 2 phenotypes. The latter derivation is unlikely to be accurate in all cases because it necessarily requires a reduction in model dimension. However, we provide it because it provides additional evidence supporting our claim that the results from our continuous trait model (S1) can provide useful insight about the dynamics of systems where prey defense can take on many different levels.

**Approximation of dimorphic model:** Assume an individual prey's defense level can be the values $\alpha_1 = \alpha_{min}$ or $\alpha_2 = \alpha_{max}$. Let $x_i$ be the density of prey with defense level $\alpha_i$. Let $P_1^2$ be the induction rate for individuals of phenotype 1 and $P_2^1$ be the loss of induction rate for individuals with phenotype 2. The dynamics of the prey classes and predator are given by

$$\begin{aligned}
\frac{dx_1}{dt} &= \overbrace{x_1 f(x_1, x_2, \alpha_1)}^{\text{reproduction}} - \overbrace{x_1 g(x_1, x_2, \alpha_1)}^{\text{predation}} \overbrace{-x_1 P_i^2}^{\text{induction}} \quad \overbrace{+x_2 P_2^1}^{\text{loss of induction}} \\
\frac{dx_2}{dt} &= \overbrace{x_2 f(x_1, x_2, \alpha_2)}^{\text{reproduction}} - \overbrace{x_2 g(x_1, x_2, \alpha_2)}^{\text{predation}} \overbrace{+x_1 P_1^2}^{\text{induction}} \quad \overbrace{-x_2 P_2^1}^{\text{loss of induction}} \\
\frac{dy}{dt} &= \overbrace{yh(x_1, x_2, y)}^{\text{harvesting}} - \overbrace{ym(y)}^{\text{mortality}}
\end{aligned} \tag{S7}$$

where $f(x_1, x_2, \alpha_i)$ and $g(x_1, x_2, y, \alpha_i)$ are the reproduction and predation rates of individuals with phenotype $\alpha_i$. We assume the functional forms of the prey reproduction rates, prey predation rates, and predator harvesting rates depend on total prey density ($x = \sum_i x_i$), predator density (when applicable), the average level of defense ($\bar{\alpha} = \sum_i \alpha_i x_i / x$), and the individual's phenotype ($\alpha_i$, when applicable). Mathematically, we assume $f(x_1, x_2, \alpha_i) = f(x, \bar{\alpha}, \alpha_i)$, $g(x_1, x_2, y, \alpha_i) = g(x, y, \bar{\alpha}, \alpha_i)$, and $h(x_1, x_2, y, ) = h(x, y, \bar{\alpha})$.

We now derive equations that approximate the dynamics of the total density and the average level of defense of the prey population. To help simplify the calculations, we use $q_i = x_i/x$ to denote the frequency of phenotype $i$. The dynamics of total prey density are approximated as

$$\frac{dx}{dt} = \sum_i x_i f(x, \bar{\alpha}, \alpha_i) - x_i g(x, y, \bar{\alpha}, \alpha_i) \tag{S8}$$

$$= x \left[ \sum q_i f(x, \bar{\alpha}, \alpha_i) - q_i g(x, \bar{\alpha}, \alpha_i) \right] \tag{S9}$$

$$\approx x \left[ f(x, \bar{\alpha}, \bar{\alpha}) - g(x, y, \bar{\alpha}, \bar{\alpha}) \right]. \tag{S10}$$

To get the dynamics of the average trait value, we compute

$$\frac{d\bar{\alpha}}{dt} = \frac{1}{x^2} \left[ x \sum_i \alpha_i \frac{dx_i}{dt} - \frac{dx}{dt} \sum_i \alpha_i x_i \right] \tag{S11}$$

$$= \overbrace{\sum_i q_i \left[ f(x, \bar{\alpha}, \alpha_i) - g(x, y, \bar{\alpha}, \alpha_i) \right] (\alpha_i - \bar{\alpha})}^{Q_{sorting},\ \text{phenotypic sorting}} + \overbrace{-\alpha_1 q_1 P_1^2 + \alpha_1 q_2 P_2^1 + \alpha_2 q_1 P_1^2 - \alpha_2 q_2 P_2^1}^{Q_{plasticity},\ \text{phenotypic plasticity}}. \tag{S12}$$

First consider the phenotypic sorting terms in equation (S12); they account for changes in the mean trait value due to differential reproduction and survival of the different phenotypes. Using $x_1 = x(\alpha_2 - \bar{\alpha})/(\alpha_2 - \alpha_1)$, $x_2 = x(\bar{\alpha} - \alpha_1)/(\alpha_2 - \alpha_1)$, and $q_i = x_i/x$, we get

$$Q_{sorting} = q_1 q_2 (\alpha_2 - \alpha_1)^2 \left[ \frac{f(x, \bar{\alpha}, \alpha_2) - g(x, y, \bar{\alpha}, \alpha_2) - f(x, \bar{\alpha}, \alpha_1) + g(x, y, \bar{\alpha}, \alpha_1)}{\alpha_2 - \alpha_1} \right] \tag{S13}$$

$$\approx V(\bar{\alpha}) \left[ \frac{\partial f}{\partial \alpha_i}(x, \bar{\alpha}, \alpha_i) - g \frac{\partial g}{\partial \alpha_i}(x, y, \bar{\alpha}, \alpha_i) \right] \Big|_{\alpha_i = \bar{\alpha}} \tag{S14}$$

where $V(\bar{\alpha}) = q_1 q_2 (\alpha_2 - \alpha_1)^2 = (\alpha_2 - \bar{\alpha})(\bar{\alpha} - \alpha_1)$ is the population variance of the trait. The error in the approximation was derived in Cortez (2011).

Second, consider the phenotypic plasticity terms in equation (S12); they account for changes in the mean trait value due to induction and loss of induction. Using the same substitutions as above, and after some algebra, the phenotypic plasticity terms can be written as

$$Q_{plasticity} = (\alpha_1 - \alpha_1) q_i P_1^2 - (\alpha_2 - \alpha_1) q_2 P_2^1 = (\alpha_2 - \bar{\alpha}) P_1^2 - (\bar{\alpha} - \alpha_1) P_2^1. \tag{S15}$$

For convenience, we write induction and loss of induction rates in terms of their maximum rates, i.e.,$P_1^2 = \epsilon_I P_I$ and $P_2^1 = \epsilon_L P_L$ where $\epsilon_I = \max P_1^2$, $\epsilon_L = \max P_2^1$, and $0 \leq P_I, P_L \leq 1$. Combining the formulas for $Q_{sorting}$ and $Q_{plasticity}$ and the formulas for the other derivatives yields the continuous trait model (S1).

**Approximation of multimorphic model:** Assume an individual prey's defense level can be $\alpha_i$ ($1 \leq i \leq n$) where $\alpha_1 = \alpha_{min}$, $\alpha_i < \alpha_{i+1}$, and $\alpha_n = \alpha_{max}$. Let $x_i$ be the density of prey with defense level $\alpha_i$. We assume individuals can only switch from their current defense level to the next higher or next lower level of defense, where $P_i^{i+1}$ is the induction rate at which individuals of phenotype $i$ switch to phenotype $i+1$ and $P_i^{i-1}$ is the loss of induction rate at which individuals of phenotype $i$ switch to phenotype $i-1$. Under this assumption, the dynamics of the prey classes and predator are given by

$$\begin{aligned}
\frac{dx_1}{dt} &= \overbrace{x_1 f(x_1, ..., x_n, \alpha_1)}^{\text{reproduction}} - \overbrace{x_1 g(x_1, ..., x_n, \alpha_1)}^{\text{predation}} \overbrace{-x_1 P_i^2}^{\text{induction}} \quad \overbrace{+x_2 P_2^1}^{\text{loss of induction}} \\
\frac{dx_i}{dt} &= \overbrace{x_i f(x_1, ..., x_n, \alpha_i)}^{\text{reproduction}} - \overbrace{x_i g(x_1, ..., x_n, \alpha_i)}^{\text{predation}} \overbrace{-x_i P_i^{i+1} + x_{i-1} P_{i-1}^i}^{\text{induction}} \overbrace{-x_i P_i^{i-1} + x_{i+1} P_{i+1}^i}^{\text{loss of induction}} \\
\frac{dx_n}{dt} &= \overbrace{x_n f(x_1, ..., x_n, \alpha_n)}^{\text{reproduction}} - \overbrace{x_n g(x_1, ..., x_n, \alpha_n)}^{\text{predation}} \overbrace{+x_{n-1} P_{n-1}^n}^{\text{induction}} \quad \overbrace{-x_n P_n^{n-1}}^{\text{loss of induction}} \\
\frac{dy}{dt} &= \overbrace{y h(x_1, ..., x_n, y)}^{\text{harvesting}} - \overbrace{y m(y)}^{\text{mortality}}
\end{aligned} \tag{S16}$$

where $f(x_1, ..., x_n, \alpha_i)$ and $g(x_1, ..., x_n, y, \alpha_i)$ are the reproduction and predation rates of individuals with phenotype $\alpha_i$. We assume the functional forms of the prey reproduction rates, prey predation rates, and predator harvesting rates depend on total prey density ($x = \sum_i x_i$), predator density (when applicable), the average level of defense ($\bar{\alpha} = \sum_i \alpha_i x_i / x$), and the individual's phenotype ($\alpha_i$, when applicable). Mathematically, we assume $f(x_1, ..., x_n, \alpha_i) = f(x, \bar{\alpha}, \alpha_i)$, $g(x_1, ..., x_n, y, \alpha_i) = g(x, y, \bar{\alpha}, \alpha_i)$, and $h(x_1, ..., x_n, y, ) = h(x, y, \bar{\alpha})$.

We now derive equations that approximate the dynamics of the total density and the average level of defense of the prey population. To help simplify the calculations, we use $q_i = x_i/x$ to denote the frequency of phenotype $i$. The dynamics of total prey density are approximated as

$$\frac{dx}{dt} = \sum_i x_i f(x, \bar{\alpha}, \alpha_i) - x_i g(x, y, \bar{\alpha}, \alpha_i) \tag{S17}$$

$$= x \left[ \sum q_i f(x, \bar{\alpha}, \alpha_i) - q_i g(x, \bar{\alpha}, \alpha_i) \right] \tag{S18}$$

$$\approx x \left[ f(x, \bar{\alpha}, \bar{\alpha}) - g(x, y, \bar{\alpha}, \bar{\alpha}) \right]. \tag{S19}$$

To get the dynamics of the average trait value, we compute

$$\frac{d\bar{\alpha}}{dt} = \frac{1}{x^2} \left[ x \sum_i \alpha_i \frac{dx_i}{dt} - \frac{dx}{dt} \sum_i \alpha_i x_i \right] \tag{S20}$$

$$= \overbrace{\sum_i q_i \left[f(x,\bar{\alpha},\alpha_i) - g(x,y,\bar{\alpha},\alpha_i)\right](\alpha_i - \bar{\alpha})}^{Q_{sorting},\text{ phenotypic sorting}}$$
$$+ \overbrace{\sum_i \alpha_i(-q_i P_i^{i+1} + q_{i-1}P_{i-1}^{i} - q_i P_i^{i-1} + q_{i+1}P_{i+1}^{i})}^{Q_{plasticity},\text{ phenotypic plasticity}} . \tag{S21}$$

First consider the phenotypic sorting terms in equation (S21); they account for changes in the mean trait value due to differential reproduction and survival of the different phenotypes. Using the definition of $\bar{\alpha}$ and $\sum_i q_i = 1$, we write

$$\alpha_i - \bar{\alpha} = \alpha_i \sum_j q_j - \sum_j \alpha_j q_j = \sum_j (\alpha_i - \alpha_j) q_j \tag{S22}$$

and substitute to get

$$Q_{sorting} = \sum_{i \neq j} q_i q_j (\alpha_i - \alpha_j)^2 \left[\frac{f(x,\bar{\alpha},\alpha_i) - g(x,y,\bar{\alpha},\alpha_i) - f(x,\bar{\alpha},\alpha_j) + g(x,y,\bar{\alpha},\alpha_j)}{\alpha_i - \alpha_j}\right] \tag{S23}$$

$$\approx V \sum_{i \neq j} \frac{q_i q_j (\alpha_i - \alpha_j)^2}{V} \left[\frac{\partial f}{\partial \alpha_i}(x,\bar{\alpha},[\alpha_i - \alpha_j]/2) - \frac{\partial g}{\partial \alpha_i}(x,y,\bar{\alpha},[\alpha_i - \alpha_j]/2)\right] \tag{S24}$$

$$\approx V \left[\frac{\partial f}{\partial \alpha_i}(x,\bar{\alpha},\alpha_i) - g\frac{\partial g}{\partial \alpha_i}(x,y,\bar{\alpha},\alpha_i)\right]\Big|_{\alpha_i = \bar{\alpha}} \tag{S25}$$

where $V = \sum_{i \neq j} q_i q_j (\alpha_i - \alpha_j)^2$ is the population variance of the trait. The second line uses a centered difference to approximate the derivatives of $f$ and $g$ and the third line approximates the derivatives of $f$ and $g$ with respect to $\alpha_i$ at $\alpha_i = \bar{\alpha}$ using the weighted sums of the approximated derivatives. The end result is that the phenotypic sorting part of the trait dynamics is approximated by the product of the population phenotypic variance and the fitness gradient.

Second, consider the phenotypic plasticity terms in equation (S21); they account for changes in the mean trait value due to induction and loss of induction. Using equation (S22) and after some algebra, the phenotypic plasticity terms can be written as

$$Q_{plasticity} = \sum_{i \neq n} (\alpha_{i+1} - \alpha_i) q_i P_i^{i+1} - \sum_{i \neq n} (\alpha_{i+1} - \alpha_i) q_{i+1} P_{i+1}^{i} \tag{S26}$$

$$= (\alpha_n - \bar{\alpha}) \frac{\sum_{i \neq n} q_i (\alpha_{i+1} - \alpha_i) P_i^{i+1}}{\sum_{j \neq n} (\alpha_n - \alpha_j) q_j} - (\bar{\alpha} - \alpha_1) \frac{\sum_{i \neq n} q_i (\alpha_{i+1} - \alpha_i) P_{i+1}^{i}}{\sum_{j \neq 1} (\alpha_j - \alpha_1) q_j} \tag{S27}$$

$$= (\alpha_n - \bar{\alpha}) \hat{P}_I - (\bar{\alpha} - \alpha_1) \hat{P}_L. \tag{S28}$$

The first term of equation (S27) shows that the average induction rate, $\hat{P}_I$, is the average change in the trait values due to induction ($\sum_{i \neq n} q_i (\alpha_{i+1} - \alpha_i) P_i^{i+1}$) divided by the average displacement from the maximum trait value ( $\sum_{j \neq n} (\alpha_n - \alpha_j) q_j$). When multiplied by $\alpha_n - \bar{\alpha}$, this yields the average change in the mean trait value due to induction. Similarly, the second term of equation (S27) shows that the average loss of induction rate, $\hat{P}_L$, is the average change in the trait due to loss of induction ($\sum_{i\neq n} q_i(\alpha_{i+1} - \alpha_i)P^i_{i+1}$) divided by the average displacement from the minimum trait value ($\sum_{j\neq 1}(\alpha_j - \alpha_1)q_j$). When multiplied by $\bar{\alpha} - \alpha_1$, this yields the average change in the mean trait value due to loss of induction.

For convenience, we write the average rates of induction and loss of induction in terms of their maximum rates, i.e.,$\hat{P}_I = \epsilon_I P_I$ and $\hat{P}_L = \epsilon_L P_L$ where $\epsilon_I = \max P_I$, $\epsilon_L = \max P_L$, and $0 \leq P_I, P_L \leq 1$. Combining the formulas for $Q_{sorting}$ and $Q_{plasticity}$ and the formulas for the other derivatives yields the continuous trait model (S1).

Note that the derivatives of the average induction and loss of rates are defined by the sums of the induction and loss of induction rates of the prey classes. For example, if increased predator density increases the induction rates for all classes ($\partial P_i^{i+1}/\partial y \geq 0$) and decreases the loss of induction rates for all classes ($\partial P_i^{i-1}/\partial y \leq 0$), then increased predator density will increase the average induction rate ($\partial P_I/\partial y \geq 0$) and decrease the average loss of induction rate ($\partial P_I/\partial y \leq 0$). Second, except in the special case of a dimorphic trait ($n = 2$), the continuous trait model (S1) will necessarily have fewer dimensions than the multimorphic model (S16). Thus, the continuous trait model (S1) provides a useful starting point for understanding how continuous and discrete trait inducible defenses affect predator-prey dynamics, but it may not fully capture all of the dynamics produced by the multimorphic model (S16).

#### S1.2 Predator-prey model with an intragenerational inducible defense and all prey offspring are initially uninduced

For this model, we assume prey can change their phenotype during their lifetime, induction is reversible, and all prey offspring are initially uninduced. The mean trait value can take on values in the range $[\alpha_{min}, \alpha_{max}]$ where $\alpha_{min}$ and $\alpha_{max}$ are the minimum and maximum allowable trait values. The switch after birth model where all prey offspring are initially uninduced is

$$\begin{aligned}
\frac{dx}{dt} &= \overbrace{xf(x,\bar{\alpha},\alpha_i)}^{\text{reproduction}} - \overbrace{xg(x,y,\bar{\alpha},\alpha_i)}^{\text{predation}}\Bigg|_{\alpha_i=\bar{\alpha}} \\
\frac{dy}{dt} &= \overbrace{yh(x,y,\bar{\alpha})}^{\text{harvesting}} - \overbrace{ym(y)}^{\text{mortality}} \\
\frac{d\bar{\alpha}}{dt} &= \underbrace{\overbrace{(\alpha_{max}-\bar{\alpha})\epsilon_I P_I}^{\text{induction}} - \overbrace{(\bar{\alpha}-\alpha_{min})\epsilon_L P_L}^{\text{loss of induction}}}_{\text{phenotypic plasticity}} \underbrace{- (\bar{\alpha}-\alpha_{min})f - V(\bar{\alpha})g_{\alpha_i}}_{\text{phenotypic sorting}}\Bigg|_{\alpha_i=\bar{\alpha}}
\end{aligned} \tag{S29}$$

where $\epsilon_I P_I$ and $\epsilon_L P_L$ are the rates of induction and loss of induction and $\epsilon_I$ and $\epsilon_L$ denote the maximum rates. In the equations, $\alpha_i$ denotes an individual's trait value and $\bar{\alpha}$ is the mean trait value. An important special case of this model is when the prey has an irreversible trait; this corresponds to $\epsilon_L = 0$.

##### S1.2.1 Model assumptions

The assumptions about equilibria and the density dynamics are the same as those in Section S1.1.1.

**Assumptions about trait dynamics:** As before, we separate the trait dynamics into a phenotypic plasticity component and a phenotypic sorting component. The phenotypic plasticity components of models (S29) and (S1) are identical.

The phenotypic sorting terms of the two models differ because of the different assumptions about prey reproduction. The phenotypic sorting component accounts for changes in the mean trait value due to reproduction and predation. In particular, reproduction term is negative because reproduction causes the mean trait value to shift to smaller values because all offspring are initially undefended. The predation term is positive because more defended prey have lower predation rates, which causes the mean trait value to shift to larger values. In the predation term, the trait variances, $V(\bar{\alpha})$, depends on the mean trait value because the variance necessarily converges to zero as the mean trait value approaches $\alpha_{min}$ or $\alpha_{max}$.

##### S1.2.2 Jacobian

When evaluated at an equilibrium, $\rho = (x^*, y^*, \alpha^*)$, the Jacobian of model (S29) simplifies to

$$\begin{aligned} J &= \begin{bmatrix} \frac{\partial}{\partial x}\frac{dx}{dt} & \frac{\partial}{\partial y}\frac{dx}{dt} & \frac{\partial}{\partial \bar{\alpha}}\frac{dx}{dt} \\ \frac{\partial}{\partial x}\frac{dy}{dt} & \frac{\partial}{\partial y}\frac{dy}{dt} & \frac{\partial}{\partial \bar{\alpha}}\frac{dy}{dt} \\ \frac{\partial}{\partial x}\frac{d\bar{\alpha}}{dt} & \frac{\partial}{\partial y}\frac{d\bar{\alpha}}{dt} & \frac{\partial}{\partial \bar{\alpha}}\frac{d\bar{\alpha}}{dt} \end{bmatrix} = \begin{bmatrix} J_{11} & J_{12} & J_{13} \\ J_{21} & J_{22} & J_{23} \\ J_{31} & J_{32} & J_{33} \end{bmatrix} \\ &= \begin{bmatrix} x(f_x - g_x) & -xg_y & f_{\bar{\alpha}} - g_{\bar{\alpha}} + f_{\alpha_i} - g_{\alpha_i} \\ yh_x & y(h_y - m_y) & yh_{\bar{\alpha}} \\ J_{31} & J_{32} & J_{33} \end{bmatrix} \end{aligned} \tag{S30}$$

where subscript variables denote partial derivatives (e.g., $f_x = \partial f/\partial x$) and

$$J_{31} = \overbrace{\epsilon_I(\alpha_{max} - \bar{\alpha})\frac{\partial P_I}{\partial x} - \epsilon_L(\bar{\alpha} - \alpha_{min})\frac{\partial P_L}{\partial x} - (\bar{\alpha} - \alpha_{min})f_x}^{\text{phenotypic plasticity}} \overbrace{-V(\bar{\alpha})g_{x\alpha_i}}^{\text{phenotypic sorting}} \tag{S31}$$

$$J_{32} = \overbrace{\epsilon_I(\alpha_{max} - \bar{\alpha})\frac{\partial P_I}{\partial y} - \epsilon_L(\bar{\alpha} - \alpha_{min})\frac{\partial P_L}{\partial y}}^{\text{phenotypic plasticity}} \overbrace{-V(\bar{\alpha})g_{y\alpha_i}}^{\text{phenotypic sorting}} \tag{S32}$$

$$\begin{aligned} J_{33} = &\overbrace{-\epsilon_I P_I - \epsilon_L P_L + \epsilon_I(\alpha_{max} - \bar{\alpha})\frac{\partial P_I}{\partial \alpha} - \epsilon_L(\bar{\alpha} - \alpha_{min})\frac{\partial P_L}{\partial \alpha}}^{\text{phenotypic plasticity}} \\ &\overbrace{-f - (\bar{\alpha} - \alpha_{min})(f_{\bar{\alpha}} + f_{\alpha_i}) - V(\bar{\alpha})(g_{\bar{\alpha}\alpha_i} + g_{\alpha_i\alpha_i}) - V'(\bar{\alpha})g_{\alpha_i}}^{\text{phenotypic sorting}} \end{aligned} \tag{S33}$$

The signs and interpretations of the entries in the first two rows are identical to those for the Jacobian (S2) in Section S1.1.2. For the entries in the bottom row, the phenotypic plasticity terms are identical to those for the Jacobian (S2) in Section S1.1.2; they define the sensitivities of the induction and loss of induction rates to variation in the state variables. The phenotypic sorting terms account for how variation in the state variables influences the prey reproduction rates (terms involving $f$ and its derivatives) and predation rates (terms involving derivatives of $g$; see below for biological interpretations ). In simulations the phenotypic sorting terms were always much smaller in magnitude than the phenotypic plasticity terms, unless rates of induction were very small ($\epsilon_I$ and $\epsilon_L$ close to zero). For completeness, our analysis discusses how the phenotypic sorting terms affect equilibrium stability and predator-prey phase lags, but we expect the effects of the phenotypic sorting are negligibly small compared to the effects of phenotypic plasticity in most systems.

- $\underline{g_{x\alpha_i}}$: Effect of increased prey density on the benefits of increased defense, where benefits are measured in terms of individual prey fitness. Positive (negative) values imply greater (reduced) benefits of increased individual defense when prey density is higher.

- $\underline{-g_{y\alpha_i}}$: Effect of increased predator density on the benefits of increased defense, where benefits are measured in terms of individual prey fitness. Positive and negative values, respectively, imply greater and reduced benefits of increased individual defense when predator density is higher. Throughout we assume $-g_{y\alpha_i} > 0$. Our reasoning is that it is very difficult to construct models where $-g_{y\alpha_i} < 0$, which suggests that that condition is unlikely to be satisfied in natural systems.

- $\underline{g_{\bar{\alpha}\alpha_i} + g_{\alpha_i\alpha_i}}$: Effect of increased mean defense on the benefits of increased defense, where benefits are measured in terms of individual prey fitness. Positive (negative) values imply greater (reduced) benefits of increased individual defense when mean defense is higher.

##### S1.2.3 Derivation from multimorphic discrete trait model

Here, we show how the continuous trait model (S29) can approximate the dynamics of discrete trait models. We start with a dimorphic model. Then, we use similar calculations to show that a continuous trait model with a similar form can be derived from a discrete trait model with more than 2 phenotypes. As before, the latter derivation is unlikely to be accurate in all cases because it necessarily requires a reduction in model dimension. However, we provide it because it provides additional evidence supporting our claims that the results from our continuous trait model (S29) can provide insight about the dynamics of systems where prey defense levels can take on may different values.

**Approximation of dimorphic model:** Assume an individual prey's defense level can be the values $\alpha_1 = \alpha_{min}$ or $\alpha_2 = \alpha_{max}$. Let $x_i$ be the density of prey with defense level $\alpha_i$. Let $P_1^2$ be the induction rate for individuals of phenotype 1 and $P_2^1$ be the loss of induction rate for individuals with phenotype 2. The dynamics of the prey classes and predator are given by

$$\begin{aligned}
\frac{dx_1}{dt} &= \overbrace{x_1 f(x_1,x_2,\alpha_1) + x_2 f(x_1,x_2,\alpha_2)}^{\text{reproduction}} - \overbrace{x_1 g(x_1,x_2,\alpha_1)}^{\text{predation}} \overbrace{-x_1 P_i^2}^{\text{induction}} \quad \overbrace{+x_2 P_2^1}^{\text{loss of induction}} \\
\frac{dx_2}{dt} &= -\overbrace{x_2 g(x_1,x_2,\alpha_2)}^{\text{predation}} \overbrace{+x_1 P_1^2}^{\text{induction}} \quad \overbrace{-x_2 P_2^1}^{\text{loss of induction}} \\
\frac{dy}{dt} &= \overbrace{y h(x_1,x_2,y)}^{\text{harvesting}} - \overbrace{y m(y)}^{\text{mortality}}
\end{aligned} \tag{S34}$$

where $f(x_1, x_2, \alpha_i)$ and $g(x_1, x_2, y, \alpha_i)$ are the reproduction and predation rates of individuals with phenotype $\alpha_i$. We assume the functional forms of the prey reproduction rates, prey predation rates, and predator harvesting rates depend on total prey density ($x = \sum_i x_i$), predator density (when applicable), the average level of defense ($\bar{\alpha} = \sum_i \alpha_i x_i/x$), and the individual's phenotype ($\alpha_i$, when applicable). Mathematically, we assume $f(x_1, x_2, \alpha_i) = f(x, \bar{\alpha}, \alpha_i)$, $g(x_1, x_2, y, \alpha_i) = g(x, y, \bar{\alpha}, \alpha_i)$, and $h(x_1, x_2, y, ) = h(x, y, \bar{\alpha})$.

We now derive equations that approximate the dynamics of the total density and the average level of defense of the prey population. To help simplify the calculations, we use $q_i = x_i/x$ to denote the frequency of phenotype $i$. The dynamics of total prey density are approximated as

$$\frac{dx}{dt} = \sum_i x_i f(x, \bar{\alpha}, \alpha_i) - x_i g(x, y, \bar{\alpha}, \alpha_i) \tag{S35}$$

$$= x\left[\sum q_i f(x, \bar{\alpha}, \alpha_i) - q_i g(x, \bar{\alpha}, \alpha_i)\right] \tag{S36}$$

$$\approx x\left[f(x, \bar{\alpha}, \bar{\alpha}) - g(x, y, \bar{\alpha}, \bar{\alpha})\right]. \tag{S37}$$

To get the dynamics of the average trait value, we compute

$$\frac{d\bar{\alpha}}{dt} = \frac{1}{x^2}\left[x\sum_i \alpha_i \frac{dx_i}{dt} - \frac{dx}{dt}\sum_i \alpha_i x_i\right] \tag{S38}$$

$$\begin{aligned}
&= \overbrace{\alpha_1 \sum_i q_i f(x, \bar{\alpha}, \alpha_i) - \alpha \sum_i q_i f(x, \bar{\alpha}, \alpha_i) - \sum_i q_i g(x, y, \bar{\alpha}, \alpha_i)(\alpha_i - \bar{\alpha})}^{Q_{sorting},\ \text{phenotypic sorting}} \\
&+ \overbrace{-\alpha_1 q_1 P_1^2 + \alpha_1 q_2 P_2^1 + \alpha_2 q_1 P_1^2 - \alpha_2 q_2 P_2^1}^{Q_{plasticity},\ \text{phenotypic plasticity}}.
\end{aligned} \tag{S39}$$

First consider the phenotypic sorting terms in equation (S12); they account for changes in the mean trait value due to differential reproduction and survival of the different phenotypes. Using $x_1 = x(\alpha_2 - \bar{\alpha})/(\alpha_2 - \alpha_1)$, $x_2 = x(\bar{\alpha} - \alpha_1)/(\alpha_2 - \alpha_1)$, and $q_i = x_i/x$, we get

$$Q_{sorting} = (\alpha_1 - \bar{\alpha})\sum_i q_i f(x, \bar{\alpha}, \alpha_i) + q_1 q_2 (\alpha_2 - \alpha_1)^2 \left[\frac{-g(x, y, \bar{\alpha}, \alpha_2) + g(x, y, \bar{\alpha}, \alpha_1)}{\alpha_2 - \alpha_1}\right] \tag{S40}$$

$$\approx(\alpha_1-\bar{\alpha})f(x,\bar{\alpha},\bar{\alpha})-V(\bar{\alpha})\Big[\frac{\partial g}{\partial \alpha_i}(x,y,\bar{\alpha},\alpha_i)\Big]\Big|_{\alpha_i=\bar{\alpha}} \tag{S41}$$

where $V(\bar{\alpha})=q_1q_2(\alpha_2-\alpha_1)^2=(\alpha_2-\bar{\alpha})(\bar{\alpha}-\alpha_1)$ is the population variance of the trait.

The calculations for the phenotypic plasticity terms in equation (S39) are identical to the calculations in Section S1.1.4. Combining the formulas for $Q_{sorting}$ and $Q_{plasticity}$ and the formulas for the other derivatives yields the continuous trait model (S29).

**Approximation of multimorphic model:** Assume an individual prey's defense level can be $\alpha_i$ ($1\leq i\leq n$) where $\alpha_1=\alpha_{min}$, $\alpha_i<\alpha_{i+1}$, and $\alpha_n=\alpha_{max}$. Let $x_i$ be the density of prey with defense level $\alpha_i$. We assume all individuals are born with trait $\alpha_1$ and individuals can switch from their current defense level to the next higher or next lower level of defense, where $P_i^{i+1}$ is the induction rate at which individuals of phenotype $i$ switch to phenotype $i+1$ and $P_i^{i-1}$ is the loss of induction rate at which individuals of phenotype $i$ switch to phenotype $i-1$.

Under these assumptions, the dynamics of the prey classes and predator are given by

$$\begin{aligned}
\frac{dx_1}{dt}&=\overbrace{\sum_i x_if(x_1,...,x_n,\alpha_i)}^{\text{reproduction}}-\overbrace{x_1g(x_1,...,x_n,\alpha_1)}^{\text{predation}}\ \overbrace{-x_1P_i^2}^{\text{induction}}\ \overbrace{+x_2P_2^1}^{\text{loss of induction}}\\
\frac{dx_i}{dt}&=-\overbrace{x_ig(x_1,...,x_n,\alpha_i)}^{\text{predation}}\ \overbrace{-x_iP_i^{i+1}+x_{i-1}P_{i-1}^i}^{\text{induction}}\ \overbrace{-x_iP_i^{i-1}+x_{i+1}P_{i+1}^i}^{\text{loss of induction}}\\
\frac{dx_n}{dt}&=-\overbrace{x_ng(x_1,...,x_n,\alpha_n)}^{\text{predation}}\ \overbrace{+x_{n-1}P_{n-1}^n}^{\text{induction}}\ \overbrace{-x_nP_n^{n-1}}^{\text{loss of induction}}\\
\frac{dy}{dt}&=\overbrace{yh(x_1,...,x_n,y)}^{\text{harvesting}}-\overbrace{ym(y)}^{\text{mortality}}
\end{aligned} \tag{S42}$$

where $f(x_1,...,x_n,\alpha_i)$ and $g(x_1,...,x_n,y,\alpha_i)$ are the reproduction and predation rates of individuals with phenotype $\alpha_i$. We assume the functional forms of the prey reproduction rates, prey predation rates, and predator harvesting rates depend on total prey density ($x\sum_i x_i$), predator density (when applicable), the average level of defense ($\bar{\alpha}=\sum_i\alpha_ix_i/x$), and the individual's phenotype ($\alpha_i$, when applicable). Mathematically, we assume $f(x_1,...,x_n,\alpha_i)=f(x,\bar{\alpha},\alpha_i)$, $g(x_1,...,x_n,y,\alpha_i)=g(x,y,\bar{\alpha},\alpha_i)$, and $h(x_1,...,x_n,y,)=h(x,y,\bar{\alpha})$.

We now derive equations that approximate the dynamics of the total density and the average level of defense of the prey population; the derivations are similar to those in Section S1.1.4. To help simplify the calculations, we use $q_i=x_i/x$ to denote the frequency of phenotype $i$. The dynamics of total prey density are approximated as

$$\frac{dx}{dt}=\sum_i x_if(x,\bar{\alpha},\alpha_i)-x_ig(x,y,\bar{\alpha},\alpha_i) \tag{S43}$$

$$=x\left[\sum q_if(x,\bar{\alpha},\alpha_i)-q_ig(x,\bar{\alpha},\alpha_i)\right] \tag{S44}$$

$$\approx x\left[f(x,\bar{\alpha},\bar{\alpha}) - g(x,y,\bar{\alpha},\bar{\alpha})\right]. \tag{S45}$$

Following the steps in Section S1.1.4, the dynamics of the mean level of defense are approximated as

$$\frac{d\bar{\alpha}}{dt} = \frac{1}{x^2}\left[x\sum_i \alpha_i \frac{dx_i}{dt} - \frac{dx}{dt}\sum_i \alpha_i x_i\right] \tag{S46}$$

$$\begin{aligned} =& \overbrace{\alpha_1\sum_i q_i f(x,\bar{\alpha},\alpha_i) - \alpha\sum_i q_i f(x,\bar{\alpha},\alpha_i) + \sum_i q_i\left[-g(x,y,\bar{\alpha},\alpha_i)\right](\alpha_i-\bar{\alpha})}^{Q_{sorting},\ \text{phenotypic sorting}} \\ &+ \overbrace{\sum_i \alpha_i(-q_iP_i^{i+1} + q_{i-1}P_{i-1}^{i} - q_iP_i^{i-1} + q_{i+1}P_{i+1}^{i})}^{Q_{plasticity},\ \text{phenotypic plasticity}}. \end{aligned} \tag{S47}$$

$$\approx \overbrace{-(\bar{\alpha}-\alpha_i)f(x,\bar{\alpha},\bar{\alpha}) - Vg_{\alpha_i}(x,y,\bar{\alpha},\bar{\alpha})}^{\text{phenotypic sorting}} + \overbrace{(\alpha_n-\bar{\alpha})\hat{P}_I - (\bar{\alpha}-\alpha_1)\hat{P}_L}^{\text{phenotypic plasticity}} \tag{S48}$$

where $V$ is the population variance of the trait, $\hat{P}_I$ is the average change in the trait values due to induction divided by the average displacement from the maximum trait value, and $\hat{P}_L$ is the average change in the trait values due to loss of induction divided by the average displacement from the minimum trait value. For convenience, we write the average rates of induction and loss of induction in terms of their maximum rates, i.e., $\hat{P}_I = \epsilon_I P_I$ and $\hat{P}_L = \epsilon_L P_L$ where $\epsilon_I = \max P_I$, $\epsilon_L = \max P_L$, and $0 \leq P_I, P_L \leq 1$. This yields the continuous trait model (S29).

#### S1.3 Predator-prey model with a transgenerational inducible defense

For this model, we assume the prey has a transgenerational inducible defense, i.e., the phenotype of an offspring is determined by the parent's environment. In addition, we assume an individual prey's defense value is determined at birth or hatching, depending on rates of induction and loss of induction. The function $P_I$ determines the fraction of offspring that are more induced than the parent and $P_L$ determines the fraction of offspring that are less induced than the parent. In a dimorphic population where individuals are either induced or uninduced, $P_I$ is the fraction of offspring produced by uninduced individuals that are induced and $P_L$ is the fraction of offspring produced by uninduced individuals that are induced; see Section S1.3.3. To facilitate comparisons between the switch at birth and switch after birth models, we abuse terminology and refer to $P_I$ as the induction rate and $P_L$ as the loss of induction rate. This model and our subsequent analysis generalizes the results for the 'switch at birth' model in Appendix E of Cortez (2011).

We assume the mean trait value can take on values in the range $[\alpha_{min}, \alpha_{max}]$ where $\alpha_{min}$ and $\alpha_{max}$ are the minimum and maximum allowable trait values. The switch at birth model with true breeding prey is

$$\frac{dx}{dt} = \overbrace{xf(x,\bar{\alpha},\alpha_i)}^{\text{reproduction}} \quad \overbrace{-xs(x)}^{\text{non-predation mortality}} \quad - \overbrace{xg(x,y,\bar{\alpha},\alpha_i)}^{\text{predation}}\Big|_{\alpha_i=\bar{\alpha}} \tag{S49a}$$

$$\frac{dy}{dt} = \overbrace{yh(x,y,\bar{\alpha})}^{\text{harvesting}} - \overbrace{ym(y)}^{\text{mortality}} \tag{S49b}$$

$$\frac{d\bar{\alpha}}{dt} = \overbrace{[(\alpha_{max}-\bar{\alpha})P_I - (\bar{\alpha}-\alpha_{min})P_L]f}^{\text{phenotypic plasticity}} \overbrace{-V(\bar{\alpha})g_{\alpha_i}\big|_{\alpha_i=\bar{\alpha}}}^{\text{phenotypic sorting}}. \tag{S49c}$$

Prey reproduction rates and non-predation mortality rates are typically combined into one term, but we keep them separate in this model in order to avoid confusion about the effects of induction. All other variables and functions are interpreted as in the switch after birth models.

**Special case of rapid prey turnover:** An important special case to consider is when there is rapid turnover in the prey population. Here, rapid turnover means the prey population has large per capita reproduction rates ($f$) and large non-predation mortality rates ($s$). In this special case, the phenotypic sorting term in the $d\bar{\alpha}/dt$ equation of model (S49) is negligible, which result in the trait equation being approximately $d\bar{\alpha}/dt = [(\alpha_{max} - \bar{\alpha})P_I - (\bar{\alpha} - \alpha_{min})P_L]f$. Importantly, this means that the trait dynamics of the transgenerational inducible defense model (S49) in the rapid turnover limit are identical to the trait dynamics of the intragenerational inducible defense models (S1) and (S29) in the limit of very fast induction and loss of induction rates ($\epsilon_I$ and $\epsilon_L$ large relative to $V(\bar{\alpha})$). Consequently, intragenerational and transgenerational inducible defense have nearly identical dynamics and effects on predator-prey dynamics when induction rates are fast. Said another way, the timing of induction does not influence the effects of inducible defenses on predator-prey dynamics, provided induction and loss of rates are sufficiently fast. Thus, the timing of induction only alters the effects when there is not rapid turnover in the prey population.

The special case of rapid prey turnover is defined mathematically in the following way. We write the prey per capita reproduction rate as $f(x, \bar{\alpha}, \alpha_i) = f_1(x) + \epsilon + f_2(x, \bar{\alpha}, \alpha_i)$ where $f_1 + \epsilon$ are trait independent components of reproduction and $f_2$ is the trait dependent component of reproduction. Then, we write the per capita non-predation mortality rate as $s(x) + \epsilon$. When $\epsilon$ is large in magnitude, there is rapid turnover in the prey population because prey reproduction rates and non-predation mortality are large. In the large $\epsilon$ limit, the trait dynamics are approximately $d\bar{\alpha}/dt = [(\alpha_{max} - \bar{\alpha})P_I - (\bar{\alpha} - \alpha_{min})P_L]f$. Note that the prey density equation is unchanged because the $\epsilon$ terms in $f$ and $s$ cancel out.

##### S1.3.1 Model assumptions

The assumptions about equilibria and the density dynamics are the same as those in Section S1.1.1. The only exception is that the transgenertional inducible defense model (S49) uses a separate function for non-predation mortality of the prey, $s(x)$. We assume the non-predation mortality rate is an increasing function of prey density, i.e., $ds/dx < 0$. For simplicity we assume non-predation mortality is independent of the prey trait, however our results are qualitatively unchanged if non-predation mortality rates are increasing functions of defense (i.e., $\partial s/\partial\bar{\alpha} < 0$ and $\partial s/\partial\alpha_i < 0$).

**Assumptions about trait dynamics:** The trait equation (S49c) decomposes changes in mean defense into phenotypic plasticity and phenotypic sorting components. For the phenotypic plasticity component, we assume the fractions of more induced and less induced offspring are the same for all parents. Thus, an individual's induction level does not influence the distribution of phenotypes of its offspring.

The phenotypic sorting term accounts for changes in the mean trait value due differences in predation rates. The trait variance, $V(\bar{\alpha})$, depends on the mean trait value because the variance necessarily converges to zero as the mean trait value approaches $\alpha_{min}$ or $\alpha_{max}$. The derivative $-g_{\alpha_i} > 0$ accounts for more defended prey having lower predation rates (which causes the mean trait value to shift to larger values).

##### S1.3.2 Jacobian

When evaluated at an equilibrium, $\rho = (x^*, y^*, \alpha^*)$, the Jacobian of model (S49) simplifies to

$$J = \begin{bmatrix} \frac{\partial}{\partial x}\frac{dx}{dt} & \frac{\partial}{\partial y}\frac{dx}{dt} & \frac{\partial}{\partial \bar{\alpha}}\frac{dx}{dt} \\ \frac{\partial}{\partial x}\frac{dy}{dt} & \frac{\partial}{\partial y}\frac{dy}{dt} & \frac{\partial}{\partial \bar{\alpha}}\frac{dy}{dt} \\ \frac{\partial}{\partial x}\frac{d\bar{\alpha}}{dt} & \frac{\partial}{\partial y}\frac{d\bar{\alpha}}{dt} & \frac{\partial}{\partial \bar{\alpha}}\frac{d\bar{\alpha}}{dt} \end{bmatrix} = \begin{bmatrix} J_{11} & J_{12} & J_{13} \\ J_{21} & J_{22} & J_{23} \\ J_{31} & J_{32} & J_{33} \end{bmatrix} \tag{S50}$$

$$= \begin{bmatrix} x(f_x - s_x - g_x) & -xg_x & f_{\bar{\alpha}} - g_{\bar{\alpha}} + f_{\alpha_i} - g_{\alpha_i} \\ yh_x & y(h_y - m_y) & yh_\alpha \\ J_{31} & J_{32} & J_{33} \end{bmatrix} \tag{S51}$$

where subscript variables denote partial derivatives (e.g., $f_x = \partial f/\partial x$) and

$$J_{31} = \overbrace{\left[(\alpha_{max}-\bar{\alpha})\frac{\partial P_I}{\partial x} - (\bar{\alpha}-\alpha_{min})\frac{\partial P_L}{\partial x}\right] f}^{\text{phenotypic plasticity}} + \overbrace{[(\alpha_{max}-\bar{\alpha})P_I - (\bar{\alpha}-\alpha_{min})P_L]f_x}^{\text{transgenerational response}} \overbrace{-V(\bar{\alpha})g_{x\alpha_i}}^{\text{phenotypic sorting}} \tag{S52}$$

$$J_{32} = \overbrace{\left[(\alpha_{max}-\alpha)\frac{\partial P_I}{\partial y} - (\bar{\alpha}-\alpha_{min})\frac{\partial P_L}{\partial y}\right] f}^{\text{phenotypic plasticity}} \overbrace{-V(\bar{\alpha})g_{y\alpha_i}}^{\text{phenotypic sorting}} \tag{S53}$$

$$J_{33} = \overbrace{\left[(\alpha_{max}-\bar{\alpha})\frac{\partial P_I}{\partial \bar{\alpha}} - (\bar{\alpha}-\alpha_{min})\frac{\partial P_L}{\partial \bar{\alpha}}\right] f - [P_I + P_L]f}^{\text{phenotypic plasticity}} + \overbrace{[(\alpha_{max}-\bar{\alpha})P_I - (\bar{\alpha}-\alpha_{min})P_L](f_{\bar{\alpha}} + f_{\alpha_i})}^{\text{transgenerational response}} \overbrace{-V(\bar{\alpha})(g_{\bar{\alpha}\alpha_i} + g_{\alpha_i\alpha_i}) - V'(\bar{\alpha})g_{\alpha_i}}^{\text{phenotypic sorting}} \tag{S54}$$

The signs and assumptions of the entries in the first two rows of the Jacobian are identical to those for Jacobian (S2) in Section S1.1.2.

For the entries in the bottom row, we interpret the terms as effects from phenotypic plasticity (i.e., effects due to the processes of induction and loss of induction), phenotypic sorting (i.e., effects due to differing predation rates on the phenotypes), and transgenerational responses (i.e., effects to the delayed response). The phenotypic plasticity terms in equations (S52)-(S54) are identical to the terms in the Jacobian of the intragenerational models (S1) and (S29) after replacing $f$ with $\epsilon_I = \epsilon_L$. The phenotypic sorting terms in equations (S52)-(S54) are also identical to those in the Jacobian of the intragenerational models (S1) and (S29). The transgenerational response terms in equations (S52) and (S54) are unique to the Jacobian of the transgenerational inducible defense models; this why we associate those terms with the effects of transgenerational responses. Note that the equilibrium condition $d\bar{\alpha}/dt = 0$ implies $[(\alpha_{max} - \bar{\alpha})P_I - (\bar{\alpha} - \alpha_{min})P_L] = Vg_{\alpha_i}/f < 0$. Thus the transgenerational response terms in $J_{31}$ and $J_{33}$ are both positive.

Similar to our previous models, in simulations the phenotypic sorting terms were always smaller in magnitude than the transgenerational response terms and negligibly small relative to the phenotypic plasticity terms, unless prey reproduction rates were very small ($f$ close to zero). For completeness, our analysis discusses how the phenotypic sorting terms affect equilibrium stability and predator-prey phase lags, but we expect the effects of the phenotypic sorting are negligibly small compared to the effects of phenotypic plasticity in most systems. This is because low prey reproduction rates tend to result in the exclusion of the predator or very low predator densities.

- $\underline{g_{x\alpha_i}}$: Effect of increased prey density on the benefits of increased defense, where benefits are measured in terms of individual prey fitness. Positive (negative) values imply greater (reduced) benefits of increased individual defense when prey density is higher.

- $\underline{-g_{y\alpha_i}}$: Effect of increased predator density on the benefits of increased defense, where benefits are measured in terms of individual prey fitness. Positive and negative values, respectively, imply greater and reduced benefits of increased individual defense when predator density is higher. Throughout we assume $-g_{y\alpha_i} > 0$. Our reasoning is that it is very difficult to construct models where $-g_{y\alpha_i} < 0$, which suggests that that condition is unlikely to be satisfied in natural systems.

- $\underline{g_{\bar{\alpha}\alpha_i} + g_{\alpha_i\alpha_i}}$: Effect of increased mean defense on the benefits of increased defense, where benefits are measured in terms of individual prey fitness. Positive (negative) values imply greater (reduced) benefits of increased individual defense when mean defense is higher.

**Special case of rapid prey turnover:** In this special case, the phenotypic sorting component of trait change is negligibly small and the trait dynamics are approximately $d\bar{\alpha}/dt = [(\alpha_{max} - \bar{\alpha})P_I - (\bar{\alpha} - \alpha_{min})P_L)f$. This which means that the coexistence equilibrium satisfies the condition $(\alpha_{max} - \bar{\alpha})\frac{\partial P_I}{\partial x} - (\bar{\alpha} - \alpha_{min})\frac{\partial P_L}{\partial x} \approx 0$. As a consequence, the entries in the bottom row of the Jacobian simplify to

$$J_{31} = [(\alpha_{max} - \bar{\alpha})\frac{\partial P_I}{\partial x} - (\bar{\alpha} - \alpha_{min})\frac{\partial P_L}{\partial x}]f \qquad \text{(S55)}$$

$$J_{32} = [(\alpha_{max} - \bar{\alpha})\frac{\partial P_I}{\partial y} - (\bar{\alpha} - \alpha_{min})\frac{\partial P_L}{\partial y}]f \tag{S56}$$

$$J_{33} = [(\alpha_{max} - \bar{\alpha})\frac{\partial P_I}{\partial \bar{\alpha}} - (\bar{\alpha} - \alpha_{min})\frac{\partial P_L}{\partial \bar{\alpha}}]f - (P_I + P_L)f \tag{S57}$$

Because $f > 0$, these entries have the same signs as the corresponding entries of the Jacobian (S2) of the intragenerational inducible defense model (S1) when $V = 0$.

##### S1.3.3 Derivation from multimorphic discrete trait models

Here, we show how the continuous trait model (S49) can approximately the dynamics of discrete trait models with two phenotypes (generalizing the derivation in Cortez (2011)) and more than two phenotypes. As before, the derivation of the latter is unlikely to be accurate in all cases because it necessarily requires a reduction in model dimension. However, we provide it because it provides additional evidence supporting our claims that the results from our continuous trait model (S49) can provide insight about the dynamics of systems where prey defense levels can take on may different values.

**Derivation from a dimorphic model:** Assume an individual prey's defense level can be the values $\alpha_1 = \alpha_{min}$ or $\alpha_2 = \alpha_{max}$. Let $x_i$ be the density of prey with defense level $\alpha_i$. Let $P_i^j$ be the proportion of offspring with trait $\alpha_j$ produced by an individual with trait $\alpha_i$ and note that $P_i^1 + P_i^2 = 1$.

The dynamics of the prey classes and predator are given by

$$\begin{aligned}
\frac{dx_1}{dt} &= \overbrace{x_1 f(x_1, x_2, \alpha_1)P_1^1 + x_2 f(x_1, x_2, \alpha_2)P_2^1}^{\text{reproduction}} \overbrace{-x_1 s(x_1, x_2)}^{\text{non-predation mortality}} + \overbrace{x_1 g(x_1, x_2, \alpha_1)}^{\text{predation}} \\
\frac{dx_2}{dt} &= \overbrace{x_1 f(x_1, x_2, \alpha_1)P_1^2 + x_2 f(x_1, x_2, \alpha_2)P_2^2}^{\text{reproduction}} \overbrace{-x_2 s(x_1, x_2)}^{\text{non-predation mortality}} - \overbrace{x_2 g(x_1, x_2, \alpha_2)}^{\text{predation}} \\
\frac{dy}{dt} &= \overbrace{yh(x_1, x_2, y)}^{\text{harvesting}} - \overbrace{ym(y)}^{\text{mortality}}
\end{aligned} \tag{S58}$$

where $f(x_1, x_2, \alpha_i)$ and $g(x_1, x_2, y, \alpha_i)$ are the reproduction and predation rates of individuals with phenotype $\alpha_i$. We assume the functional forms of the prey reproduction rates, prey predation rates, and predator harvesting rates depend on total prey density ($x \sum_i x_i$), predator density (when applicable), the average level of defense ($\bar{\alpha} = \sum_i \alpha_i x_i / x$), and the individual's phenotype ($\alpha_i$, when applicable). Mathematically, we assume $f(x_1, x_2, \alpha_i) = f(x, \bar{\alpha}, \alpha_i)$, $g(x_1, x_2, y, \alpha_i) = g(x, y, \bar{\alpha}, \alpha_i)$, and $h(x_1, x_2, y, ) = h(x, y, \bar{\alpha})$. In addition, for simplicity, we assume non-predation mortality only depends on total prey density (i.e., $s(x_1, x_2) = s(x)$), however including trait dependence only results in all instances of $f$ and its derivatives be replaced by $f + s$ and its derivatives. Finally, we assume that the proportions of induced and uninduced offspring are the same for both phenotypes, i.e., the parent's phenotype does not affect the trait distribution of its offspring. Mathematically, this means we assume $P_1^1 = P_2^1 = P_-^1$ and $P_1^2 = P_2^2 = P_-^2$ where $P_-^1 + P_-^2 = 1$.

We now derive equations that approximate the dynamics of the total density and the average level of defense of the prey population. To help simplify the calculations, we use $q_i = x_i/x$ to denote the frequency of phenotype $i$ and we suppress the dependence of the functions on $x$, $y$, and $\bar{\alpha}$, i.e., we use $f(\alpha_i)$ and $g(\alpha_i)$. The dynamics of total prey density are approximated as

$$\frac{dx}{dt} = \sum_i x_i f(\alpha_i) - x_i s(x) - x_i g(\alpha_i) \tag{S59}$$

$$= x\left[\sum_i q_i f(\alpha_i) - q_i s(x) - q_i g(\alpha_i)\right] \tag{S60}$$

$$\approx x\left[f(x,\bar{\alpha},\bar{\alpha}) - s(x) - g(x,y,\bar{\alpha},\bar{\alpha})\right] \tag{S61}$$

where per capita mean reproduction rate, $\sum_i q_i f(x,\bar{\alpha},\alpha_i)$, non-predation mortality rate, $\sum_i q_i s(x)$, and predation rate, $\sum_i q_i g(x,y,\bar{\alpha},\alpha_i)$ are approximated by $f(x,\bar{\alpha},\bar{\alpha})$, $s(x)$, and $g(x,y,\bar{\alpha},\bar{\alpha})$, respectively.

To get the dynamics of the average trait value, we compute

$$\frac{d\bar{\alpha}}{dt} = \frac{1}{x^2}\left[x\sum_i \alpha_i \frac{dx_i}{dt} - \frac{dx}{dt}\sum_i \alpha_i x_i\right] \tag{S62}$$

$$\begin{aligned} =&q_1 f(\alpha_1)P_1^1(\alpha_1-\bar{\alpha}) + q_2 f(\alpha_2)P_2^1(\alpha_1-\bar{\alpha}) + q_1 f(\alpha_1)P_1^2(\alpha_2-\bar{\alpha}) + q_2 f(\alpha_2)P_2^2(\alpha_2-\bar{\alpha}) \\ &+\overbrace{\sum_i \alpha_i q_i s(x) - \bar{\alpha}s(x)}^{\text{zero}} - \sum_i q_i g(\alpha_i)(\alpha_i-\bar{\alpha}) \end{aligned} \tag{S63}$$

$$=[q_1 f(\alpha_1) + q_2 f(\alpha_2)][P_-^1(\alpha_1-\bar{\alpha}) + P_-^2(\alpha_2-\bar{\alpha})] - \sum_i q_i g(\alpha_i)(\alpha_i-\bar{\alpha}) \tag{S64}$$

$$\approx f(x,\bar{\alpha},\bar{\alpha})[P_-^2(\alpha_2-\bar{\alpha}) - P_-^1(\bar{\alpha}-\alpha_1)] - V(\bar{\alpha})g_{\alpha_i} \tag{S65}$$

where $V(\bar{\alpha}) = q_1 q_2(\alpha_2-\alpha_1)^2 = (\alpha_2-\bar{\alpha})(\bar{\alpha}-\alpha_1)$ is the population variance of the trait and the approximations $f(x,\bar{\alpha},\bar{\alpha}) \approx \sum_i q_i f(\alpha_i)$ and $Vg_{\alpha_i} \approx \sum_i q_i g(\alpha_i)(\alpha_i-\bar{\alpha})$ follow from the work in Section S1.1.4. Combining the equations for the total prey density and average prey trait and setting $P_I = P_-^2$, $P_L = P_-^1$, $\alpha_{max} = \alpha_2$ and $\alpha_{min} = \alpha_1$ yields model (S49).

**Derivation from a multimorphic model:** Assume an individual prey's defense level can be $\alpha_i$ $(1 \le i \le n)$ where $\alpha_1 = \alpha_{min}$, $\alpha_i < \alpha_{i+1}$, and $\alpha_n = \alpha_{max}$. Let $x_i$ be the density of prey with defense level $\alpha_i$. Let $P_i^j$ be the proportion of offspring with trait $\alpha_j$ produced by an individual with trait $\alpha_i$ and note that $P_i^1 + P_i^2 = 1$.

The dynamics of the prey classes and predator are

$$\begin{aligned} \frac{dx_i}{dt} &= \overbrace{\sum_j x_j f(x_1,..,x_n,\alpha_j)P_j^i}^{\text{reproduction}} \overbrace{-x_i s(x_1,...,x_n)}^{\text{non-predation mortality}} + \overbrace{x_i g(x_1,...,x_n,\alpha_i)}^{\text{predation}} \\ \frac{dy}{dt} &= \overbrace{yh(x_1,...,x_n,y)}^{\text{harvesting}} - \overbrace{ym(y)}^{\text{mortality}} \end{aligned} \tag{S66}$$

where $f(x_1,...,x_n,\alpha_i)$ and $g(x_1,...,x_n,y,\alpha_i)$ are the reproduction and predation rates of individuals with phenotype $\alpha_i$. We assume the functional forms of the prey reproduction rates, prey predation rates, and predator harvesting rates depend on total prey density ($x\sum_i x_i$), predator density (when applicable), the average level of defense ($\bar{\alpha}=\sum_i \alpha_i x_i/x$), and the individual's phenotype ($\alpha_i$, when applicable). Mathematically, we assume $f(x_1,...,x_n,\alpha_i)=f(x,\bar{\alpha},\alpha_i)$, $g(x_1,...,x_n,y,\alpha_i)=g(x,y,\bar{\alpha},\alpha_i)$, and $h(x_1,...,x_n,y,)=h(x,y,\bar{\alpha})$. In addition, for simplicity, we assume non-predation mortality only depends on total prey density (i.e., $s(x_1,...,x_n)=s(x)$), however including trait dependence only results in all instances of $f$ and its derivatives be replaced by $f+s$ and its derivatives. Finally, we assume that the proportions of induced and uninduced offspring are the same for both phenotypes, i.e., the parent's phenotype does not affect the trait distribution of its offspring. Mathematically, this means we assume $P_i^1=P_-^1$ and $P_i^2=P_-^2$ for all $i$ where $\sum_j P_-^j=1$.

We now derive equations that approximate the dynamics of the total density and the average level of defense of the prey population. To help simplify the calculations, we use $q_i=x_i/x$ to denote the frequency of phenotype $i$ and we suppress the dependence of the functions on $x$, $y$, and $\bar{\alpha}$. The dynamics of total prey density are approximated as

$$\frac{dx}{dt}=\sum_i x_i f(\alpha_i)-x_i s(x)-x_i g(\alpha_i) \tag{S67}$$

$$=x\left[\sum_i q_i f(\alpha_i)-q_i s(x)-q_i g(\alpha_i)\right] \tag{S68}$$

$$\approx x\left[f(x,\bar{\alpha},\bar{\alpha})-s(x)-g(x,y,\bar{\alpha},\bar{\alpha})\right] \tag{S69}$$

where per capita mean reproduction rate, $\sum_i q_i f(x,\bar{\alpha},\alpha_i)$, non-predation mortality rate, $\sum_i q_i s(x)$, and predation rate, $\sum_i q_i g(x,y,\bar{\alpha},\alpha_i)$ are approximated by $f(x,\bar{\alpha},\bar{\alpha})$, $s(x)$, and $g(x,y,\bar{\alpha},\bar{\alpha})$, respectively.

To get the dynamics of the average trait value, we compute

$$\frac{d\bar{\alpha}}{dt}=\frac{1}{x^2}\left[x\sum_i \alpha_i\frac{dx_i}{dt}-\frac{dx}{dt}\sum_i \alpha_i x_i\right] \tag{S70}$$

$$=\sum_j \alpha_j\sum_i q_i f_i P_-^j-\bar{\alpha}\sum_i q_i f_i-\sum_i q_i g(\alpha_i)(\alpha_i-\bar{\alpha}) \tag{S71}$$

$$=\left[\sum_i q_i f_i\right]\sum_j(\alpha_j-\bar{\alpha})P_-^j-\sum_i q_i g(\alpha_i)(\alpha_i-\bar{\alpha}) \tag{S72}$$

$$=\left[\sum_i q_i f_i\right]\sum_j(\alpha_j-\bar{\alpha})P_-^j\left[\sum_i q_i\right]-\sum_i q_i g(\alpha_i)(\alpha_i-\bar{\alpha}) \tag{S73}$$

$$=\left[\sum_i q_i f_i\right]\left[\sum_{i\neq j}(\alpha_j-\bar{\alpha})P_-^j q_i+\sum_j(\alpha_j-\bar{\alpha})P_-^j q_j\right]-\sum_i q_i g(\alpha_i)(\alpha_i-\bar{\alpha}) \tag{S74}$$

$$=\left[\sum_i q_i f_i\right]\left[\sum_{i\neq j}(\alpha_j-\bar{\alpha})P_-^j q_i+\sum_j(\alpha_j-\bar{\alpha})\big(1-\sum_{i\neq j}P_-^j\big)q_j\right]-\sum_i q_i g(\alpha_i)(\alpha_i-\bar{\alpha}) \tag{S75}$$

$$=\left[\sum_i q_i f_i\right]\left[\sum_{i\neq j}(\alpha_j-\alpha_i)P_-^j q_i+\sum_j(\alpha_j-\bar{\alpha})q_j\right]-\sum_i q_i g(\alpha_i)(\alpha_i-\bar{\alpha}) \tag{S76}$$

$$= \left[\sum_i q_i f_i\right] \left[ \overbrace{(\alpha_n - \bar{\alpha}) \frac{\sum_{i<j} (\alpha_j - \alpha_i) P_-^j q_i}{\sum_{j \neq n} (\alpha_n - \alpha_j) q_j}}^{\text{offspring more induced than parents}} + \overbrace{(\bar{\alpha} - \alpha_1) \frac{\sum_{i>j} (\alpha_j - \alpha_i) P_-^j q_i}{\sum_{j \neq 1} (\alpha_j - \alpha_1) q_j}}^{\text{offspring less induced than parents}} \right] - \sum_i q_i g(\alpha_i)(\alpha_i - \bar{\alpha}) \tag{S77}$$

$$\approx f(x, \bar{\alpha}, \bar{\alpha})[P_I(\alpha_n - \bar{\alpha}) - P_L(\bar{\alpha} - \alpha_1)] - V g_{\alpha_i}. \tag{S78}$$

The second to last line was simplified using the equality $\sum_j (\alpha_j - \bar{\alpha}) q_j = 0$. In addition, reproduction from individuals were split into terms accounting for offspring that are more induced than their parents and offspring that are less induced than their parents. To get an equation that is similar in form to equation (S27), we also used equality (S22) from Section S1.1.4. The final line follow from the work in Section S1.1.4. Specifically, $V = \sum_{i \neq j} (\alpha_i - \alpha_j)^2$ is the population variance of the trait and we use the approximations $f(x, \bar{\alpha}, \bar{\alpha}) \approx \sum_i q_i f(\alpha_i)$ and $V g_{\alpha_i} \approx \sum_i q_i g(\alpha_i)(\alpha_i - \bar{\alpha})$. The average induction rate, $P_I$, is the average change in the trait values due to induction ($\sum_{i \neq n} q_i (\alpha_{i+1} - \alpha_i) P_i^{i+1}$) divided by the average displacement from the maximum trait value ( $\sum_{j \neq n} (\alpha_n - \alpha_j) q_j$). When multiplied by $\alpha_n - \bar{\alpha}$, this yields the average change in the mean trait value due to induction. Similarly, the average loss of induction rate, $P_L$, is the average change in the trait due to loss of induction ($\sum_{i \neq n} q_i (\alpha_{i+1} - \alpha_i) P_{i+1}^i$) divided by the average displacement from the minimum trait value ($\sum_{j \neq 1} (\alpha_j - \alpha_1) q_j$). When multiplied by $\bar{\alpha} - \alpha_1$, this yields the average change in the mean trait value due to loss of induction. Combining the equations for the total prey density and average prey trait and setting $\alpha_{max} = \alpha_n$ and $\alpha_{min} = \alpha_1$ yields model (S49).

#### S1.4 Classes of inducible defenses with different stimuli

We consider four classes of inducible defenses that differ based on the stimuli affecting the induction and loss of induction rates, $P_I$ and $P_L$. The signs of the derivatives of $P_I$ for each class are summarized in Table S1; the signs of $P_L$ are the opposite. Interpretations for the quantities in the last column are given in section S1.1.2.

**Class 1: Induction stimulus is predator density, $P_I(y)$**
We assume the induction and loss of induction rates only depend on predator density. This corresponds to inducible defenses driven by a cue (e.g., karimones) released by the predator. Mathematically, we assume the induction rate increases with predator density ($\partial P_I / \partial y > 0$) and is independent of prey density and mean defense. Similarly, we assume the loss of induction rate decreases with predator density ($\partial P_L / \partial y < 0$) and is independent of prey density and mean defense.

**Table S1:** Signs of the derivatives of the induction rate, $P_i$, for the four classes of inducible defenses

| Derivative | Class 1 $P_I(y)$ | Class 2 $P_i(x,y)$ | Class 3 $P_I\,(xg)$ | Class 4 $P_i\,(f_{\alpha_i} - g_{\alpha_i})$ |
|---|---|---|---|---|
| $\dfrac{\partial P_I}{\partial x}$ | 0 | $-$ | $+$ | $f_{x\alpha_i} - g_{x\alpha_i}$ |
| $\dfrac{\partial P_I}{\partial y}$ | $+$ | $+$ | $+$ | $-g_{y\alpha_i}$ |
| $\dfrac{\partial P_I}{\partial \alpha}$ | 0 | 0 | $-$ | $f_{\alpha_i\bar{\alpha}} - g_{\alpha_i\bar{\alpha}} + f_{\alpha_i\alpha_i} - g_{\alpha_i\alpha_i}$ |

The signs of the derivatives of $P_L$ are the opposite of the signs of $P_i$.

**Class 2: Induction stimuli are prey and predator densities,** $P_I(x, y)$
We assume the induction and loss of induction rates depend on prey and predator densities. This corresponds to inducible defenses driven by a cue (e.g., karimones) released by the predator and a cue released by conspecifics. Mathematically, we assume the induction rate decreases with prey density ($\partial P_I/\partial x < 0$), increases with predator density ($\partial P_I/\partial y > 0$), and is independent of mean defense. Similarly, we assume the loss of induction rate increases with prey density ($\partial P_L/\partial x > 0$), decreases with predator density ($\partial P_L/\partial x < 0$), and is independent of mean defense.

**Class 3: Induction stimulus is predation rate,** $P_I(xg(x, y, \alpha))$
We assume the induction and loss of induction rates depend on the rate of predation of prey ($xg$). This corresponds to inducible defenses driven by a cue related to conspecific mortality due to predation, e.g., the entrails of conspecifics that have been released into the environment by messy consumers. Mathematically, we assume the induction rate increases and the loss of induction rate decreases with increasing predation rates ($xg$). Because predation rates increase with increased prey and predator densities and decrease with increased defense, the induction rate is an increasing function of prey and predator density ($\partial P_I/\partial x > 0$) and a decreasing function of defense ($\partial P_I/\partial \alpha < 0$), and the loss of induction rate is a decreasing function of prey and predator density ($\partial P_I/\partial y > 0$) and an increasing function of defense ($\partial P_I/\partial \bar{\alpha} > 0$).

**Class 4: Induction stimulus is the fitness gradient,** $P_I(f_{\alpha_i} - g_{\alpha_i})$
We assume the induction and loss of induction rates depend on the fitness gradient ($f_{\alpha_i} - g_{\alpha_i}$). This corresponds to inducible defenses driven by a collection of cues that allow individuals to accurately assess individual fitness. Mathematically, we assume $P_I$ is an increasing function of its argument ($P_I' > 0$) and $P_L$ is a decreasing function of its argument ($P_L' < 0$). This means that the partial derivatives of $P_I$ satisfy $\partial P_I/\partial x = C_1(x, y, \alpha)(f_{x\alpha_i} - g_{x\alpha_i})$, $\partial P_I/\partial y = C_2(x, y, \alpha)(-g_{x\alpha_i})$, and $\partial P_I/\partial \alpha = C_3(x, y, \alpha)(f_{\alpha_i\bar{\alpha}} - g_{\alpha_i\bar{\alpha}} + f_{\alpha_i\alpha_i} - g_{\alpha_i\alpha_i})$ where the $C_i$ are positive functions. Similarly, the partial derivatives of $P_L$ satisfy $\partial P_L/\partial x = -C_4(x, y, \alpha)(f_{x\alpha_i} - g_{x\alpha_i})$, $\partial P_L/\partial y = -C_5(x, y, \alpha)(-g_{x\alpha_i})$, and $\partial P_L/\partial \alpha = -C_6(x, y, \alpha)(f_{\alpha_i\bar{\alpha}} - g_{\alpha_i\bar{\alpha}} + f_{\alpha_i\alpha_i} - g_{\alpha_i\alpha_i})$ where the $C_i$ are positive functions. Biological interpretations for the derivatives of $f_{\alpha_i} - g_{\alpha_i}$ are given in Section S1.1.2.

#### S1.5 Effects of inducible defenses on equilibrium stability

We use the Routh-Hurwitz criteria to explore how inducible defenses affect equilibrium stability. The characteristic polynomial of the Jacobian (S1.1.2) is

$$\rho(\lambda) = \lambda^3 + a_1\lambda^2 + a_2\lambda + a_3 \tag{S79}$$

where

$$a_1 = -\,\mathrm{trace}(J) = -J_{11} - J_{22} - J_{33} \tag{S80}$$

$$a_2 = \begin{vmatrix} J_{11} & J_{12} \\ J_{21} & J_{22} \end{vmatrix} + \begin{vmatrix} J_{11} & J_{13} \\ J_{31} & J_{33} \end{vmatrix} + \begin{vmatrix} J_{22} & J_{23} \\ J_{32} & J_{33} \end{vmatrix} \tag{S81}$$

$$a_3 = -\,|J| \tag{S82}$$

The Routh-Hurwitz criteria for stability implies that an equilibrium is locally asymptotically stable if $a_j > 0$ for $1 \le j \le 3$ and $a_1a_2 - a_3 > 0$ where

$$\begin{aligned} a_1a_2 - a_3 =& (-J_{11} - J_{22}) \begin{vmatrix} J_{11} & J_{12} \\ J_{21} & J_{22} \end{vmatrix} + J_{33}^2(-J_{11} - J_{22}) \\ &+ \underbrace{\overbrace{J_{33}J_{13}J_{31}}^{J_{33}J_{13}J_{31}} + \overbrace{J_{33}J_{23}J_{32}}^{-J_{33}J_{32}} + \overbrace{-J_{33}(J_{11}+J_{22})^2}^{-J_{33}} + \overbrace{J_{31}J_{11}J_{13}}^{J_{31}J_{11}J_{13}} + \overbrace{J_{31}J_{12}J_{23}}^{J_{31}} + \overbrace{J_{32}J_{22}J_{23}}^{J_{32}} + \overbrace{J_{32}J_{13}J_{21}}^{J_{32}J_{13}}}_{\text{effect of inducible defense}} \end{aligned} \tag{S83}$$

The sign of each term in the second line of equation (S83) is listed above its overbrace. The terms on the first line of equation (S83) are $\mathcal{O}(1)$, the first two terms on the second of equation (S83) are $\mathcal{O}(\epsilon_I^2, \epsilon_I\epsilon_L, \epsilon_L^2)$, and the remaining terms are $\mathcal{O}(\epsilon_I, \epsilon_L)$.

Our analysis focuses on the quantity $a_1a_2 - a_3$, which determines if the equilibrium is stable (positive), undergoing a Hopf bifurcation (zero), or unstable (negative), provided $a_j > 0$ for $1 \le j \le 3$. The first line of equation (S83) determines how the dynamics of the prey and predator densities affect equilibrium stability. We assume $J_{11}J_{22} - J_{21}J_{12} > 0$, which means the two species can coexist when prey defense is a fixed quantity. Thus, the dynamics of the densities have a stabilizing effect when $J_{11} + J_{22} < 0$ and a destabilizing effect when $J_{11} + J_{22} > 0$. The second line of equation (S83) determines how the dynamics of prey defense influence system stability. Defenses is stabilizing and destabilizing when it makes $a_1a_2 - a_3$ more positive and more negative, respectively.

In the following subsections, we first show how the phenotypic plasticity components of trait change affect equilibrium stability for each class of inducible defense. We do this by assuming the phenotypic sorting components of trait change in all models are negligible (mathematically, we set $V = 0$). We discuss our results in terms of the maximum rates of induction ($\epsilon_I$) and loss of induction ($\epsilon_L$) in the switch after birth models (S1) and (S29), but all of our result also apply to the switch at birth model (S49) after replacing $\epsilon_I$ and $\epsilon_L$ with $f$; see Section S1.3.2. The remaining subsections address how the predictions are altered by the phenotypic sorting terms for each model, irreversibility, and transgenerational responses.

##### S1.5.1 Class 1: Induction stimulus is predator density, $P_i(y)$

Assume $V = 0$, $\partial P_I/\partial x = 0$, $\partial P_I/\partial y > 0$, and $\partial P_I/\partial \alpha = 0$. This implies $J_{31} = 0$, $J_{32} > 0$, and $J_{33} < 0$.

The effects of the inducible defense on equation (S83) are determined by the terms,

$$\underbrace{\overbrace{J_{33}J_{23}J_{32}}^{\text{positive}}}_{\mathcal{O}(\epsilon_I^2,\epsilon_I\epsilon_L,\epsilon_L^2)} + \underbrace{\overbrace{-J_{33}(J_{11}+J_{22})^2 + J_{32}J_{22}J_{23}}^{\text{positive}} + \overbrace{J_{32}J_{13}J_{21}}^{\text{sign of } J_{13}}}_{\mathcal{O}(\epsilon_I,\epsilon_L)}. \tag{S84}$$

The $\mathcal{O}(\epsilon_I^2, \epsilon_I\epsilon_L, \epsilon_L^2)$ terms are always positive. The $\mathcal{O}(\epsilon_I, \epsilon_L)$ terms are positive unless $J_{13}$ is negative and sufficiently large in magnitude.

Interpretation: The phenotypic plasticity component of trait change is always stabilizing if induction and loss of induction rates are sufficiently fast ($\epsilon_I$ and $\epsilon_L$ large). The phenotypic plasticity component of trait change is typically stabilizing for slower rates of induction and loss of induction, but it can be destabilizing when increased mean defense decreases individual prey fitness ($f_{\bar{\alpha}} - g_{\bar{\alpha}} + f_{\alpha_i} - g_{\alpha_i} < 0$ and large).

##### S1.5.2 Class 2: Induction stimuli are prey and predator densities, $P_i(x, y)$

Assume $V = 0$, $\partial P_I/\partial x < 0$, $\partial P_I/\partial y > 0$, and $\partial P_I/\partial \alpha = 0$. This implies $J_{31} < 0$, $J_{32} > 0$, and $J_{33} < 0$.

The effects of the inducible defense on equation (S83) are determined by the terms,

$$\underbrace{\overbrace{J_{33}J_{13}J_{31}}^{\text{sign of } J_{13}} + \overbrace{J_{33}J_{23}J_{32}}^{\text{positive}}}_{\mathcal{O}(\epsilon_I^2,\epsilon_I\epsilon_L,\epsilon_L^2)} + \underbrace{\overbrace{-J_{33}(J_{11}+J_{22})^2 + J_{32}J_{22}J_{23}}^{\text{positive}} + \overbrace{J_{31}J_{12}J_{23}}^{\text{negative}} + \overbrace{J_{13}J_{32}J_{21}}^{\text{sign of } J_{13}} + \overbrace{J_{13}J_{31}J_{11}}^{\text{sign of } -J_{11}J_{13}}}_{\mathcal{O}(\epsilon_I,\epsilon_L)}. \tag{S85}$$

The terms involving $J_{31}$ define the effects of induction and loss of induction depending on prey density. The $\mathcal{O}(\epsilon_I^2, \epsilon_I\epsilon_L, \epsilon_L^2)$ terms of equation (S85) are positive unless $J_{13}$ is negative and $J_{13}J_{31}$ is large in magnitude. The $\mathcal{O}(\epsilon_I, \epsilon_L)$ terms of equation (S85) are typically positive. However, they can be negative when $J_{31}$ is sufficiently large in magnitude or when $J_{13}$ is negative and sufficiently large in magnitude.

Interpretation: The phenotypic plasticity component of trait change is typically stabilizing. In particular, when induction and loss of induction rates are very fast ($\epsilon_I$ and $\epsilon_L$ large), the phenotypic plasticity component of trait change is stabilizing unless (i) increased mean defense decreases individual prey fitness ($f_{\bar{\alpha}} - g_{\bar{\alpha}} + f_{\alpha_i} - g_{\alpha_i} < 0$ and large). When rates are not as fast, the phenotypic plasticity component of trait change is stabilizing unless (i) increased mean defense decreases individual prey fitness ($f_{\bar{\alpha}} - g_{\bar{\alpha}} + f_{\alpha_i} - g_{\alpha_i} < 0$ and large) or (ii) the rates of induction and loss of induction are highly sensitive to changes in prey density ($\partial P_I/\partial x$ and $\partial P_L/\partial x$ large in magnitude).

##### S1.5.3 Class 3: Induction stimulus is the predation rate, $P_i(xg)$

Assume $V = 0$, $\partial P_I/\partial x > 0$, $\partial P_I/\partial y > 0$, and $\partial P_I/\partial \alpha < 0$. This implies $J_{31} > 0$, $J_{32} > 0$, and $J_{33} < 0$.

The effects of inducible defenses on equation (S83) are determined by the terms,

$$\underbrace{\overbrace{J_{33}J_{13}J_{31}}^{\text{sign of } -J_{13}} + \overbrace{J_{33}J_{23}J_{32}}^{\text{positive}}}_{\mathcal{O}(\epsilon_I^2,\epsilon_I\epsilon_L,\epsilon_L^2)} + \underbrace{\overbrace{-J_{33}(J_{11}+J_{22})^2 + J_{32}J_{22}J_{23} + J_{31}J_{12}J_{23}}^{\text{positive}} + \overbrace{J_{13}J_{32}J_{21}}^{\text{sign of } J_{13}} + \overbrace{J_{13}J_{31}J_{11}}^{\text{sign of } J_{11}J_{13}}}_{\mathcal{O}(\epsilon_I,\epsilon_L)} \tag{S86}$$

$$= \underbrace{\overbrace{J_{33}J_{13}J_{31}}^{\text{sign of } -J_{13}} + \overbrace{J_{33}J_{23}J_{32}}^{\text{positive}}}_{\mathcal{O}(\epsilon_I^2,\epsilon_I\epsilon_L,\epsilon_L^2)}$$
$$+ \underbrace{\overbrace{-J_{33}(J_{11}+J_{22})^2 + J_{32}J_{22}J_{23} + J_{31}J_{12}J_{23}}^{\text{positive}} + \overbrace{J_{13}(J_{32}J_{21} - J_{31}J_{22})}^{\text{sign of } J_{13}} + \overbrace{J_{13}J_{31}(J_{11}+J_{22})}^{\text{sign of } J_{13}(J_{11}+J_{22})}}_{\mathcal{O}(\epsilon_I,\epsilon_L)}. \tag{S87}$$

The $\mathcal{O}(\epsilon_I^2, \epsilon_I\epsilon_L, \epsilon_L^2)$ terms in equation (S86) are positive unless $J_{13}$ is positive and large in magnitude. The $\mathcal{O}(\epsilon_I, \epsilon_L)$ terms in equation (S86) are typically positive. However, they can be negative if $J_{13}$ is negative and large or if $J_{11} < 0$ and $J_{13}$ is positive and large. Equation (S86) is an altered form where $J_{13}J_{31}J_{22}$ is added and subtracted to the equation; its form emphasizes that the $\mathcal{O}(\epsilon_I, \epsilon_L)$ terms can be negative when $J_{13} > 0$, only if the ecological subsystems is strongly stable ($J_{11} + J_{22} < 0$ and large in magnitude).

Interpretation: When induction and loss of induction rates are very fast, the phenotypic plasticity component of trait change is stabilizing unless (i) the benefits of defense are sufficiently high such that increases in mean levels of defense cause large increases in per capita growth rates ($\partial f/\partial \alpha > 0$ and large). When rates are not as fast, the phenotypic plasticity component of trait change is stabilizing unless (ii) increased mean defense decreases individual prey fitness ($f_{\bar{\alpha}} - g_{\bar{\alpha}} + f_{\alpha_i} - g_{\alpha_i} < 0$ and large) or (ii) prey are not overexploited by predators ($J_{11} < 0$), increased mean defense increases individual prey fitness ($f_{\bar{\alpha}} - g_{\bar{\alpha}} + f_{\alpha_i} - g_{\alpha_i} > 0$ and large), and induction and loss of induction rates are highly sensitive to changes in prey density ($\partial P_I/\partial x$ and $\partial P_L/\partial x$ large in magnitude). The first part of the second condition can also be stated as the density dynamics are strongly stabilizing ($J_{11} + J_{22} < 0$ and large).

##### S1.5.4 Class 4: Induction stimulus is the prey fitness gradient, $P_i(\frac{\partial}{\partial \alpha_i}\frac{1}{x}\frac{dx}{dt})$

Assume $V = 0$. Assume the signs of $\partial P_I/\partial x$, $\partial P_I/\partial y$, and $\partial P_I/\partial \alpha$ are the same as $f_{x\alpha_i} - g_{x\alpha_i}$, $f_{y\alpha_i} - g_{y\alpha_i}$, and $f_{\alpha_i\bar{\alpha}} - g_{\alpha_i\bar{\alpha}} + f_{\alpha_i\alpha_i} - g_{\alpha_i\alpha_i}$, respectively, and assume the signs of $\partial P_L/\partial x$, $\partial P_L/\partial y$, and $\partial P_L/\partial \alpha$ are the opposite. This implies $J_{31} = \epsilon C_{31}(f_{x\alpha_i} - g_{x\alpha_i})$, $J_{32} = \epsilon C_{32}(f_{y\alpha_i} - g_{y\alpha_i})$, and $J_{33} = -\epsilon P_I - \epsilon P_L + \epsilon C_{33}(f_{\alpha_i\bar{\alpha}} - g_{\alpha_i\bar{\alpha}} + f_{\alpha_i\alpha_i} - g_{\alpha_i\alpha_i})$ where the $C_{ij}$ are positive.

The effects of phenotypic plasticity on equation (S83) are determined by the terms,

$$\begin{aligned} &\underbrace{\overbrace{J_{33}J_{13}J_{31}}^{(f_{\alpha_i\alpha_i}-g_{\alpha_i\alpha_i}+f_{\alpha_i\bar{\alpha}}-g_{\alpha_i\bar{\alpha}})J_{13}(f_{x\alpha_i}-g_{x\alpha_i})} + \overbrace{J_{33}J_{23}J_{32}}^{-(f_{\alpha_i\alpha_i}-g_{\alpha_i\alpha_i}+f_{\alpha_i\bar{\alpha}}-g_{\alpha_i\bar{\alpha}})(-g_{y\alpha_i})}}_{\mathcal{O}(\epsilon_I^2,\epsilon_I\epsilon_L,\epsilon_L^2)} + \\ &+ \underbrace{\overbrace{-J_{33}(J_{11}+J_{22})^2}^{-(f_{\alpha_i\alpha_i}-g_{\alpha_i\alpha_i}+f_{\alpha_i\bar{\alpha}}-g_{\alpha_i\bar{\alpha}})} + \overbrace{J_{31}J_{11}J_{13}}^{J_{11}J_{13}(f_{x\alpha_i}-g_{x\alpha_i})} + \overbrace{J_{31}J_{12}J_{23}}^{f_{x\alpha_i}-g_{x\alpha_i}} + \overbrace{J_{32}J_{22}J_{23}}^{-g_{y\alpha_i}} + \overbrace{J_{32}J_{13}J_{21}}^{-g_{y\alpha_i}J_{13}}}_{\mathcal{O}(\epsilon_I,\epsilon_L)} \end{aligned} \tag{S88}$$

where the sign of each term is shown above the overbrace.

First, consider systems where $J_{33} > 0$, i.e., $f_{\alpha_i\alpha_i} - g_{\alpha_i\alpha_i} + f_{\alpha_i\bar{\alpha}} - g_{\alpha_i\bar{\alpha}} > 0$. The equilibrium will be unstable for sufficiently large $\epsilon_I, \epsilon_L$ because $a_1$ in equation (S80) will be negative. This means that when $f_{\alpha_i\alpha_i} - g_{\alpha_i\alpha_i} + f_{\alpha_i\bar{\alpha}} - g_{\alpha_i\bar{\alpha}} > 0$ the inducible defense typically has a destabilizing effect on the equilibrium. The only exception is when $f_{\alpha_i\alpha_i} - g_{\alpha_i\alpha_i} + f_{\alpha_i\bar{\alpha}} - g_{\alpha_i\bar{\alpha}}$ is positive but very small in magnitude such that $J_{33}$ is negative. In this very rare case, the effects on stability are the same as the next case.

Second, now consider systems where $J_{33} < 0$, which typically implies $f_{\alpha_i\alpha_i} - g_{\alpha_i\alpha_i} + f_{\alpha_i\bar{\alpha}} - g_{\alpha_i\bar{\alpha}} < 0$. The $\mathcal{O}(\epsilon_I^2, \epsilon_I\epsilon_L, \epsilon_L^2)$ terms of equation (S88) are positive unless $J_{13}J_{31} > 0$ or $J_{32} < 0$. The $\mathcal{O}(\epsilon_I, \epsilon_L)$ terms are positive unless $J_{31} < 0$, $J_{32} < 0$, $J_{13}J_{32} < 0$, or $J_{11}J_{13}J_{31} < 0$. Because $J_{32} < 0$ is unlikely to arise, combining the above yields that equation (S88) will be positive when $J_{33} < 0$ unless $J_{31} < 0$ or $J_{13} < 0$ are sufficiently large in magnitude.

Interpretation: First, consider systems where individual fitness is maximized by extreme trait values (i.e., $f_{\alpha_i\alpha_i} - g_{\alpha_i\alpha_i} + f_{\alpha_i\bar{\alpha}} - g_{\alpha_i\bar{\alpha}} > 0$). In this case, inducible defenses are expected to be destabilizing. In the special case where $-\epsilon(P_I + P_L)$ is larger in magnitude than $C_{33}\epsilon(f_{\alpha_i\bar{\alpha}} - g_{\alpha_i\bar{\alpha}} + f_{\alpha_i\alpha_i} - g_{\alpha_i\alpha_i})$, the sign of $J_{33}$ will be negative. Biologically, this only occurs when induction and loss of induction rates are weakly sensitive to changes in prey defense meaning (a) changes in mean defense have small effects on prey fitness (i.e., the local fitness landscape is relatively flat; $f_{\alpha_i\bar{\alpha}} - g_{\alpha_i\bar{\alpha}} + f_{\alpha_i\alpha_i} - g_{\alpha_i\alpha_i}$ is small in magnitude) or (b) large changes in mean defense only cause small changes in the induction rate (i.e., $\partial P_I/\partial\alpha$ and $\partial P_L/\partial\alpha$ small in magnitude). Because the mathematical conditions defining this special case are restrictive, we expect it to be rare in nature.

Second, consider systems where individual fitness is maximized by intermediate trait values (i.e., $f_{\alpha_i\alpha_i} - g_{\alpha_i\alpha_i} + f_{\alpha_i\bar{\alpha}} - g_{\alpha_i\bar{\alpha}} < 0$). In this case, inducible defenses are typically stabilizing. However, they can be destabilizing if (i) the benefits of increased defense decrease as prey density decreases ($f_{x\alpha_i} - g_{x\alpha_i} < 0$) or (ii) prey fitness decreases with increased average defense ($J_{13} < 0$). The inducible defenses can also be destabilizing when (iii) the benefits of increased defense decrease as predator density increases ($-g_{y\alpha_i} < 0$). However, we expect condition (ii) is rarely satisfied in natural systems for two reasons. First, our assumptions that the predation rate increases with predator density ($\partial g/\partial y > 0$) and decreases with increased defense ($\partial g/\partial\alpha$) make it less likely that the function will also satisfy $-g_{y\alpha_i} < 0$. Second, in practice, it is difficult to construct biologically reasonable functional responses that satisfy all three of the inequalities in the previous sentence.

##### S1.5.5 Effects of phenotypic sorting in the intragenerational inducible defense model with true breeding prey

The effects of phenotypic sorting in model (S1) are determined by the following terms in the Jacobian entries

$$\begin{aligned} J_{31}: &\quad V(\bar{\alpha})(f_{x\alpha_i} - g_{x\alpha_i}) \\ J_{32}: &\quad -V(\bar{\alpha})g_{y\alpha_i} \\ J_{33}: &\quad V(\bar{\alpha})(f_{\alpha_i\alpha_i} - g_{\alpha_i\alpha_i} + f_{\alpha_i\bar{\alpha}} - g_{\alpha_i\bar{\alpha}}) + V'(\bar{\alpha})(f_{\alpha_i} - g_{\alpha_i}) \end{aligned} \tag{S89}$$

For $J_{31}$, if $J_{33}$ is negative, then the phenotypic sorting terms typically have a destabilizing effect when they make $J_{31}$ more negative. However, for Class 2 inducible defenses, they can have a destabilizing effect if they make $J_{31}$ more positive. For $J_{32}$, we expect the phenotypic sorting terms have a stabilizing effect because we expect $-Vg_{y\alpha_i} > 0$ in most systems; see the end of Section S1.5.4 for details. For $J_{33}$, the phenotypic sorting terms have a destabilizing effect when they make $J_{33}$ more positive.

Interpretation: In general, the effects of phenotypic sorting will be larger in magnitude when the trait variance is large (e.g., due to imperfect switching or high rates of suboptimal switching; $V(\bar{\alpha})$ large) or induction and loss of induction drive the mean trait far from the optimal trait value (resulting in $f_{\alpha_i} - g_{\alpha_i}$ large in magnitude). We predict that the effects of phenotypic sorting on equilibrium stability are negligibly small relative to the effects of phenotypic plasticity in most systems. This is because in simulations the magnitudes of the phenotypic sorting terms were negligibly small compared to the phenotypic plasticity terms. The only exception was when the induction and loss of induction rates were very slow ($\epsilon_I$ and $\epsilon_L$ very small) relative to the reproduction and predation rates of the prey. However, we expect the opposite in most natural systems, i.e., we expect prey reproduction and predation rates to be slow relative to induction and loss of induction rates.

For completeness, the following lists the biological conditions under which phenotypic sorting has a destabilizing effect. First, phenotypic sorting is destabilizing if individual fitness is maximized at extreme trait values (i.e., $f_{\alpha_i\alpha_i} - g_{\alpha_i\alpha_i} + f_{\alpha_i\bar{\alpha}} - g_{\alpha_i\bar{\alpha}} > 0$). It is also destabilizing if interactions between the trait variance and the individual prey fitness gradient are destabilizing (i.e., $V'(\bar{\alpha})(f_{\alpha_i} - g_{\alpha_i}$ positive). Second, if individual fitness is maximized at intermediate trait values (i.e., $f_{\alpha_i\alpha_i} - g_{\alpha_i\alpha_i} + f_{\alpha_i\bar{\alpha}} - g_{\alpha_i\bar{\alpha}} < 0$), then the small effects can be stabilizing when (i) the benefits of increased defense decrease as prey density decreases ($f_{x\alpha_i} - g_{x\alpha_i} < 0$) or (ii) for Class 2 inducible defenses, the benefits of increased defense increase as prey density decreases ($f_{x\alpha_i} - g_{x\alpha_i} > 0$).

##### S1.5.6 Effects of phenotypic sorting in the intragenerational inducible defense model where all prey offspring are initially uninduced

The effects of phenotypic sorting in model (S29) are determined by the following terms in the Jacobian entries

$$
\begin{aligned}
J_{31}: &\quad \overbrace{-(\bar{\alpha}-\alpha_{min})f_x}^{\text{positive}} - V(\bar{\alpha})g_{x\alpha_i}, \\
J_{32}: &\quad -V(\bar{\alpha})g_{y\alpha_i}, \\
J_{33}: &\quad \overbrace{-f}^{\text{negative}} \; \overbrace{-(\bar{\alpha}-\alpha_1)(f_{\bar{\alpha}}+f_{\alpha_i})}^{\text{positive}} - V(\bar{\alpha})(g_{\alpha_i\alpha_i}+g_{\alpha_i\bar{\alpha}}).
\end{aligned}
\tag{S90}
$$

For term $J_{31}$, if $J_{33}$ is negative, then the phenotypic sorting terms typically have a destabilizing effect when they make $J_{31}$ more negative. However, for Class 3 inducible defenses, they can have a destabilizing effect if they make $J_{31}$ more positive. For $J_{32}$, we expect the phenotypic sorting terms have a stabilizing effect because we expect $-Vg_{y\alpha_i} > 0$ in most systems; see the end of Section S1.5.4 for details. For $J_{33}$, the phenotypic sorting terms have a destabilizing effect when they make $J_{33}$ more positive. Note that the term $-f$ makes $J_{33}$ more negative (a destabilizing effect) and the term $-(\bar{\alpha}-\alpha_{min})(f_{\bar{\alpha}}+f_{\alpha_i})$ makes $J_{33}$ more positive (a destabilizing effect).

Interpretation: First, consider the effects of all prey offspring being initially uninduced (terms involving $f$ and its derivatives). This typically has a stabilizing effect on an equilibrium (because it makes $J_{31}$ more positive and typically makes $J_{33}$ more negative) and the stabilizing effect is larger in magnitude when per capita prey reproduction rates are large ($f$ large). A destabilizing effect is only possible if reproduction rates are low ($f$ small), nearly all prey are defended ($\bar{\alpha}$ is close to $\alpha_{max}$), and the costs for increased defense are high ($f_{\bar{\alpha}}+f_{\alpha_i}$ large in magnitude).

Now consider the effects of phenotypic sorting due to differing predation rates on the prey phenotypes (terms involving derivatives of $g$). We predict that these effects are negligibly small relative to the effects of phenotypic plasticity in most systems. This is because in simulations the magnitudes of the terms were negligibly small compared to the phenotypic plasticity terms. The only exception was when the induction and loss of induction rates were very slow ($\epsilon_I$ and $\epsilon_L$ very small) relative to the reproduction and predation rates of the prey. However, we expect the opposite in most natural systems, i.e., we expect prey reproduction and predation rates to be slow relative to induction and loss of induction rates. For completeness, we note that the terms have a destabilizing effect when (i) there are accelerating benefits of prey defense (i.e., predation rates decrease at accelerating rates as mean defense increases; $-Vg_{\alpha_i\alpha_i} - Vg_{\alpha_i\bar{\alpha}} > 0$ makes $J_{33}$ more positive), or (ii) increased defense reduces predation rates less when prey density is higher ($-Vg_{x\alpha_i} < 0$ makes $J_{31}$ more negative). For Class 2 inducible defenses, destabilizing effects can also occur if increased defense reduces predation rates more when prey density is higher ($-Vg_{x\alpha_i} > 0$ makes $J_{31}$ more positive).

##### S1.5.7 Effects of irreversibility on equilibrium stability

We explore how irreversibility alters the effects of inducible defenses by decreasing the value of $\epsilon_L$ in the intragenerational inducible defense model (S29) where all prey offspring are initially undefended. Decreasing $\epsilon_L$ has both stabilizing and destabilizing effects.

First, decreasing $\epsilon_L$ decreases the magnitudes of the phenotypic plasticity terms in the bottom row of the Jacobian (S30), which typically has a destabilizing effect. The reasoning is the following. Decreasing $\epsilon_L$ causes the terms multiplied by $\epsilon_L$ in the bottom row of the Jacobian to become smaller in magnitude. This decreases the (de)stabilizing effects caused by phenotypic plasticity component of trait change. Because the phenotypic plasticity component typically has a stabilizing effect, this mean irreversible traits are typically less stabilizing than reversible traits.

Second, decreasing $\epsilon_L$ causes the density dynamics to become more stable, which has a stabilizing effect on the system. The reasoning is the following. Decreasing $\epsilon_L$ causes the equilibrium trait value to increase because $\partial\bar{\alpha}^*/\partial\epsilon_L = -(\partial\dot{\bar{\alpha}}/\partial\epsilon_L)(-1)^6(J_{11}J_{22} - J_{21}J_{22})/|J| < 0$, where $\dot{\bar{\alpha}} = d\bar{\alpha}/dt$, $J_{11}J_{22} - J_{21}J_{12} > 0$, $|J| < 0$,and $\partial\dot{\bar{\alpha}}/\partial\epsilon_L < 0$. The increased value of $\bar{\alpha}$ stabilizes the density dynamics of the system because it decreases prey growth rates and decreases predation, which causes $J_{11} + J_{22}$ to decrease.

<u>Interpretation:</u> Irreversibility has both stabilizing and destabilizing effects on equilibria. The stabilizing effect of irreversibility is due to higher mean defense levels: increased defense means lower prey growth rates and lower predation rates, both of which have stabilizing effects on the density dynamics. The destabilizing effect of irreversibility is due to a weakening of the stabilizing feedback of the trait on its own dynamics. This means that irreversible traits can be more or less (de)stabilizing then reversible traits and the outcome depends on the specifics of the system. Stabilization is expected when increases in defense cause large decreases in prey growth rates and predation rates; destabilization is expected when the decreases are not as large.

Figure S1 shows three examples comparing equilibrium stability when the maximum loss of induction rate is (left column) equal to the maximum induction rate, (middle column) half of the maximum induction rate, or (right column) equal to zero. The top row of Figure S1 shows an example where an irreversible trait is more stabilizing than a reversible trait. In Figure S1AB, the equilibrium is unstable if mean defense is held fixed at its equilibrium value and the stabilizing effects of a reversible inducible defense are insufficiently strong to stabilize the equilibrium. Reducing $\epsilon_L$ to zero causes the density dynamics to become stable, and that stabilizing effect is greater than the destabilizing effect due to a weakening the feedback of the trait on its own dynamics. As a result, the systems converges to and equilibrium point when the trait is irreversible (Figure S1C).

The middle row of Figure S1 shows an example where an irreversible trait is less destabilizing than a reversible trait. In Figure S1DE, the equilibrium is stable if mean defense is held fixed at its equilibrium value and the destabilizing effects of a reversible trait destabilize the equilibrium. However, for an irreversible trait, the equilibrium is stable (Figure S1F) because the density dynamics become more stable and that stabilizing effect is greater than the destabilizing effect due to a weakening of the feedback of the trait on its own dynamics.

The bottom row of Figure S1 shows an example where an irreversible trait is more destabilizing than a reversible strait. In Figure S1GH, the equilibrium is stable if mean defense is held fixed at its equilibrium value and the equilibrium is not destabilized by a reversible inducible defense. However, for an irreversible trait, the equilibrium is destabilized (Figure S1I) via $J_{13}$ being large and negative. This destabilization is possible because while the density dynamics become more stable at $\epsilon_L$ decreases to 0, those stabilizing effects are weaker than the destabilizing effect due to a weakening of the feedback of the trait on its own dynamics.

###### S1.5.8 Effects of phenotypic sorting and transgenerational responses in the transgenerational inducible defense model

The effects of phenotypic sorting and transgenerational responses in the transgenerational inducible defense model (S49) are determined by the following terms in the entries of Jacobian (S51). Recall that the equilibrium condition $d\bar{\alpha}/dt = 0$ implies $[(\alpha_{max} - \bar{\alpha})P_I - (\bar{\alpha} - \alpha_{min})P_L] < 0$; see Section S1.3.2.

$$\begin{aligned}
J_{31}: &\quad \overbrace{[(\alpha_{max} - \bar{\alpha})P_I - (\bar{\alpha} - \alpha_{min})P_L]f_x}^{\text{transgenerational response}} \quad \overbrace{-Vg_{x\alpha_i}}^{\text{phenotypic sorting}} \\
J_{32}: &\quad \overbrace{-Vg_{y\alpha_i}}^{\text{phenotypic sorting}} \\
J_{33}: &\quad \overbrace{[(\alpha_{max} - \bar{\alpha})P_I - (\bar{\alpha} - \alpha_{min})P_L](f_{\bar{\alpha}} + f_{\alpha_i})}^{\text{transgenerational response}} \overbrace{-V(g_{\bar{\alpha}\alpha_i} + g_{\alpha_i\alpha_i})}^{\text{phenotypic sorting}}
\end{aligned} \tag{S91}$$

In numerical simulations, the magnitudes of the phenotypic sorting terms were much smaller in magnitude than the transgenerational response terms and negligibly small compared to the phenotypic plasticity terms, unless the induction and loss of induction rates were very slow ($\epsilon_I$ and $\epsilon_L$ very small). Because of this, we focus on the transgenerational response terms.

The transgenerational response term in $J_{31}$ makes it more positive, which has a stabilizing effect, except possibly for inducible defenses of Class 3. The transgenerational response term in $J_{33}$ make it more positive, which has a destabilizing effect.

**Interpretation:** In principle, transgenerational inducible defenses can be more or less stabilizing than intragenerational responses because of the stabilizing transgenerational term in $J_{31}$ and destabilizing transgenerational term in $J_{33}$. However, in our numerical simulations, the destabilizing effect in $J_{33}$ was larger in magnitude and it destabilized an equilibrium much more often than the stabilizing effect stabilized an equilibrium. From this we predict that transgenerational inducible defenses are less stabilizing and more destabilizing than intragenerational inducible defenses.

Figure S2 shows an example where a transgenerational inducible defense is less stabilizing than an otherwise equivalent intragenerational response. The equilibrium is unstable when the mean trait value is held fixed at its equilibrium value. In Figure S2A, the transgenerational inducible defense stabilizes the equilibrium when there is rapid turnover in the prey population ($\epsilon$ large, meaning prey reproduction rates and non-predator mortality rates are high; see Section S1.3). In Figure S2B, the transgenerational inducible defense is unable to stabilize the equilibrium when turnover in the prey population is low ($\epsilon = 0$, meaning reproduction rates are not high and all prey mortality is due to predation). Figure S2C shows a simulation from the intragenerational inducible defense model (S1) with true breeding prey that is otherwise identical to the transgenerational defense model in Figure S2B. Specifically,

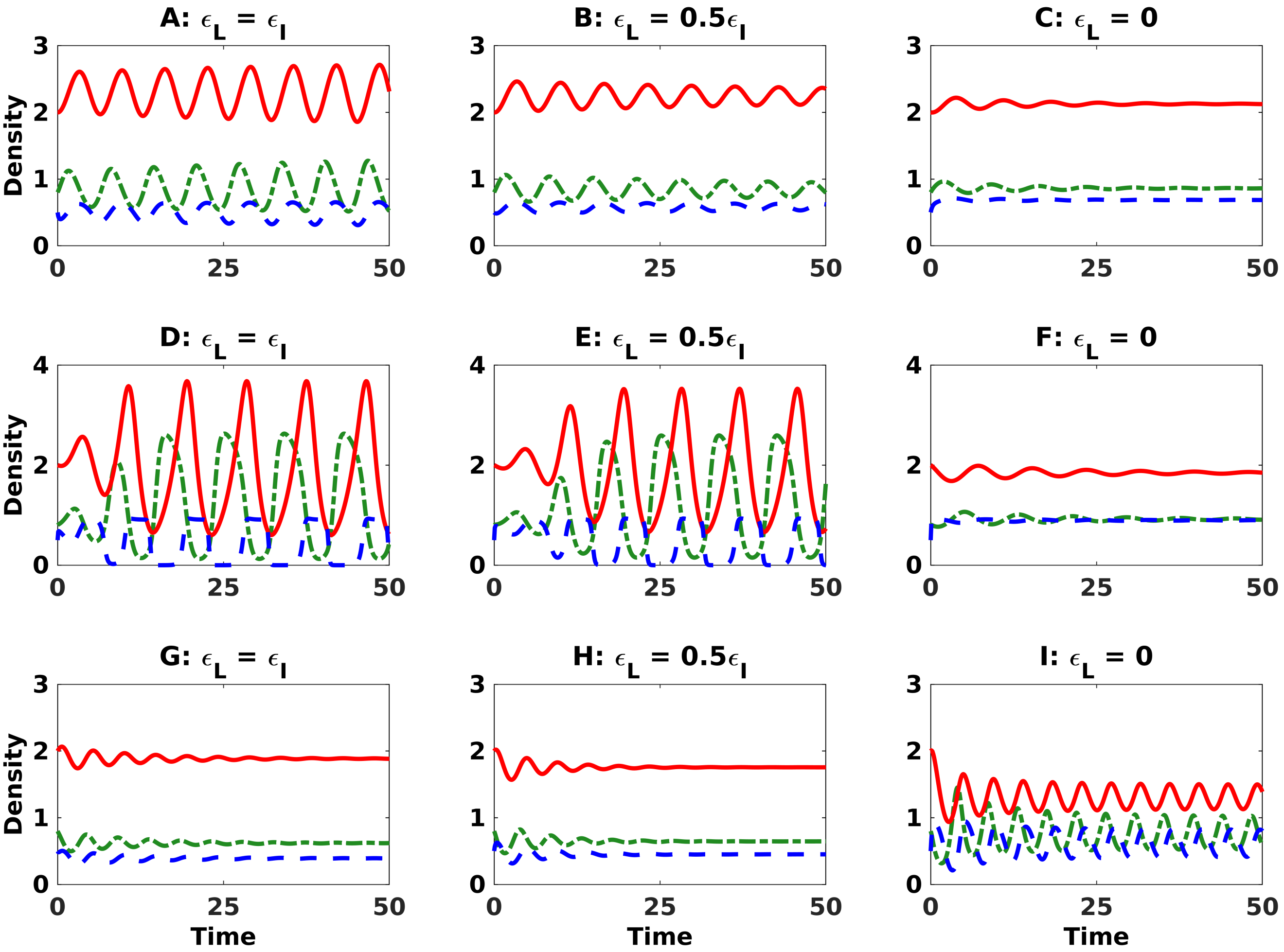

Figure S1: Examples of an irreversible inducible defense being (top row) more stabilizing, (middle row) less destabilizing, and (bottom row) more destabilizing than a reversible inducible defense. All panels show time series of prey density (dash-dot green), predator density (solid red), and mean defense (dashed blue). In each row, the maximum rate of induction is (left panel) equal to the maximum rate of induction, (middle panel) half the maximum rate of induction, or (right panel) zero, which corresponds to an irreversible defense. Additional details about the simulations are given in Section S1.5.7. Models and parameter values are given in Appendix S2.

all parameter values are the same and the values of $\epsilon_I$ and $\epsilon_L$ are set equal to $f$ at the equilibrium of the transgenerational defense model. Thus, at equilibrium, the Jacobians of the transgenerational and intragenerational defense models are nearly identical; differences only arise because of the intragenerational inducible defense model includes phenotypic sorting terms due to differential reproduction (i.e., $Vf_{\bar{\alpha}} + Vf_{\alpha_i}$) and whereas the Jacobian of the transgenerational defense has the transgenerational response terms listed in equation (S91). Importantly, while the transgenerational inducible defense cannot stabilize the equilibrium in Figure S2B, the otherwise identical intragenerational defense in Figure S2C stabilizes the equilibrium. In total, Figure S2BC shows that the transgenerational responses terms in the Jacobian have a destabilizing effect on the equilibrium.

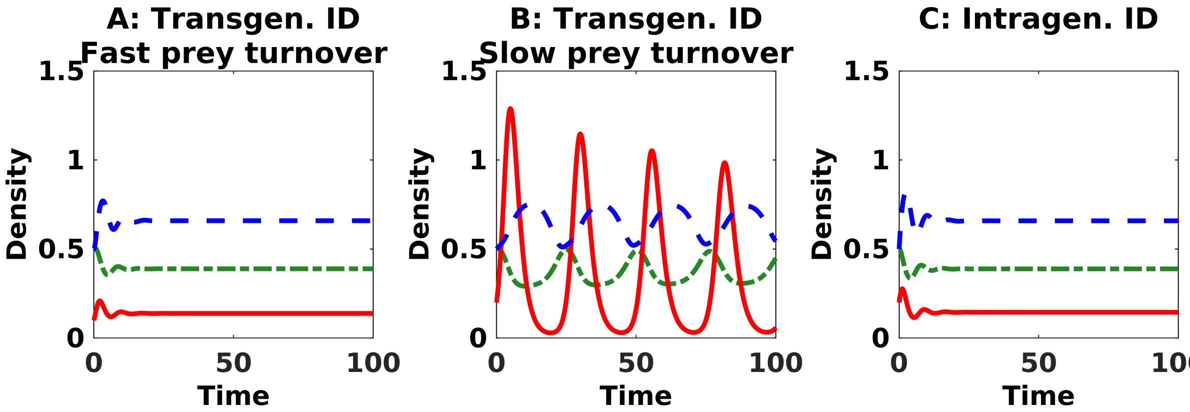

Figure S2: Example illustrating our prediction that transgenerational inducible defenses are typically less stabilizing or more destabilizing then intragenerational inducible defenses. (A) The transgenerational inducible defense stabilizes an unstable equilibrium when there is rapid turnover in the prey population. (B) The transgenerational inducible defense does not stabilize an unstable equilibrium when turnover turnover in the prey population is slower. (C) A intragenerational inducible defense that is otherwise equivalent to the transgenerational response in Panel B is stabilizing. Additional details about the simulations are given in Section S1.5.8. Models and parameter values are given in Appendix S2.

We predict that the effects of phenotypic sorting on equilibrium stability are negligibly small relative to the effects of phenotypic plasticity and transgenerational responses in most systems. This is because in simulations the magnitudes of the phenotypic sorting terms were much smaller than the transgenerational response terms and negligibly small compared to the phenotypic plasticity terms. The only exception was when the induction and loss of induction rates were very slow ($\epsilon_I$ and $\epsilon_L$ very small) relative to the reproduction and predation rates of the prey. However, we expect the opposite in most natural systems, i.e., we expect prey reproduction and predation rates to be slow relative to induction and loss of induction rates. For completeness, we list the biological conditions under which phenotypic sorting has a destabilizing effect. Destabilizing effects occur when (i) there are accelerating benefits of prey defense (i.e., predation rates decrease at accelerating rates as mean defense increases; $-Vg_{\alpha_i\alpha_i} - Vg_{\alpha_i\bar{\alpha}} > 0$ makes $J_{33}$ more positive), or (ii) increased defense reduces predation rates less when prey density is higher ($-Vg_{x\alpha_i} < 0$ makes $J_{31}$ more negative). For

Class 3 inducible defenses, destabilizing effects can also occur if increased defense reduces predation rates more when prey density is higher ($-V g_{x\alpha_i} > 0$ makes $J_{31}$ more positive). Note that the phenotypic sorting terms are larger in magnitude when the trait variance is large (e.g., due to imperfect switching or high rates of suboptimal switching; $V(\bar{\alpha})$ large) or induction and loss of induction rates drive the mean trait to values far from the optimal trait value (resulting in $f_{\alpha_i} - g_{\alpha_i}$ large in magnitude).

#### S1.6 Effects of inducible defenses on predator-prey phase lags

Following Cortez (2016), we compute the phase lags between the predator and prey oscillations at parameter values where the model undergoes a Hopf bifurcation. The approach is the following. Hopf bifurcations occur at parameter values satisfying $a_i > 0$ and $a_1 a_2 - a_3 = 0$, where the $a_i$ are defined in equations (S80)-(S82). We assume $-a_1 = J_{11} + J_{22} + J_{33} < 0$ and $J_{21} - J_{23}J_{31}/J_{33} > 0$. The first assumption guarantees that the cycles born during the Hopf bifurcation are attracting. The second assumption implies that there is a positive total effect of increased prey density on predator growth rate at the equilibrium, which maintains the sign of the predatory interaction. If that condition is not met, then the prey has a negative effect on the predator's growth rate, which aligns more with competitive interactions.

We further assume that at the Hopf bifurcation the prey oscillations are $x(t) = a\sin(\omega t)$, where $a$ is the amplitude and $\omega$ is the frequency of the oscillations. To compute $\omega$, we solve $a_1 a_2 - a_3 = 0$ for $J_{12}$, substitute into the Jacobian, and compute the eigenvalues explicitly using Maple 2019 (Maplesoft, Waterloo, Ontario). The eigenvalues at the Hopf bifurcation are $\lambda_1 = J_{11} + J_{22} + J_{33}$ and $\lambda_\pm = \pm\omega i$ where

$$\omega = \sqrt{-\frac{(J_{21}J_{33} - J_{23}J_{31})(J_{11}J_{33} - J_{13}J_{31} + J_{22}J_{33} - J_{23}J_{32}) + (J_{21}J_{32} - J_{22}J_{31})(J_{11}J_{23} - J_{13}J_{21})}{J_{23}J_{31} + J_{11}J_{21} + J_{21}J_{22}}}. \tag{S92}$$

The solutions to $y(t)$ and $\alpha(t)$ are found by solving the linearized system

$$\begin{aligned} \frac{dy}{dt} &= J_{21}x + J_{22}y + J_{23}\alpha \\ \frac{d\alpha}{dt} &= J_{31}x + J_{32}y + J_{33}\alpha \end{aligned} \tag{S93}$$

with $x(t) = a\sin(\omega t)$, which yields

$$\begin{aligned} y(t) =& \overbrace{\frac{a}{C}\left(-\omega^2[J_{21}J_{22} + J_{23}J_{31}] + J_{33}^2\left[-J_{22} + \frac{J_{23}J_{32}}{J_{33}}\right]\left[J_{21} - \frac{J_{23}J_{31}}{J_{33}}\right]\right)}^{C_{\sin}}\sin(\omega t) \\ & \overbrace{-\frac{a}{C}\omega\left(J_{21}\omega^2 + J_{33}^2\left[J_{21} - \frac{J_{23}J_{31}}{J_{33}}\right] + J_{23}[J_{21}J_{32} - J_{22}J_{31}]\right)}^{C_{\cos}}\cos(\omega t) \end{aligned} \tag{S94}$$

$$\begin{aligned} \alpha(t) =& -\frac{a}{C}\left([J_{31}J_{22} - J_{21}J_{32}][J_{22}J_{33} - J_{23}J_{32}] + [J_{31}J_{33} + J_{32}J_{21}]\omega^2\right)\sin(\omega t) \\ & -\frac{a}{C}\omega\left(J_{31}[\omega^2 + J_{22}^2] + J_{32}[J_{23}J_{31} - J_{21}J_{33}] - J_{21}J_{22}J_{32}\right)\cos(\omega t) \end{aligned} \tag{S95}$$

$$C = [J_{22} + J_{33}]^2\omega^2 + [J_{23}J_{32} - J_{22}J_{33} + \omega^2]^2 > 0. \tag{S96}$$

The predator lags behind the prey (i) by less than quarter-period when $C_{\sin} > 0$ and $C_{\cos} < 0$, (ii) between a quarter and a half period when $C_{\sin} < 0$ and $C_{\cos} < 0$, and (iii) by more than a half period when $C_{\cos} > 0$.

The goal in this section is to determine the signs of $C_{\sin}$ and $C_{\cos}$. For $C_{\sin}$, this reduces down to determining the signs of $J_{33}$ and $J_{31}$. This is because $C_{\sin}$ is positive unless $J_{33}$ is positive and sufficiently large in magnitude (making $J_{23}J_{32}/J_{33}$ negative) or $J_{31}$ is negative and and sufficiently large in magnitude (making $-\omega^2 J_{23}J_{31}$ negative). For $C_{\cos}$, we focus exclusively on systems where $C_{\cos} < 0$. The reason is that our numerical simulations suggest that for any model, $C_{\cos}$ is positive only in very small regions of parameter space (if at all). In particular, we constructed $3 \times 3$ matrices with random entries constrained to satisfy $J_{12} < 0$, $J_{21} > 0$, $J_{22} < 0$, and $J_{23} < 0$, $|J| < 0$, $J_{11} + J_{22} + J_{33}$, and $a_1a_2 - a_3 = 0$; the first four constraints ensure the Jacobian entries have signs consistent with the predator-prey interaction and the latter three constraints are necessary for a stable equilibrium to undergo a Hopf bifurcation. We then computed the value of $C_{\cos}$ for each matrix. Our numerical results show that matrices yielding $C_{\cos} > 0$ exist, but only in tiny regions of parameter space. That in turn implies that for any given model, $C_{\cos} > 0$ can only occur in tiny regions of parameter space. Thus, we expect that most, if not all, empirical systems will lie outside of the conditions corresponding to $C_{\cos} > 0$.

In the following subsections, we first show how the phenotypic plasticity components of trait change affect predator-prey phase lags for each class of inducible defense. As before, we do this by assuming the phenotypic sorting components of trait change in all models are negligible (mathematically, we set $V = 0$). We discuss our results in terms of the maximum rates of induction ($\epsilon_I$) and loss of induction ($\epsilon_L$) in the switch after birth models (S1) and (S29), but all of our result also apply to the switch at birth model (S49) after replacing $\epsilon_I$ and $\epsilon_L$ with $f$; see Section S1.3.2. The remaining subsections address how the predictions are altered by the phenotypic sorting terms for each model, irreversibility, and transgenerational responses.

##### S1.6.1 Class 1: Induction stimulus is predator density, $P_i(y)$

Assume $V = 0$, $\partial P_I/\partial x = 0$, $\partial P_I/\partial y > 0$, and $\partial P_I/\partial \alpha = 0$. This results in $J_{31} = 0$, $J_{32} > 0$, and $J_{33} < 0$. Substituting into the formulas for $C_{\sin}$ and $C_{\cos}$ yields

$$C_{\sin} = \frac{a}{C}\left( \overbrace{-\omega^2[J_{21}J_{22}]}^{\text{positive}} \overbrace{-J_{33}^2 \left[J_{22} - \frac{J_{23}J_{32}}{J_{33}}\right]\left[J_{21} - \frac{J_{23}J_{31}}{J_{33}}\right]}^{\text{positive}} \right) > 0 \tag{S97}$$

$$C_{\cos} = \frac{a}{C}\omega\left( \overbrace{-J_{21}\omega^2 - J_{33}^2 J_{21}}^{\text{negative}} \overbrace{-J_{23}J_{21}J_{32}}^{\text{positive}} \right). \tag{S98}$$

As explained in Section S1.6, we assume $C_{\cos} < 0$, which holds for all but a tiny region of parameter space.

Interpretation: Class 1 inducible defenses shorten predator-prey phase lags, yielding lags less than a quarter-period.

##### S1.6.2 Class 2: Induction stimulus are predator and prey densities, $P_i(x, y)$

Assume $V = 0$, $\partial P_I/\partial x < 0$, $\partial P_I/\partial y > 0$, and $\partial P_I/\partial \alpha = 0$. This results in $J_{31} < 0$, $J_{32} > 0$, and $J_{33} < 0$. Substituting into the formulas for $C_{\sin}$ and $C_{\cos}$ yields

$$C_{\sin} = \frac{a}{C}\left(\overbrace{-\omega^2[J_{21}J_{22}]}^{\text{positive}}\overbrace{-\omega^2 J_{23}J_{31}}^{\text{negative}}\overbrace{-J_{33}^2\left[J_{22} - \frac{J_{23}J_{32}}{J_{33}}\right]\left[J_{21} - \frac{J_{23}J_{31}}{J_{33}}\right]}^{\text{positive}}\right) \tag{S99}$$

$$C_{\cos} = \frac{a}{C}\omega\left(\overbrace{-J_{21}\omega^2 - J_{33}^2\left[J_{21} - \frac{J_{23}J_{31}}{J_{33}}\right] + J_{23}J_{22}J_{31}}^{\text{negative}}\overbrace{-J_{23}J_{21}J_{32}}^{\text{positive}}\right). \tag{S100}$$

In most cases, $C_{\sin}$ is positive. However, $C_{\sin}$ can be negative if $J_{31}$ is negative and large in magnitude. As explained in Section S1.6, we assume $C_{\cos} < 0$, which holds for all but a tiny region of parameter space.

Now consider the special case where induction and loss of induction rates are much faster than changes in prey and predator densities. Collecting the highest order terms for $\omega$ and substituting into the formulas for $C_{\sin}$ and $C_{\cos}$ yields

$$\omega = \sqrt{-\frac{(J_{21}J_{33} - J_{23}J_{31})(J_{11}J_{33} + J_{22}J_{33} - J_{23}J_{32} - J_{13}J_{31}}{J_{23}J_{31})}} \tag{S101}$$

$$C_{\sin} = -J_{23}J_{31}\omega^2 < 0 \tag{S102}$$

$$C_{\cos} = -J_{33}^2\left(J_{21} - \frac{J_{23}J_{31}}{J_{33}}\right) - J_{21}\omega^2 < 0. \tag{S103}$$

Thus, the lags are between a quarter-period and a half period.

Interpretation: Class 2 inducible defenses typically shorten predator-prey phase lags, yielding lags less than a quarter-period. However, Class 2 inducible defenses can increase predator-prey phase lags, yielding lags between a quarter-period and a half-period, when (i) induction rates are very sensitive to prey density ($J_{31} > 0$ large in magnitude) or (ii) rates of induction and loss of induction are so fast that changes in the mean trait value are much faster than changes in densities.

##### S1.6.3 Class 3: Induction stimulus is the predation rate, $P_i(xg)$

Assume $V = 0$, $\partial P_I/\partial x > 0$, $\partial P_I/\partial y > 0$, and $\partial P_I/\partial \alpha < 0$. This results in $J_{31} > 0$, $J_{32} > 0$, and $J_{33} < 0$. Substituting into the formulas for $C_{\sin}$ and $C_{\cos}$ yields

$$C_{\sin} = \frac{a}{C}\left(\overbrace{-\omega^2[J_{21}J_{22}]}^{\text{positive}}\overbrace{-\omega^2 J_{23}J_{31}}^{\text{positive}}\overbrace{-J_{33}^2\left[J_{22} - \frac{J_{23}J_{32}}{J_{33}}\right]\left[J_{21} - \frac{J_{23}J_{31}}{J_{33}}\right]}^{\text{more positive}}\right) > 0 \tag{S104}$$

$$C_{\cos} = \frac{a}{C}\omega\left(\overbrace{-J_{21}\omega^2 - J_{33}^2\left[J_{21} - \frac{J_{23}J_{31}}{J_{33}}\right]}^{\text{more negative}} + \overbrace{J_{23}J_{22}J_{31} - J_{23}J_{21}J_{32}]}^{\text{more positive}}\right). \tag{S105}$$

As explained in Section S1.6, we assume $C_{\cos} < 0$, which holds for all but a tiny region of parameter space.

Underlined: Interpretation: Class 3 inducible defenses shorten predator-prey phase lags, yielding lags less than a quarter-period.

##### S1.6.4 Class 4: Induction stimulus is the prey fitness gradient, $P_i(\frac{\partial}{\partial\alpha_i}\frac{1}{x}\frac{dx}{dt})$

Assume $V = 0$. Assume the signs of $\partial P_I/\partial x$, $\partial P_I/\partial y$, and $\partial P_I/\partial\alpha$ are the same as $f_{x\alpha_i} - g_{x\alpha_i}$, $f_{y\alpha_i} - g_{y\alpha_i}$, and $f_{\alpha_i\bar{\alpha}} - g_{\alpha_i\bar{\alpha}} + f_{\alpha_i\alpha_i} - g_{\alpha_i\alpha_i}$, respectively, and assume the sign of $\partial P_L/\partial x$, $\partial P_L/\partial y$, and $\partial P_L/\partial\alpha$ are the opposite. This means $J_{31} = \epsilon C_{31}(f_{x\alpha_i} - g_{x\alpha_i})$, $J_{32} = \epsilon C_{32}(-g_{y\alpha_i})$, and $J_{33} = \epsilon P_I - \epsilon P_L + \epsilon C_{33}(f_{\alpha_i\bar{\alpha}} - g_{\alpha_i\bar{\alpha}} + f_{\alpha_i\alpha_i} - g_{\alpha_i\alpha_i})$.

The coefficients $C_{\sin}$ and $C_{\cos}$ are

$$C_{\sin} = \frac{a}{C}\left(\omega^2[\overbrace{-J_{21}J_{22}}^{\text{positive}}\overbrace{-J_{23}J_{31}}^{J_{31}}] + J_{33}^2\left[\overbrace{-J_{22}}^{\text{positive}} + \overbrace{\frac{J_{23}J_{32}}{J_{33}}}^{-J_{32}J_{33}}\right]\overbrace{\left[J_{21} - \frac{J_{23}J_{31}}{J_{33}}\right]}^{\text{positive}}\right) \tag{S106}$$

$$C_{\cos} = \frac{a}{C}\omega\left(\overbrace{-J_{21}\omega^2}^{\text{negative}}\overbrace{-J_{33}^2\left[J_{21} - \frac{J_{23}J_{31}}{J_{33}}\right]}^{\text{negative}} + \overbrace{-J_{23}J_{21}J_{32}}^{J_{32}} + \overbrace{J_{23}J_{22}J_{31}]}^{J_{31}}\right) \tag{S107}$$

In most systems, we expect $C_{\sin} > 0$. However, $C_{\sin} < 0$ is possible if (i) $J_{31}$ is negative and large in magnitude ($f_{x\alpha_i} - g_{x\alpha_i} < 0$), (ii) $J_{32}$ and $J_{33}$ are positive and large in magnitude ($-g_{y\alpha_i} > 0$ and $f_{\alpha_i\bar{\alpha}} - g_{\alpha_i\bar{\alpha}} + f_{\alpha_i\alpha_i} - g_{\alpha_i\alpha_i} > 0$), or (iii) $J_{32}$ and $J_{33}$ are negative ($-g_{y\alpha_i} < 0$ and $f_{\alpha_i\bar{\alpha}} - g_{\alpha_i\bar{\alpha}} + f_{\alpha_i\alpha_i} - g_{\alpha_i\alpha_i} < 0$). The latter condition is unlikely to be satisfied because we expect $-g_{y\alpha_i} < 0$ to rarely be satisfied in natural systems; see the last paragraph of Section S1.5.4. As explained in Section S1.6, we assume $C_{\cos} < 0$, which holds for all but a tiny region of parameter space.

Now consider the special case where induction and loss of induction rates are much faster than changes in prey and predator densities. Note that the analysis of this case is only biologically informative when $J_{33}$ is negative. Collecting the highest order terms for $\omega$ and substituting into the formulas for $C_{\sin}$ and $C_{\cos}$ yields

$$C_{\sin} = -J_{23}J_{31}\omega^2 \tag{S108}$$

$$C_{\cos} = -J_{33}^2\left(J_{21} - \frac{J_{23}J_{31}}{J_{33}}\right) - J_{21}\omega^2 < 0. \tag{S109}$$

Thus, $C_{\sin}$ can be negative if $J_{31}$ is negative and sufficiently large in magnitude. $C_{\sin}$ is positive otherwise.

Interpretation: First consider systems where prey fitness is maximized at extreme trait values $(f_{\alpha_i\bar{\alpha}} - g_{\alpha_i\bar{\alpha}} + f_{\alpha_i\alpha_i} - g_{\alpha_i\alpha_i} > 0)$. Then the phenotypic plasticity component can lengthen the phase lags whenever the curvature of the fitness landscape is sufficiently large ($f_{\alpha_i\bar{\alpha}} - g_{\alpha_i\bar{\alpha}} + f_{\alpha_i\alpha_i} - g_{\alpha_i\alpha_i}$ large in magnitude). Note that it is possible for $J_{33}$ to be negative when prey fitness is maximized at extreme trait values. In this special case, the conditions for increased phase lags are the same as those in the next paragraph. However, as noted in Section S1.5.4, we expect this case to be rare in nature.

Second, consider systems where individual prey fitness is maximized by intermediate trait values $(f_{\alpha_i\bar{\alpha}} - g_{\alpha_i\bar{\alpha}} + f_{\alpha_i\alpha_i} - g_{\alpha_i\alpha_i} < 0)$. The phenotypic plasticity component typically shortens the phase lags less in predator-prey cycles, yielding lags less than a quarter-period. The phase lags can increase when (i) the benefits of higher mean defense decrease with increased prey density $(f_{x\alpha_i} - g_{x\alpha_i} < 0)$, or (ii) the benefits of higher defense decrease with increased predator density $(-g_{y\alpha_i} < 0)$. As noted in Section S1.5.4, condition (ii) is unlikely to be satisfied in natural systems.

##### S1.6.5 Effects of phenotypic sorting in the intragenerational inducible defense model with true breeding prey

For model (S1), the effects of phenotypic sorting on Jacobian entries $J_{31}$ and $J_{33}$ are determined by the following terms

$$\begin{aligned} J_{31}:&\quad V(\bar{\alpha})(f_{x\alpha_i} - g_{x\alpha_i}) \\ J_{33}:&\quad V(\bar{\alpha})(f_{\alpha_i\alpha_i} - g_{\alpha_i\alpha_i} + f_{\alpha_i\bar{\alpha}} - g_{\alpha_i\bar{\alpha}}) + V'(\bar{\alpha})(f_{\alpha_i} - g_{\alpha_i}) \end{aligned} \tag{S110}$$

The phenotypic sorting terms make $J_{31}$ more negative when $f_{x\alpha_i} - g_{x\alpha_i} < 0$. The phenotypic sorting terms make $J_{33}$ more positive when $f_{\alpha_i\alpha_i} - g_{\alpha_i\alpha_i} + f_{\alpha_i\bar{\alpha}} - g_{\alpha_i\bar{\alpha}} > 0$ or $V'(\bar{\alpha})(f_{\alpha_i} - g_{\alpha_i}) > 0$.

Interpretation: In general, the effects of phenotypic sorting will be larger in magnitude when the trait variance is large (e.g., due to imperfect switching or high rates of suboptimal switching; $V(\bar{\alpha})$ large) or induction and loss of induction drive the mean trait far from the optimal trait value (implying $f_{\alpha_i} - g_{\alpha_i}$ large in magnitude). We predict that the effects of phenotypic sorting on predator-prey phase lags are negligibly small relative to the effects of phenotypic plasticity in most systems. This is because in simulations the magnitudes of the phenotypic sorting terms were negligibly small compared to the phenotypic plasticity terms. The only exception was when the induction and loss of induction rates were very slow ($\epsilon_I$ and $\epsilon_L$ very small) relative toe reproduction and predation rates of the prey. However, we expect the opposite in most natural systems, i.e., we expect prey reproduction and predation rates to be slow relative to induction and loss of induction rates.

For completeness, the following lists the biological conditions under which phenotypic sorting increases predator-prey phase lags. First, phenotypic sorting typically increases the phase lag when individual prey fitness is maximized at extreme trait values $(f_{\alpha_i\bar{\alpha}} - g_{\alpha_i\bar{\alpha}} + f_{\alpha_i\alpha_i} - g_{\alpha_i\alpha_i} > 0)$. Second, when individual prey fitness is maximized at intermediate trait values $(f_{\alpha_i\bar{\alpha}} - g_{\alpha_i\bar{\alpha}} + f_{\alpha_i\alpha_i} - g_{\alpha_i\alpha_i} < 0)$, the phenotypic sorting terms typically shorten the phase lag unless the benefits of higher mean defense decrease with increased prey density

($f_{x\alpha_i} - g_{x\alpha_i} < 0$). These effects are identical to the effects of an evolving defense (Cortez 2016).

##### S1.6.6 Effects of phenotypic sorting in the intragenerational inducible defense model where all prey offspring are initially uninduced

For model (S29), the effects of phenotypic sorting on Jacobian entries $J_{31}$ and $J_{33}$ are determined by the following terms

$$\begin{aligned} J_{31} : &\quad \overbrace{-(\bar{\alpha} - \alpha_{min}) f_x}^{\text{positive}} - V(\bar{\alpha}) g_{x\alpha_i}, \\ J_{33} : &\quad \overbrace{-f}^{\text{negative}} \; \overbrace{-(\bar{\alpha} - \alpha_1)(f_{\bar{\alpha}} + f_{\alpha_i})}^{\text{positive}} - V(\bar{\alpha})(g_{\alpha_i\alpha_i} + g_{\alpha_i\bar{\alpha}}). \end{aligned} \tag{S111}$$

The phenotypic sorting terms typically make $J_{31}$ more positive, but they can make $J_{31}$ more negative when $f_{x\alpha_i} - g_{x\alpha_i} < 0$ is sufficiently large in magnitude. The phenotypic sorting terms typically make $J_{33}$ more negative, but they can make $J_{33}$ more positive if (i) $f_{\bar{\alpha}} + f_{\alpha_i} < 0$ is sufficiently large in magnitude or (ii) $f_{\alpha_i\alpha_i} - g_{\alpha_i\alpha_i} + f_{\alpha_i\bar{\alpha}} - g_{\alpha_i\bar{\alpha}} > 0$ and $V'(\bar{\alpha})(f_{\alpha_i} - g_{\alpha_i}) > 0$ are sufficiently large in magnitude.

Interpretation: First, consider the effects of all prey offspring being initially uninduced (terms involving $f$ and its derivative). This typically reduces predator-prey phase lags (because it makes $J_{31}$ more positive and typically makes $J_{33}$ more negative). The phase lags can be increased only if reproduction rates are low ($f$ small), nearly all prey are defended ($\bar{\alpha}$ is close to $\alpha_{max}$), and the costs for increased defense are high ($f_{\bar{\alpha}} + f_{\alpha_i}$ large in magnitude).

Now consider the effects of phenotypic sorting due to differing predation rates on the prey phenotypes (terms involving derivatives of $g$). We predict that these effects are negligibly small relative to the magnitudes of the effects of phenotypic plasticity. This is because in simulations the magnitudes of the terms were negligibly small compared to the phenotypic plasticity terms. The only exception was when the induction and loss of induction rates were very slow ($\epsilon_I$ and $\epsilon_L$ very small) relative to the reproduction and predation rates of the prey. However, we expect the opposite in most natural systems, i.e., we expect prey reproduction and predation rates to be slow relative to induction and loss of induction rates. For completeness, we note that phenotypic sorting due to differential predation rates can increase the phase lags when (i) there are accelerating benefits of prey defense (i.e., predation rates decrease at accelerating rates as mean defense increases; $-Vg_{\bar{\alpha}_i} - Vg_{\alpha_i\alpha_i} > 0$ makes $J_{33}$ more positive) or (ii) increased defense reduces predation rates less when prey density is higher ($-Vg_{x\alpha_i} < 0$ makes $J_{31}$ more negative).

##### S1.6.7 Effects of irreversibility on predator-prey phase lags

As in Section S1.5.7, we explore how irreversibility alters the effects of inducible defense by decreasing the value of $\epsilon_L$ in the intragenerational inducible defense model (S29) where all prey offspring are initially undefended.

Decreasing $\epsilon_L$ can cause the phase lag to increase or decrease. However, in numerical simulations, decreasing $\epsilon_L$ never caused the phase lag to increase above a quarter-period. There are two reason for this. First, decreasing $\epsilon_L$ weakens the effects of each variable on the trait dynamics because the terms multiplied by $\epsilon_L$ decrease to zero. This weakening of effects cannot cause $J_{33}$ to change from negative to positive or $J_{31}$ to change from positive to negative; one of these is necessary for lags greater than quarter-period. Second, decreasing $\epsilon_L$ causes the density dynamics to become more stable (see Section S1.5.7 for the explanation). That increases the effect of the density dynamics on the phase lags, making the phase lags closer to a quarter period. Note that this could cause the phase lags to increase or decrease depending on whether the lags were longer or shorter than a quarter-period before $\epsilon_L$ was decreased. Combining the above two effects yields that decreasing $\epsilon_L$ causes the phase lags to increase or decrease towards a quarter-period.

Interpretation: Irreversible traits can yield phase lags that are greater or less than the phase lags caused by reversible traits. Our results suggest that when compared to reversible inducible defenses, irreversible inducible defenses are less likely increase predator-prey phase lags above a quarter-period and the lags are more likely to be closer to a quarter-period. Note that the analysis above does not address the phenotypic sorting terms due to differing predation rates across phenotypes (terms involving derivatives of $g$). This is because in our numerical simulations those terms were negligibly small relative to the phenotypic plasticity terms and the other phenotypic sorting terms.

##### S1.6.8 Effects of phenotypic sorting and transgenerational responses in the transgenerational inducible defense model

For the transgenerational inducible defense model (S49), the effects of phenotypic sorting and transgenerational responses on Jacobian entries $J_{31}$ and $J_{33}$ are determined by the following terms. Recall that the equilibrium condition $d\bar{\alpha}/dt = 0$ implies $[(\alpha_{max} - \bar{\alpha})P_I - (\bar{\alpha} - \alpha_{min})P_L] < 0$; see Section S1.3.2.

$$\begin{aligned} J_{31}: &\quad \overbrace{[(\alpha_{max} - \bar{\alpha})P_I - (\bar{\alpha} - \alpha_{min})P_L]f_x}^{\text{transgenerational response}} \quad \overbrace{-V g_{x\alpha_i}}^{\text{phenotypic sorting}} \\ J_{33}: &\quad \overbrace{[(\alpha_{max} - \bar{\alpha})P_I - (\bar{\alpha} - \alpha_{min})P_L](f_{\bar{\alpha}} + f_{\alpha_i})}^{\text{transgenerational response}} \overbrace{-V(g_{\bar{\alpha}\alpha_i} + g_{\alpha_i\alpha_i})}^{\text{phenotypic sorting}} \end{aligned} \tag{S112}$$

The transgenerational response terms make $J_{31}$ and $J_{33}$ more positive; the former reduces phase lags and the latter increases phase lags. The phenotypic sorting terms increase the phase lags when they make $J_{31}$ more negative and $J_{33}$ more positive, meaning $-g_{x\alpha_i} < 0$ and $-g_{\bar{\alpha}\alpha_i} + g_{\alpha_i\alpha_i} > 0$.

**Interpretation:** In numerical simulations, the magnitudes of the phenotypic sorting terms were much smaller in magnitude than the transgenerational response terms and negligibly small compared to the phenotypic plasticity terms, unless the induction and loss of induction rates were very slow ($\epsilon_I$ and $\epsilon_L$ very small). Because of this, we focus on effects of the transgenerational response terms.

Transgenerational responses can yield phase lags that are shorter or longer than the phase lags caused by intragenerational responses. Shorter lags occur when intraspecific prey competition is stronger (making the transgenerational response terms of $J_{31}$ large in magnitude) and longer lags occur when the costs for defense are higher (making the transgenerational terms of $J_{33}$ large in magnitude). We predict that the effects of transgenerational responses are not large enough to cause lags greater than quarter-period if such lags would not arise for an otherwise intragenerational response. Said another way, we predict transgenerational inducible defenses can drive cycles with phase lags greater than quarter-period only if those lags are caused by the response stimuli.

Our justification for this prediction is based on numerical simulations described below. We ran many numerical simulations where we compared the phase lags for transgenerational inducible defenses and otherwise equivalent intragenerational inducible defenses. Here, 'otherwise equivalent' means all parameter values are the same for the transgenerational model (S49) and the intragenerational defense model (S1) and the values of $\epsilon_I$ and $\epsilon_L$ are set equal to $f$ at the equilibrium of the transgenerational defense model. Thus, at equilibrium, the Jacobians of the transgenerational and intragenerational defense models are nearly identical, with differences only arising because of the intragenerational inducible defense model includes phenotypic sorting terms due to differential reproduction whereas the Jacobian of the transgenerational defense model has the transgenerational response terms listed in equation (S112).

In numerical simulations we found the following. First, the transgenerational terms were never large enough to make $J_{33}$ positive when the phenotypic plasticity terms were negative. Second, we found many examples where the phase lag for a transgenerational inducible defense was closer to a quarter-period than an otherwise equivalent intragenerational inducible defense. In these simulations, the transgenerational response lengthened a lag that was less than quarter-period and shortened a lag that was greater than a quarter-period. Third, we also found many examples where the phase lag for a transgenerational inducible defense was less than a quarter-period and the phase lag for an otherwise equivalent intragenerational inducible defense was greater than a quarter-period. In these simulations, the transgenerational response shortened the lag considerably. Fourth, we never found an example where the phase lag for a transgenerational inducible defense was greater than a quarter-period and the phase lag for an otherwise equivalent intragenerational inducible defense was less than a quarter-period. In total, this numerical work shows that transgenerational inducible defenses can increase or decrease predator-prey phase lags relative to intragenerational responses and relative to cycles where prey have fixed defense. It also suggests that transgenerational inducible defense can drive cycles with phase lags greater than quarter-period only if those lags are caused by the response stimuli.

### S2 Parameter values for simulations

Figures 1 and 2 of the main text use the continuous trait model (2) with true breeding prey. To illustrate that the predicted dynamics and effects can occur in relatively simple models, we use the following model.

$$\begin{aligned}
\frac{dx}{dt} &= \overbrace{x(r(\alpha) - kx)}^{\text{reproduction}} - \overbrace{\frac{b(\alpha)xy}{1 + b(\alpha)hx}}^{\text{predation}} \\
\frac{dy}{dt} &= \overbrace{\frac{cb(\alpha)xy}{1 + b(\alpha)hx}}^{\text{harvesting}} - \overbrace{my}^{\text{mortality}} \\
\frac{d\alpha}{dt} &= \underbrace{\overbrace{\epsilon_I P_i(\alpha_1 - \alpha)}^{\text{induction}} - \overbrace{\epsilon_L P_L(\alpha - \alpha_L)}^{\text{loss of induction}}}_{\text{phenotypic plasticity}} + \underbrace{V(f_{\alpha_i} - g_{\alpha_i})}_{\text{phenotypic sorting}} .
\end{aligned} \tag{S113}$$

In the prey equation, $r(\alpha) = r_0 + r_1\alpha$ is the density independent exponential growth rate of the prey, $k$ is the level of intraspecific competition, and $b(\alpha)xy/(1+b(\alpha)hx)$ is the predator's Type II functional response where $b(\alpha) = b_0 + b_1\alpha$ is the predator attack rate and $h$ is the handling time. In the predator equation, $c$ is the prey to predator conversion efficiency and $m$ is the predator per capita mortality rate. In the phenotypic plasticity component of the trait equation, $\epsilon_I$ and $\epsilon_L$ are the maximum rates of induction and loss of induction and $P_I$ and $P_L$ are non-negative functions that depend on the stimuli (i.e., state variables) and satisfy $P_I + P_L = 1$. In the phenotypic sorting component of the trait equation, $V$ is the trait variance, $f_{\alpha_i} = r_{\alpha_i}$, and $g_{\alpha_i} = b_{\alpha_i}y/(1 + bhx)$. This continuous trait model can be derived from a dimorphic model where each prey type has growth rate $x_i f_i = x_i[r(\alpha_i) - kx]$ and predation rate $x_i b(\alpha_i)y/[1+b(\alpha_1)hx_1+b(\alpha_2)hx_2]$. Without loss of generality, all simulations in the main text use the parameter values $\alpha_1 = 0$ and $\alpha_2 = 1$.

**Figure 1A-C:** $P_i(y) = [1 + (y/q)^\theta]$, $r_0 = 3$, $r_1 = -1$, $k = 1$, $b_0 = 2$, $b_1 = -1$, $h = 1$, $c = 2.5$, $m = 1$, $q = 1$, $\theta = 2$, $V = 0.1$, $\epsilon_I = \epsilon_L = 0.5$ for panel B and $\epsilon_I = \epsilon_L = 2$ for panel C.

**Figure 1D-F:** $P_i(y) = [1 + (y/q)^\theta]$, $r_0 = 3$, $r_1 = -1$, $k = 1$, $b_0 = 1.1$, $b_1 = -0.1$, $h = 1$, $c = 2.2$, $m = 1$, $q = 2$, $\theta = 5$, $V = 0.1$, $\epsilon_I = \epsilon_L = 1$ for panel E and $\epsilon_I = \epsilon_L = 1.9$ for panel F.

**Figure 1G-I:** $P_i(y) = [1 + (y/xq)^\theta]$, $r_0 = 3$, $r_1 = -1$, $k = 1$, $b_0 = 2$, $b_1 = -0.5$, $h = 1$, $c = 1.5$, $m = 1$, $q = 2$, $\theta = 5$, $V = 0.1$, $\epsilon_I = \epsilon_L = 0.1$ for panel H and $\epsilon_I = \epsilon_L = 1$ for panel I.

**Figure 2A,D:** $P_i(y) = [1 + (y/q)^\theta]$, $r_0 = 2.7$, $r_1 = -1$, $k = 1$, $b0 = 1.9$, $b1 = -0.9$, $h = 1$, $c = 2.5$, $m = 1$, $q = 2.3$, $V = 1$, $\theta = 10$, $\epsilon_I = \epsilon_L = 1.8$.

**Figure 2B,E:** $P_i(y) = [1 + (y/xq)^\theta]$, $r_0 = 3.5$, $r_1 = -0.8$, $k = 1$, $b_0 = 1.8$, $b_1 = -1$, $h = 1.3$, $c = 1.7$, $m = 1$, $q = 2$, $\theta = 10$, $V = 1$, $\epsilon_I = \epsilon_L = 1$.

**Figure 2C,F:** $P_i(y) = [1+\exp(-\theta(f_{\alpha_i} - g_{\alpha_i}))]$, $f_{\alpha_i} - g_{\alpha_i} = r' - b'y/(1+hbx)$ is the individual fitness gradient, $r_0 = 1.8$, $r_1 = 0.4$, $k = 1$, $b_0 = 6.66$, $b_1 = -6$, $h = 0.5$, $c = 1$, $m = 1$, $\theta = 25$, $V = 1$, $\epsilon_I = \epsilon_L = 1$.

Figure S1 uses a continuous trait model (4) where all prey offspring are initially undefended. The specific model is

$$
\begin{aligned}
\frac{dx}{dt} &= \overbrace{x(r(\alpha) - kx)}^{\text{reproduction}} - \overbrace{\frac{b(\alpha)xy}{1 + b(\alpha)hx}}^{\text{predation}} \\
\frac{dy}{dt} &= \overbrace{\frac{cb(\alpha)xy}{1 + b(\alpha)hx}}^{\text{harvesting}} - \overbrace{my}^{\text{mortality}} \\
\frac{d\alpha}{dt} &= \underbrace{\overbrace{\epsilon_I P_i(\alpha_1 - \alpha)}^{\text{induction}} - \overbrace{\epsilon_L P_L(\alpha - \alpha_L)}^{\text{loss of induction}}}_{\text{phenotypic plasticity}} + \underbrace{-\alpha f - V g_{\alpha_i}}_{\text{phenotypic sorting}}
\end{aligned}
\qquad \text{(S114)}
$$

where all functions are defined as above. This continuous trait model can be derived from a dimorphic model where each prey type has growth rate $x_i f_i = x_i[r(\alpha_i) - kx]$ and predation rate $x_i b(\alpha_i)y/[1 + b(\alpha_1)hx_1 + b(\alpha_2)hx_2]$. Without loss of generality, the simulations use the parameter values $\alpha_1 = 0$ and $\alpha_2 = 1$.

**Figure S1A-C:** $P_i(y) = [1 + (y/q)^\theta]$, $r_0 = 3$, $r_1 = -1$, $k = 1$, $b_0 = 2$, $b_1 = -1$, $h = 1$, $c = 2.5$, $m = 1$, $q = 2$, $\theta = 5$, and $V = 1$, where $\epsilon_I = 5$ and $\epsilon_L = 5$ for panel A, $\epsilon_I = 5$ and $\epsilon_L = 2.5$ for panel B, and $\epsilon_I = 5$ and $\epsilon_L = 0$ for panel C.

**Figure S1D-F:** $P_i(y) = [1 + (y/xq)^\theta]$, $r_0 = 3$, $r_1 = -1$, $k = 1$, $b0 = 2$, $b1 = -0.5$, $h = 1$, $c = 1.7$, $m = 1$, $q = 2$, $\theta = 5$, $V = 1$, where $\epsilon_I = 20$ and $\epsilon_L = 20$ for panel D, $\epsilon_I = 20$ and $\epsilon_L = 10$ for panel E, and $\epsilon_I = 20$ and $\epsilon_L = 0$ for panel F.

**Figure S1I-K:** $P_i(y) = [1 + (y/xq)^\theta]$, $r_0 = 3$, $r_1 = -2.2$, $k = 1$, $b_0 = 2$, $b_1 = -1$, $h = 1$, $c = 2$, $m = 1$, $q = 2$, $\theta = 5$, $V = 1$, where $\epsilon_I = 10$ and $\epsilon_L = 10$ for panel G, $\epsilon_I = 10$ and $\epsilon_L = 5$ for panel H, and $\epsilon_I = 10$ and $\epsilon_L = 0$ for panel I.

Figure S2A,B uses a continuous trait model (6) with a transgenerational inducible defense. The specific model is

$$
\begin{aligned}
\frac{dx}{dt} &= \overbrace{x(r(\alpha)-kx)}^{\text{reproduction}} - \overbrace{\epsilon x}^{\text{non-predation mortality}} - \overbrace{\frac{b(\alpha)xy}{1+b(\alpha)hx}}^{\text{predation}} \\
\frac{dy}{dt} &= \overbrace{\frac{cb(\alpha)xy}{1+b(\alpha)hx}}^{\text{harvesting}} - \overbrace{my}^{\text{mortality}} \qquad \text{(S115)} \\
\frac{d\alpha}{dt} &= \underbrace{[\overbrace{\epsilon_I P_i(\alpha_1-\alpha)}^{\text{induction}} - \overbrace{\epsilon_L P_L(\alpha-\alpha_L)}^{\text{loss of induction}}](f+\epsilon)}_{\text{phenotypic plasticity}} + \underbrace{-Vg_{\alpha_i}}_{\text{phenotypic sorting}}
\end{aligned}
$$

where all functions are defined as above. Here, $\epsilon$ is used to control the turnover rate of the prey such that higher values of $\epsilon$ mean that prey have higher reproductive output and experience higher non-predation mortality; these effects cancel out in the prey equation. This continuous trait model can be derived from a dimorphic model where each prey type has growth rate $x_i f_i = x_i[r(\alpha_i) + \epsilon - kx]$, non-predation mortality rate $\epsilon x_i$, and predation rate $x_i b(\alpha_i)y/[1 + b(\alpha_1)hx_1 + b(\alpha_2)hx_2]$. Without loss of generality, the simulations use the parameter values $\alpha_1 = 0$ and $\alpha_2 = 1$.

**Figure S2AB:** $P_i(y) = [1 + (y/q)^\theta]$, $r_0 = 1$, $r_1 = -0.9$, $k = 1$, $b_0 = 0.3$, $b_1 = -0.25$, $h = 1$, $c = 20$, $m = 1$, $q = 0.1$, $\theta = 2$, and $V = 1$, where $\epsilon = 1$ for panel A and $\epsilon = 0$ for panel B.

**Figure S2C:** This panel shows a simulation of model (S113) where $\epsilon_I = \epsilon_L = 1.0145$ and all other parameter values are identical to Figure S2B. The values of $\epsilon_I$ and $\epsilon_L$ are chosen such that they are equal to value of $f$ at the equilibrium point in Figure S2AB. See Section S1.5.8 for details.

### References

Abrams, P. A., Y. Harada, and H. Matsuda. 1993. On the relationship between quantitative genetic and ESS models. Evolution 47:982–985.

Beddington, J. R. 1975. Mutual interference between parasites or predators and its effect on searching efficiency. Journal of Animal Ecology 44:331–340.

Cortez, M. H. 2011. Comparing the qualitatively different effects rapidly evolving and rapidly induced defences have on predator-prey interactions. Ecology Letters 14:202–209.

———. 2016. How the magnitude of prey genetic variation alters predator-prey eco-evolutionary dynamics. American Naturalist 188:329–341.

Crowley, P. H., and E. K. Martin. 1989. Functional responses and interference within and between year classes of a dragonfly population. Journal of the North American Benthological Society 8:211–221.

DeAngelis, D. L., R. A. Goldstein, and R. V. O'Neill. 1975. A model for trophic interaction. Ecology 56:881–892.

Yamamichi, M., T. Klauschies, B. E. Miner, and E. van Velzen. 2019. Modelling inducible defences in predator–prey interactions: assumptions and dynamical consequences of three distinct approaches. Ecology letters 22:390–404.
